# Myosin1G promotes Nodal signaling to control Zebrafish Left-Right asymmetry

**DOI:** 10.1101/2023.09.15.557992

**Authors:** Akshai Janardhana Kurup, Florian Bailet, Maximilian Fürthauer

**Affiliations:** Université Côte d’Azur, CNRS, Inserm, iBV, France

## Abstract

Myosin1D (Myo1D) has recently emerged as a conserved regulator of animal LR asymmetry that governs the morphogenesis of the central LR Organizer (LRO). In addition to Myo1D, the zebrafish genome encodes the closely related Myo1G. While Myo1G also controls LR asymmetry, we show that it does so through an entirely different mechanism. Myo1G promotes the Nodal-mediated transfer of laterality information from the LRO to target tissues. At the cellular level, Myo1G is associated with endosomes positive for the TGFβ signaling adapter SARA. *myo1g* mutants have fewer SARA-positive Activin receptor endosomes and a reduced responsiveness to Nodal ligands that results in a delay of left-sided Nodal propagation and tissue-specific laterality defects in organs that are most distant from the LRO. Beyond LR asymmetry, Myo1G promotes signaling by different Nodal ligands in other biological contexts. Our findings therefore identify Myo1G as a novel positive regulator of the Nodal signaling pathway.

## INTRODUCTION

Left-Right (LR) asymmetries in the positioning and shape of different tissues are found in both protostome and deuterostome lineages and critically required for human organ function^1^. In spite of the importance of LR asymmetry, our understanding of the mechanisms that govern this third body axis remains fragmentary. A particularly striking feature of LR asymmetry is the fact that an evolutionary conserved mechanism of symmetry breaking has long remained elusive. Although Nodal proteins of the Transforming Growth Factor β superfamily have long been known to control LR asymmetry in all deuterostome and some protostome species^2,3^, it is only recently that the unconventional type 1 Myosin Myosin1D (Myo1D) has emerged as a potentially universal regulator of animal LR asymmetry^4-7^. Here, we identify a close orthologue of Myo1D, Myosin1G (Myo1G) as a novel positive regulator of the Nodal signaling pathway.

Seminal studies in the mouse revealed the existence of a central LR Organizer (LRO) in which the Planar Cell Polarity (PCP)-dependent orientation of motile cilia promotes the generation of a directional symmetry-breaking fluid flow^8-10^. Symmetry-breaking cilia-driven fluid flows are also present in other species including fish and frogs^11,12^. Already within the vertebrate phylum, the LROs of birds and reptiles do however lack motile cilia and rely - at least in chick - on lateralized cell flows to trigger symmetry breaking^13,14^. Additional mechanisms implicated in LR asymmetry include ion flows^15^ and Actin-dependent chiral cell remodeling^16,17^. While an increasing number of studies indicate that Actin- and PCP-dependent pathways lie at the core of a symmetry-breaking toolbox^4,5,18-22^, our understanding of the evolutionary conservation of the mechanisms controlling LR asymmetry remains fragmentary.

In vertebrates, Nodal ligands convey laterality information from the central LRO to different target tissues^1,2^. Nodal ligands propagate on the left side of the embryo by inducing their own expression, allowing them to propagate from the posteriorly located LRO to more anterior target tissues ^23^. In species with a LRO bearing motile cilia, Nodal is expressed initially in a bilaterally symmetric fashion at the LRO, together with the TGFβ signaling antagonist Dand5^24^. Upon establishment of a ciliary LRO flow, *dand5* transcripts are degraded on the left side of the LRO^25-27^, allowing Nodal to travel to the left lateral plate mesoderm and propagate by autoinduction.

Nodal ligands induce cellular responses through ligand/receptor complexes that comprise TGFβ type I and II receptors and the co-receptor Cripto/Oep^28^. Nodal ligand binding causes type II receptors to phosphorylate and activate their type I counterpart. A population of endosomes positive for the TGFβ signaling adapter Smad Anchor for Receptor Activation (SARA) promotes signal transduction by allowing Activin/Nodal receptors to recruit their transcriptional downstream mediators SMAD2 & 3^29^. Upon phosphorylation by activated type I receptors, SMAD2 & 3 associate with SMAD4 to enter the nucleus and activate target genes^23^. As Nodal ligands are highly potent, a tight regulation of Nodal signaling is essential not only for embryonic development but also to avoid tumorigenesis^23,30^. Lefty proteins act as feed-back inhibitors of Nodal signaling that prevent the formation of productive ligand/receptor complexes^23,31^. In LR asymmetry, Lefty expression at the embryonic midline is important to form a midline barrier that prevents the spreading of left-sided Nodal ligands to the contralateral side^32,33^.

The requirement of Nodal ligands for LR asymmetry is however not universally conserved^1^ and a number of protostomian species, including the fruitfly *Drosophila*, altogether lack *nodal* homologues. Studies in *Drosophila* identified Myo1D as a master regulator of LR asymmetry^34,35^. In contrast to the central LRO of vertebrate organisms that governs LR asymmetry of all lateralized organs, *Drosophila myo1d* acts in a local, tissue-autonomous fashion to control genital and visceral laterality^18,35^. Of particular interest, studies in frogs, fish and humans showed that Myo1D is also required for vertebrate LR asymmetry^4-7^.

Zebrafish Myo1D is required for the establishment of a functional symmetry-breaking ciliary LRO flow^5^. In addition to *myo1d*, the fish genome harbors the closely related gene *myosin1g (myo1g)*. Although *myo1g* mutations impair laterality and enhance the defects of *myo1d* mutants, we show that Myo1G acts independently of the LRO flow, through an entirely different mechanism. We provide evidence that Myo1G represents a novel positive regulator of the Nodal signaling pathway whose function is essential for the Nodal-mediated transfer of laterality information.

## RESULTS

### Myosin1G mutants present tissue-specific Left-Right asymmetry defects

Myo1D controls cilia orientation in the LRO to promote the generation of a symmetry-breaking LRO flow^4-6^. The closely related protein Myo1G (79% amino acid similarity) is also required for zebrafish LR asymmetry but has no detectable effect on the LRO flow^5^, suggesting that different type I Myosins regulate LR asymmetry through distinct mechanisms. To address this issue, we performed a detailed characterization of *myo1g* single and *myo1d; myo1g* double mutants.

*myo1g* single mutants present defects in the leftward jogging of cardiac progenitors, the penetrance of which is further enhanced in *myo1d; myo1g* double mutants (Fig. 1a). To study the effect of *myo1g* on brain laterality, we analyzed the expression of the Nodal ligand *cyclops/nodal related 2 (cyc/ndr2)*, its feed-back antagonist *lefty1 (lft1)* and its transcriptional effector *pitx2* which display predominantly left-sided expression in the dorsal epithalamus of wild-type embryos^33,36,37^. In contrast to the mild defects observed in *myo1d* mutants (Fig. 1b, Supplementary Fig. 1a, b), *myo1g* single mutants displayed a significantly higher proportion of brain laterality defects (Fig. 1b, Supplementary Fig. 1a, b). The penetrance of brain laterality defects in *myo1d; myo1g* double mutants is similar to the one observed in *myo1g* single mutants, confirming the predominant role of *myo1g* in brain laterality (Fig. 1b, Supplementary Fig. 1a).

**Figure 1:**
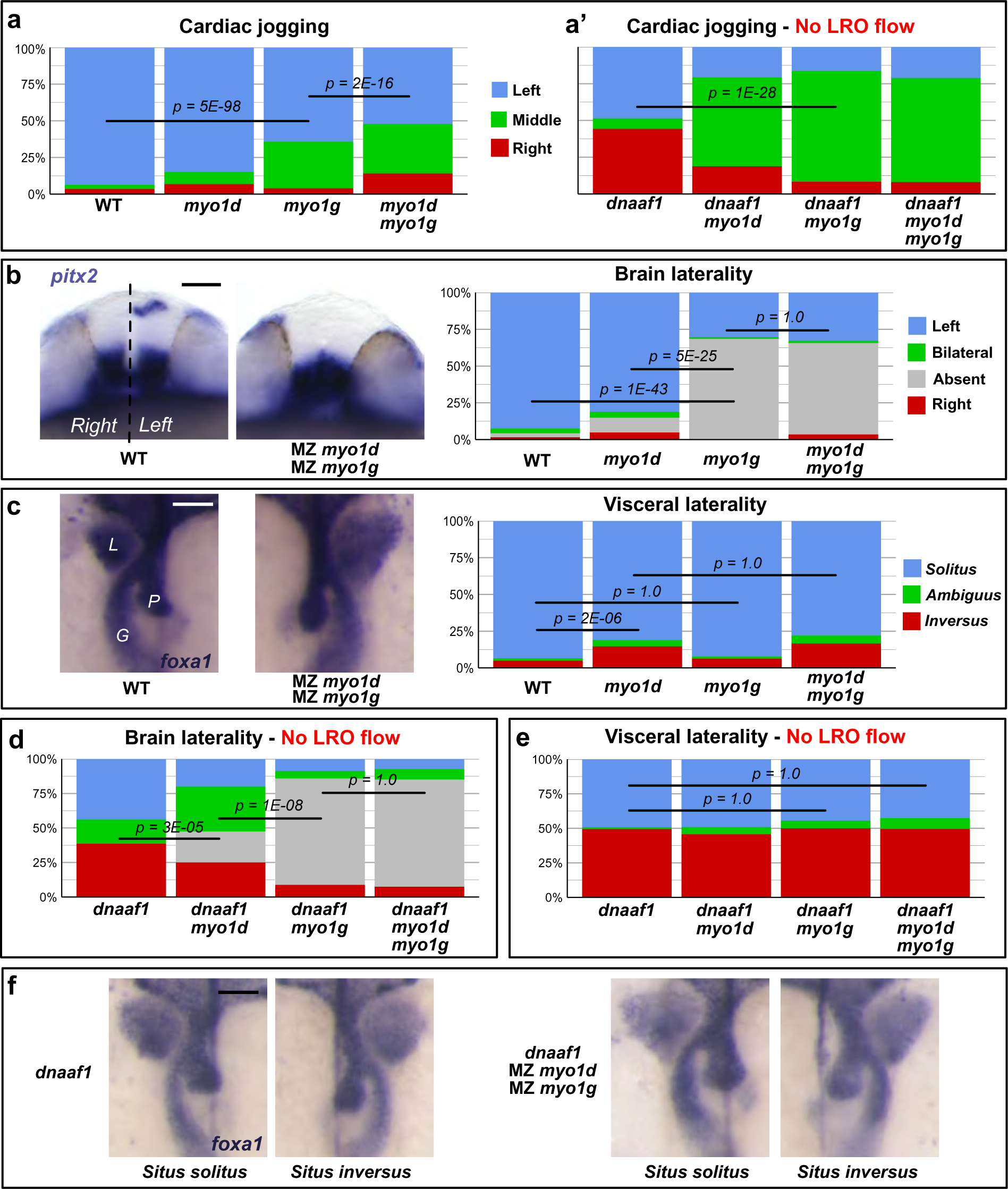
Myo1G regulates heart and brain LR asymmetry independent of the LRO flow. **a,a’** Quantification of cardiac jogging indicates that *myo1g* mutants present laterality defects that are enhanced in *myo1d;myo1g* double mutants (**a**). Concomitant inactivation of the LRO flow (through *dnaaf1* mutation) reveals that *myo1d/g* mutations enhance the cardiac jogging defects of flow-deficient animals (**a’**). **b** Brain asymmetry is impaired in *myo1g* and *myo1d;myo1g* mutants. **c** *myo1g* mutants do not show visceral LR defects. L: Liver, G: Gut, P: Pancreas. **d** *myo1d/g* inactivation enhances the brain laterality phenotypes of LRO flow-deficient *dnaaf1* mutants. **e,f** Visceral laterality phenotypes of dnaaf1 mutants are unaffected by *myo1d/g* inactivation. **b** Frontal views of *pitx2* expression at 30 somites, dorsal up. **c,f** Dorsal views of *foxa1* expression at 48h, anterior up. Scale bars: 50 µm.

In contrast to the effect of *myo1g* on brain laterality, analysis of liver, pancreas and gut laterality using the endodermal marker *foxa1* failed to reveal visceral LR asymmetry defects in *myo1g* single mutants (Fig. 1c). The observation that *myo1d; myo1g* double mutants present visceral laterality defects that are similar to *myo1d* single mutants (Fig. 1c) confirms that *myo1g* is dispensable for the establishment of visceral laterality.

Myo1D is required for LRO morphogenesis and the generation of a ciliary fluid flow^4-6^. Accordingly, *myo1d* mutants present defects at the level of all lateralized organs (Fig. 1a-c, Supplementary Fig. 1a, b). *myo1g* loss of function yields no discernable LRO flow defects^5^ and affects only in a subset of organs (Fig. 1a-c, Supplementary Fig. 1a, b), raising the question whether *myo1g* may control LR asymmetry through a flow-independent and potentially tissue-specific regulation of organ laterality, similar to the situation described for *Drosophila myo1d*^18,35^.

### Myosin1G controls Left-Right asymmetry independently of the Left-Right Organizer flow

To directly test if Myosin1 proteins exert LRO flow-independent functions in LR asymmetry, we investigated whether the LR asymmetry defects of animals that lack a LRO flow could be further modified by *myo1* gene inactivation. To this aim, we generated double and triple mutants to simultaneously inactivate *myo1d & g* and the essential regulator of ciliary motility *dnaaf1/lrrc50*^38^. Embryos that completely lack a LRO flow, as is the case for *dnaaf1* mutants, display a distinctive randomization of cardiac, brain and visceral laterality where LR asymmetry is properly established in roughly one half of the population (*situs solitus*) but inverted in the other (*situs inversus*, Fig. 1a’, d, e). Only a small fraction of the embryos that lack a LRO flow display an altogether loss of LR asymmetry (i.e. absence of cardiac jogging and brain laterality markers, visceral *situs ambiguus*, Fig. 1a’, d, e).

In contrast, animals that lack both a LRO flow and *myo1* function display a different phenotype, where the heart primordium fails to jog to either the left or the right side of the animal in most embryos (Fig. 1a’). *dnaaf1; myo1g* double mutants additionally present a lack of asymmetric *pitx2* expression in the dorsal epithalamus that contrasts with the randomization of lateralized gene expression observed in *dnaaf1* single mutants (Fig. 1b, d). In contrast to the effect observed at the levels of the heart and brain, the visceral phenotypes of *dnaaf1* mutant animals are unaffected by the loss of *myo1g* (Fig.1 e, f), confirming that *myo1g* is dispensable for visceral organ laterality.

These findings provide evidence for a novel, LRO flow-independent, function of Myosin1 proteins in LR asymmetry. The observations that i) *dnaaf1; myo1g* double mutants present a more pronounced loss of brain laterality then *dnaaf1; myo1d* mutants (Fig. 1d) and that ii) *dnaaf1; myo1d; myo1g* triple mutants are generally similar to *dnaaf1; myo1g* double mutants (Fig. 1a’, d, e) suggest that Myo1G exerts a predominant role in the flow-independent control of LR asymmetry.

### Myosin1G is required for Nodal pathway gene expression

Myo1 proteins could act in different ways to ensure a tissue-specific control of embryonic LR asymmetry. First, zebrafish *myo1d & g* could act in an organ-intrinsic fashion to promote chiral morphogenesis as in *Drosophila*^18,35^. Second, Myo1 activity could be required for the Nodal-mediated propagation of laterality information from the central LRO to different target tissues.

Already prior to the first morphological manifestations of asymmetric cell movement, the heart primordium displays asymmetries in gene expression in response to Nodal signaling from the Left Lateral Plate Mesoderm (LLPM)^39,40^. Of particular interest, the cardiac primordia of *myo1g* single and *myo1d; myo1g* double mutants present a reduced left-sided expression of the Nodal downstream target and feedback inhibitor *lefty2* (*lft2*) that could reflect impaired Nodal signaling (Fig. 2a). Accordingly, the expression of *southpaw* (*spaw*), the zebrafish Nodal ligand responsible for left-sided Nodal signaling, is reduced and extends less anteriorly in the LLPM of *myo1g* single and *myo1d; myo1g* double mutants, while being affected to a lesser degree in *myo1d* single mutants (Fig. 2b, Supplementary Fig. 2a). In further accordance with impaired Nodal signaling, *myo1*-deficient animals display a reduced expression of the Nodal-targets *pitx2* and *elovl6* in the LLPM (Fig. 2c, Supplementary Fig. 2b, c) and a reduced extension of the Nodal feedback inhibitor *lft1* in the notochord that provides a molecular midline barrier for lateralized Nodal signaling (Fig. 2d).

**Figure 2:**
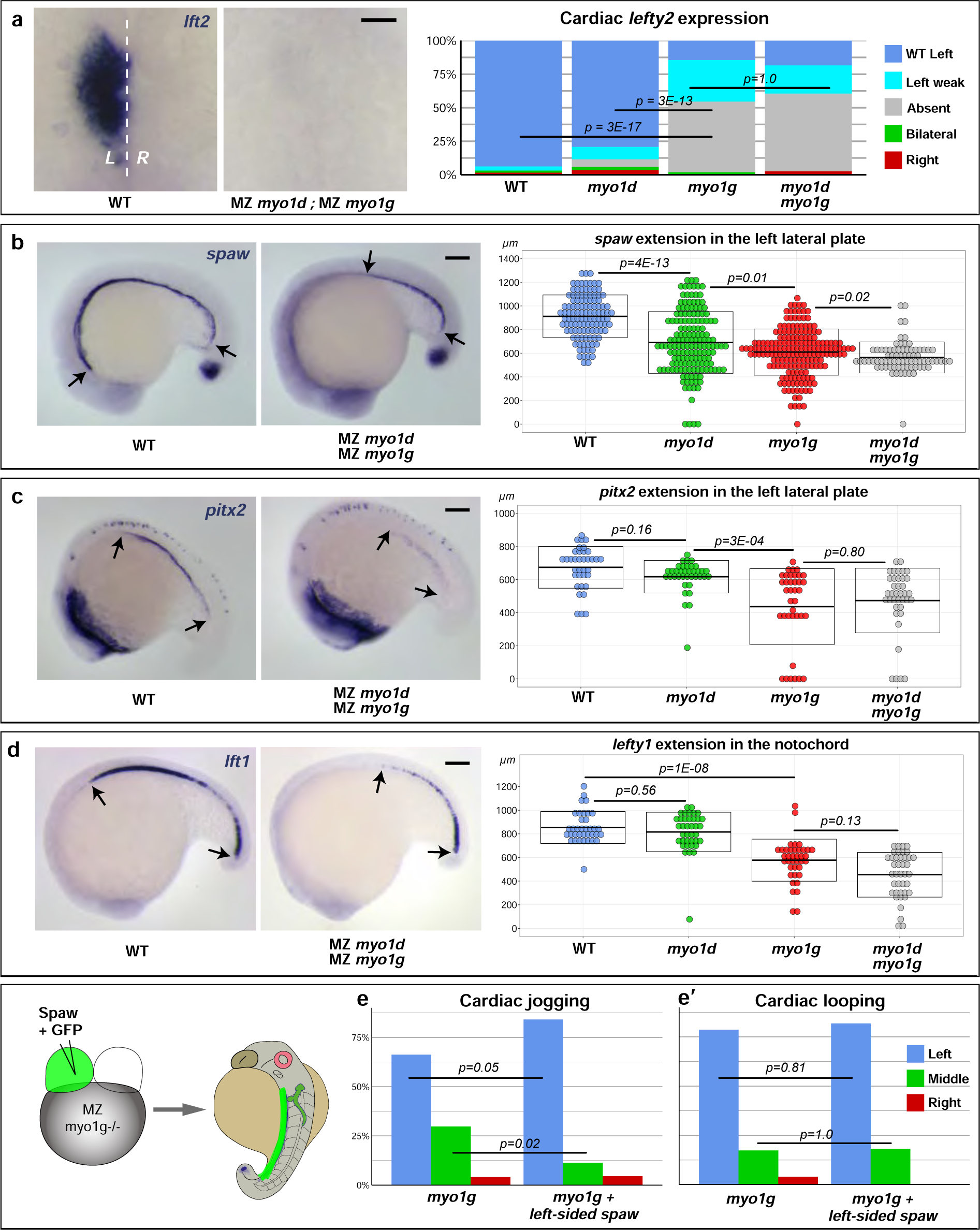
*myosin1* mutants display impaired Nodal signaling. **a** *myo1g* single and *myo1d; myo1g* double mutants fail to display asymmetric *lft2* expression in the cardiac primordium. **b-d** *myo1d/g* mutants display a reduced anterior propagation of the expression of the Nodal ligand *spaw* (**b**, see also Supplementary Fig. 2a), the Nodal effector *pitx2* (**b**, see also Supplementary Fig. 2b) and the Nodal feed-back inhibitor *lft1* (**d**). **e,e’** Left-sided expression of Spaw rescues the cardiac jogging (**e**) but not the cardiac looping (**e’**) defects of *myo1g* mutants. a Dorsal views at 22 somites, anterior up. **b-d** lateral views at 18 somites, anterior left, dorsal up. Scale bars: **a** 50 µm, **b,c** 100 µm.

### Unilateral Nodal expression restores cardiac laterality in *myosin1g* mutant animals

If the LR asymmetry defects of *myo1g* mutant animals are due to impaired Nodal signaling, restoring left-sided Nodal signaling should allow to rescue cardiac laterality. To test this hypothesis, Spaw and GFP RNAs were co-injected into a single blastomere at the two cell stage. By the end of gastrulation, the GFP tracer allowed to select animals in which the progeny of the injected blastomere was restricted to either the left or the right side of the embryo. In accordance with a potential requirement for *myo1g* in Nodal signaling, left-sided Spaw expression allowed to increase the percentage of *myo1g* mutants that present a proper leftward cardiac jogging (Fig. 2e). In contrast, the chirality of cardiac looping, a process that subsequently generates the atrial and ventricular chambers and occurs largely independently of Nodal signaling^41^ was not restored by left-sided Spaw expression (Fig. 2e’). Similarly, *myo1g* mutants in which Spaw-injected cells ended up on the right side of the embryo failed to display a restoration of embryonic laterality (Supplementary Fig. 2d, d’).

### The Left-Right organizer flow and *myosin1* genes control Nodal propagation

Through its ability to promote the unilateral degradation of transcripts encoding the Nodal signaling antagonist Dand5, the LRO flow enables the left-sided propagation of *nodal* expression^25-27^. Our observation that Myo1G and (to a lesser degree) Myo1D act to promote propagation of *spaw* expression (Fig. 2, Supplementary Fig. 2) raises the question whether the enhanced laterality defects of embryos that lack both a LRO flow and *myo1* gene function (Fig. 1) could be due to cumulative effects on *nodal* gene expression? To address this issue, we performed a comparative quantitative analysis of *spaw* expression in the Lateral Plate Mesoderm (LPM) of embryos that lack a LRO flow (due to *dnaaf1* inactivation) as well as *myo1d & g* activities.

In wild-type control embryos, *spaw* extends anteriorly up to the level of the heart and brain primordia in the left LPM, while its expression is either entirely absent or only restricted to the posterior-most LPM on the right side of the embryo (Fig. 3a). Morpholino-mediated knock-down of *dnaaf1* (Fig. 3a) or its genetic inactivation (Fig. 3b) cause a reduction in the left-sided extension of *spaw* which is likely due to a failure to downregulate *dand5* on the left side of the LRO. Additionally, *dnaaf1*-deficient animals present a roughly symmetric expression of *spaw* in the right LPM (Fig. 3a, b). Simultaneous inactivations of *myo1d/g and dnaaf1* cause a reduction of the anterior extension of *spaw* expression on both the left and the ride side the animal (Fig. 3a, b), demonstrating thereby that Myosin1 protein exert a flow-independent control of *nodal* ligand expression. Similar results were obtained using *dnaaf1* morphants or mutants, although quantitatively stronger effects are observed upon use of stable genetic mutants compared to transient morpholino knock-down. In accordance with our morphological analysis of embryonic laterality that suggested a predominant role of Myo1G in the LRO flow-independent control of LR asymmetry (Fig. 1), the inactivation of *myo1g* has a stronger effect on *spaw* expression in the LPM of *dnaaf1*-depleted animals then the loss of function of *myo1d* (Fig. 3a, b).

**Figure 3:**
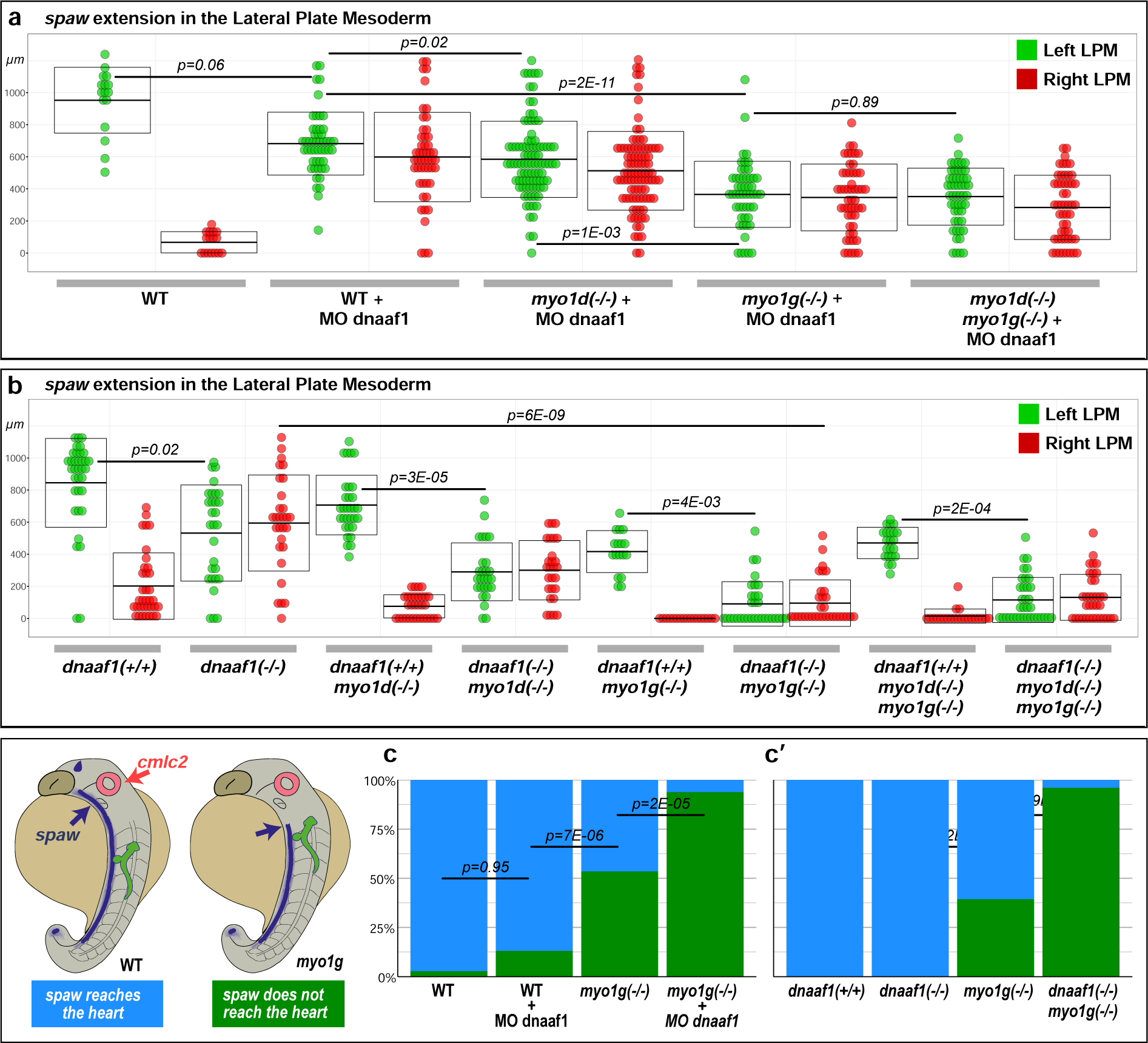
Myosin1 proteins regulate *spaw* expression independently of the LRO flow. **a,b** Quantification of spaw extension in the Left (green dots) and Right (red dots) LPM of 18 somites stage LRO flow-deficient dnaaf1 morphant (**a**) or *dnaaf1* mutant (**b**) embryos. *myo1d/g* loss of function causes a significant reduction of the antero-posterior extension of *spaw* expression in both the Left and the Right LPM. **c,c’** Double *in situ* hybridization for *spaw* and the cardiac marker *cmlc2* (see also Supplementary Fig. 3) reveals that *spaw* expression reaches the cardiac primordium in most WT control and LRO flow-deficient dnaaf1 morphant (**c**) or *dnaaf1* mutant (**c’**) embryos, but fails to do so upon inactivation of *myo1g*.

### Nodal expression fails to reach the cardiac primordium in *myosin1g* mutants

Spaw-mediated Nodal signaling is required to transmit laterality information from the LRO to target tissues. The zebrafish LRO, Kupffer’s Vesicle^11^ is located at the posterior tip of the notochord. Among the different tissues undergoing chiral morphogenesis, the visceral organ primordia are closest to the LRO while heart and brain primordia are located more anteriorly at increasing distances. As our experiments show that *myo1g* mutants present no defects in visceral LR asymmetry but increasingly severe phenotypes in the more anterior heart and brain (Fig. 1a-c), we wondered whether the reduced extension of left-sided *spaw* expression (Fig. 2b) may result in a failure to reach more anteriorly located organ primordia. To test this hypothesis, we performed two colour *in situ* hybridization to simultaneously visualize *spaw* expression and the *cmlc2*-positive cardiac primordium.

Our analysis reveals that by the 22 somites stage, *spaw* expression has reached the cardiac primordium in most wild-type embryos (Fig. 3c, c’, Supplementary Fig. 3a, a’). Similarly, *spaw* extends up to the level of the heart primordium on either the left, the right or both sides of the embryo in most animals that are mutant or morphant for the LRO flow regulator *dnaaf1* (Fig. 3c, c’, Supplementary Fig. 3a, a’). In contrast, *spaw* expression fails to reach the cardiac primordium in a significant fraction of *myo1g* mutants, providing thereby a potential explanation for their cardiac jogging defects (Fig. 3c, c’, Supplementary Fig. 3a, a’). Compound inactivations of *dnnaf1* and *myo1g* result in near complete failure of *spaw* expression to reach the cardiac primordium (Fig. 3c, c’, Supplementary Fig. 3a, a’), in accordance with the predominant lack of cardiac jogging that is observed in these animals (Fig. 1a’).

### *myosin1g* mutants display a temporal delay in *spaw* expression

To investigate the mechanism through which *myosin1* genes contribute to the LRO flow-independent regulation of Nodal signaling, we performed a time-course analysis of *spaw* expression during development. As *myo1d* contributes to both the regulation of the LRO flow^5^ and the flow-independent control of *nodal* expression (Fig. 3), we focused our analysis on *myo1g*, which plays a predominant role in the flow-independent control of Nodal signaling (Fig. 1, Fig. 3).

In wild-type embryos, *spaw* expression is initiated bilaterally in the cells that surround the LRO by the 6 somites stage (Fig. 4a). As development proceeds, *spaw* LRO levels increase until at around the 12 somites stage expression also becomes detectable in the left LPM where the ligand then propagates through auto-induction to reach more anterior target tissues (Fig. 4a). Analysis of *spaw* expression in *myo1g* mutants revealed a reduction in the initial induction of *spaw* expression at the level of the LRO and the subsequent propagation to the LPM (Fig. 4a). In contrast to the loss of *myo1g* function, a lack of LRO-flow upon depletion of *dnaaf1* is without effect on the initial induction of *spaw* expression at the LRO (Supplementary Fig. 4a).

**Figure 4:**
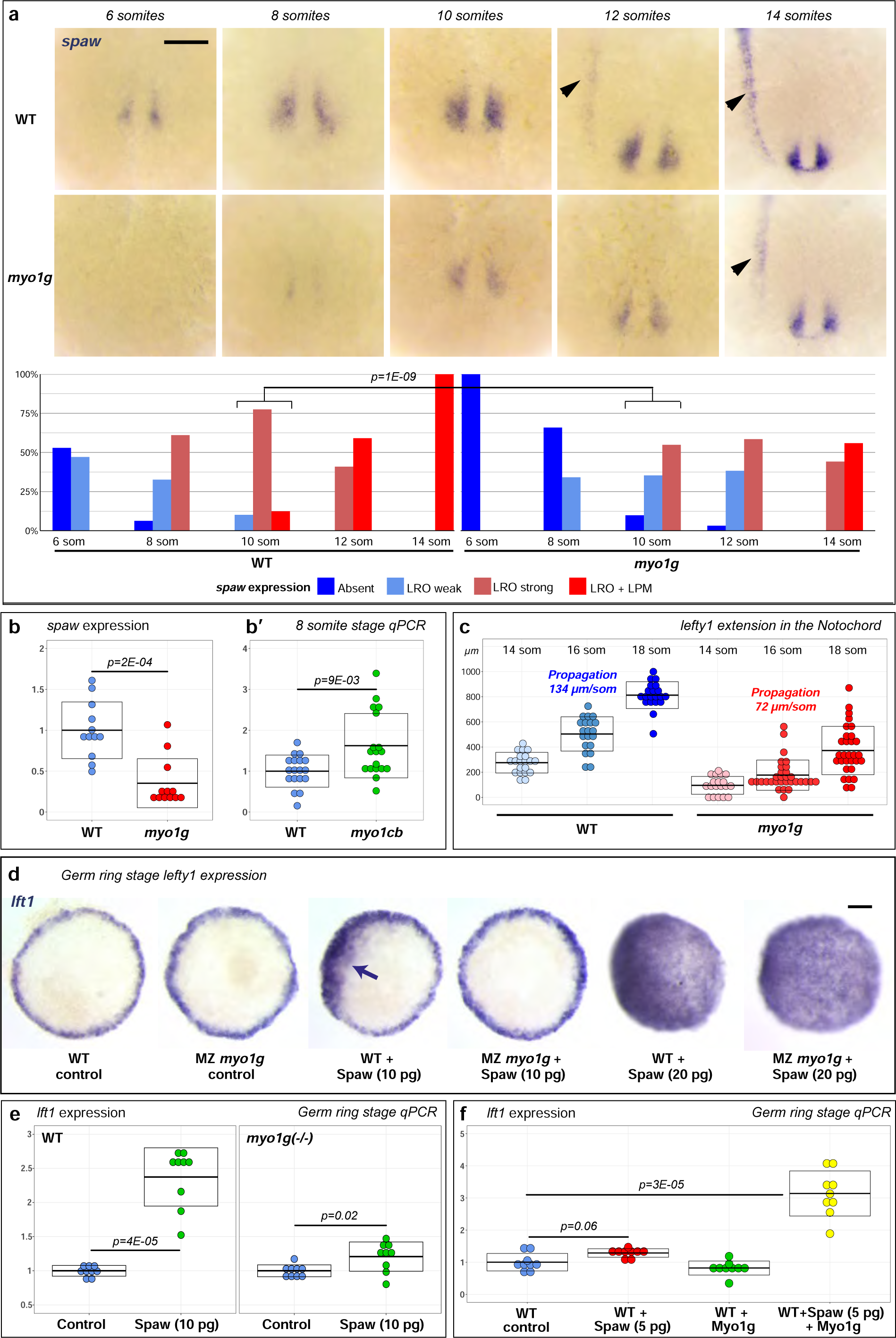
Myo1G promotes Spaw signaling. **a** Time course analysis of *spaw* expression indicates that initiation of *spaw* expression at the LRO and propagation to the Left LPM (black arrowhead) are delayed in *myo1g* mutants. **b,b’** qPCR analysis of spaw expression confirms that *spaw* expression is significantly reduced in *myo1g* mutants (**b**). Conversely, *spaw* expression increases in mutants for the *myo1d/g* antagonist *myo1cb* (**b’**, see also Supplementary Fig. 4b). **c** *myo1g* mutants present a reduced rate of anterior-ward propagation of *lft1* expression in the notochord (see also Supplementary Fig. 6a). **d,e** *myo1g* mutants display a weaker induction of the nodal target gene *lft1* in response to Spaw overexpression. While high amounts (20 pg) of Spaw RNA induce a similar *lft1* induction in *myo1g* mutants and wild-type siblings, *myo1g*-deficient embryos present a reduced response to moderate amounts (10 pg) of Spaw RNA (**d**, ectopic expression indicated by arrow, see Supplementary Fig. 6b for quantification). qPCR analysis confirms that equal amounts of Spaw RNA induce a reduced *lft1* induction response in *myo1g* mutants (**e**). f Conversely, the overexpression of Myo1G potentiates the capacity of low amounts (5 pg) of Spaw RNA to induce ectopic *lft1*. **a** vegetal views of the LRO, anterior up. **d** animal pole views. Scale bars: 100 μm.

The observation that *myo1g* mutants present a reduced *spaw* expression at the LRO was confirmed by quantitative qRT-PCR (Fig. 4b). Studies in *Drosophila* and zebrafish revealed that Myosin1C (Myo1C) proteins can act as Myo1D/G antagonists^5,42^. While our analysis failed to reveal any morphological LR asymmetry defects in maternal zygotic *myo1Cb* mutants, gene expression analysis uncovered a mild upregulation of *spaw* at the LRO (Fig. 4b’, Supplementary Fig. 4b), supporting the functional relationship between Myo1D/G agonists and their Myo1Cb antagonist.

### *myosin1g* is dispensable for Left-Right Organizer formation

The finding that zebrafish *myo1d* is required for LRO morphogenesis^5,6^ raises the question whether the loss of *myo1g* may similarly cause general defects in LRO morphogenesis that would ultimately result in reduced Nodal signaling at the LRO. Our analysis of different markers genes involved in LRO specification and function does however not support this hypothesis. Analysis of the endodermal markers *sox17* and *sox32* indicates that the specification and clustering of LRO precursor cells occurs normally in *myo1g* mutants (Supplementary Fig. 5a, b). In accordance with the fact that *myo1g* controls LR asymmetry independently of the LRO flow, *myo1g* loss of function has no effect on the expression of the ciliary motility genes *foxj1a, dnah9* and *odad1* (Supplementary Fig. 5c, d, e).

### Myosin1G promotes Nodal signaling

In mice and zebrafish, Nodal expression at the LRO is initially induced by Notch signaling^43^, and then further upregulated through the capacity of Nodal ligands to induce their own expression^44^. While our analysis of the Notch target genes *her4.1* and *her15.1* suggests that *myo1g* mutants present normal Notch signaling levels (Supplementary Fig. 5f, g), the reduced expression of *spaw* at the LRO (Fig. 4a) is similar to the one reported in *spaw* mutants^44^.

If *myo1g* mutants present a defect in Spaw autoinduction, this should result not only in an initial delay in the appearance of *spaw* expression in the LPM (Fig. 4a), but also in a delayed subsequent anterior propagation. Accordingly, time course analysis of *lft1* expression at the notochordal midline barrier reveals that *myo1g* mutants present a nearly two-fold reduction in the rate of Nodal target gene propagation (Fig. 4c, Supplementary Fig. 6a).

As Nodal ligands induce their own expression^23^, the reduced propagation of *spaw* expression in *myo1g* mutants could be indicative of a defect in Nodal signal transduction. To establish if Myo1G is important for Spaw signaling, we misexpressed Spaw in germ ring stage embryos that lack endogenous *spaw* expression and analyzed its capacity to upregulate the Nodal target gene *lft1*. While high doses (20 pg) of Spaw readily induce ectopic *lft1* expression in both WT and *myo1g* mutant embryos (Fig. 4d, Supplementary Fig.6b), *myo1g* mutants present a reduced response to moderate (10 pg) doses of Spaw RNA (Fig. 4d, Supplementary Fig. 6b). Analysis of *lft1* expression by quantitative RT-PCR reveals that while Spaw is still able to significantly induce *lft1* in *myo1g* mutants, the observed effect is weaker than in homozygous WT sibling controls (Fig. 4e, Cohen’s d effect size = 1.27 for *myo1g* mutants versus 4.48 for WT controls).

Taken together, our observations suggest that Myo1G, while not strictly required for Spaw signal transduction, is essential to promote full strength Nodal signaling. Injecting wild-type Myo1G RNA into *myo1g* mutants significantly rescues the capacity of Spaw to induce *lft1* expression, demonstrating the specificity of the observed effect (Supplementary Fig. 6c). To confirm that Myo1G promotes Spaw signaling, a lower amount of Spaw RNA (5 pg), that is on its own barely capable of inducing ectopic *lft1* expression, was co-injected with wild-type Myo1G RNA into WT animals. qRT-PCR analysis shows that Myo1G overexpression promotes the capacity of this subliminal amount of Spaw to induce *lft1* expression (Fig. 4f).

### *myosin1g* regulates Activin receptor trafficking

How does Myo1G promote Spaw signaling? Proteomic studies identified Myo1G on exosomes, suggesting that this factor may be implicated in exovesicular secretion^45^. To determine whether Myo1G is required for Spaw ligand secretion, we took advantage of a functional GFP-Spaw fusion construct that has previously been used to visualize Spaw secretion^46^. GFP-Spaw RNA injection into wild-type or *myo1g* mutant animals results in a similar labeling of the extracellular space (Fig. 5a, b), suggesting that Myo1G is unlikely to control Spaw ligand production and secretion.

**Figure 5:**
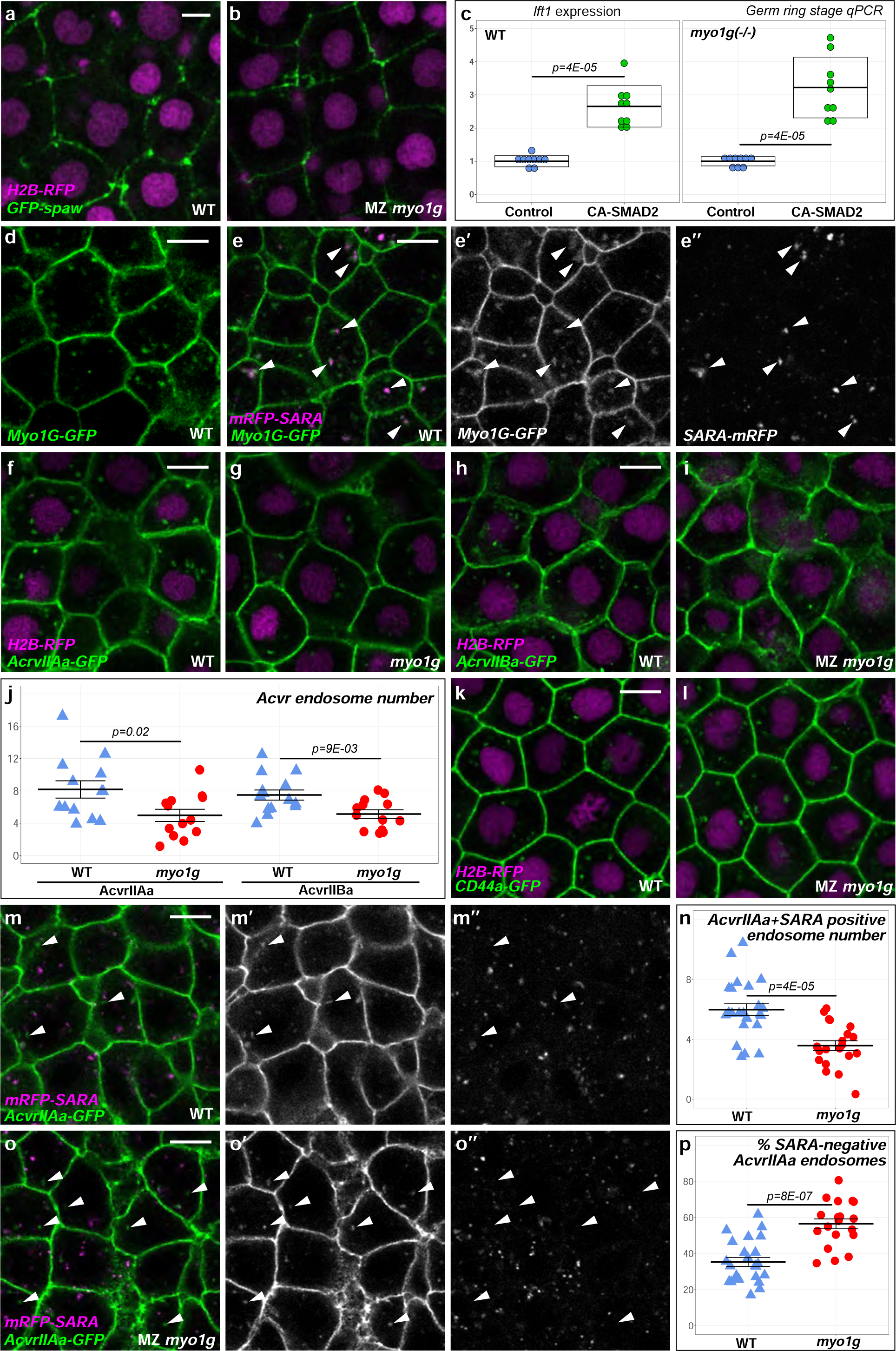
Myo1G regulates Nodal receptor trafficking. **a,b** Spaw-GFP localization is similar in WT (n=20) and *myo1g* mutants (n=17). H2B-RFP was injected as a tracer to ascertain that embryos that had received equal amounts of RNA. **c** A constitutively activated form of the Nodal signal transducer SMAD2 (CA-SMAD2) elicits similar responses in WT and *myo1g* mutants. **d** Myo1G-GFP is detected at the cell cortex and in intracellular compartments (n = 10). **e,e’,e’’** Myo1G-GFP is present on endosomes positive for the TGF® signaling adapter SARA (see also Supplementary Fig. 7a,b). **f-j** *myo1g* mutants present a reduced number of endosomes positive for the Nodal receptors Acvr2Aa-GFP (**f,g,j)** and Acvr2Ba-GFP (**h,i,j**). **k,l** *myo1g* mutants and WT siblings present a similar number of CD44a-positive endosomes (see also Supplementary Fig. 7c). **m-p** *myo1g* mutants present a lower number of endosomes that are positive for both AcvrIIAa and SARA (**n**). Conversely, *myo1g* loss of function causes an increase in the fraction of SARA-negative AcvrIIAa endosomes (**p**, arrowheads in **m,n**) despite having a similar number of SARA endosomes (Supplementary Fig. 7d). **a,b,d-i,k-m,o** animal pole views, germ ring stage. Data points in **j,n,p** represent the mean number of endosomes per cell for a particular embryo (see Supplementary material for complete statistical information). Scale bars: 10 μm.

While cytoplasmic Myo1C exerts important roles in membrane trafficking^47^, nuclear isoforms of mammalian Myo1C can regulate TGFβ-responsive gene expression^48^. To determine if Myo1G controls the SMAD-mediated transcriptional downstream response to Spaw signaling, we injected RNA encoding a Constitutively Activated variant of SMAD2 (CA-SMAD2) into wild-type and *myo1g* mutant animals and analyzed the effect on *lft1* target gene induction by qRT-PCR. In contrast to the reduced induction of *lft1* that is observed upon Spaw overexpression in *myo1g* mutants (Fig. 4e), CA-SMAD2 elicited a similar induction of *lft1* expression in *myo1g*-deficient animals (Fig. 5c, Cohen’s d effect size = 3.39 for *myo1g* mutants versus 3.62 for WT controls).

The pharmacological Myosin antagonist Pentachloropseudilin (PCIP) inhibits TGFβ signaling by regulating the membrane trafficking of TGFβ type II receptors^49^. In accordance with a potential function in the membrane trafficking of cell surface receptors, Myo1G-GFP localizes to both the cell cortex and to intracellular, potentially endosomal, compartments (Fig. 5d).

As endosomal compartments positive for the TGFβ signaling adapter SARA promote Nodal signal transduction^29^, we investigated if Myo1G-GFP positive intracellular compartments correspond to SARA endosomes. Strikingly, use of an established mRFP-SARA construct^50^ revealed that in 24/24 embryos, SARA positive compartments were always associated with Myo1G-GFP. Both standard laser scanning microscopy (Fig. 5e) and Airyscan super-resolution microscopy (Supplementary Fig. 7a, b) revealed that SARA-positive compartments are often part of larger, Myo1G-positive structures, suggesting that Myo1G may be important for the biology of TGFβ signaling endosomes.

SARA endosomes promote Nodal signaling by providing a subcellular platform that enables TGFβ receptors to activate downstream SMADs^29^. As Myosins have been linked to TGFβ type II receptor trafficking in other biological contexts^49^, we analyzed the effect of *myo1g* loss of function on the two Nodal type II receptors AcvrIIAa and AcvrIIBa. GFP-tagged versions of the two proteins indeed revealed a significant reduction of the number of Activin receptor-positive endosomes in *myo1g* mutants (Fig. 5f-j). In murine lymphocytes, Myo1G regulates the endocytic trafficking of the adhesion protein CD44 ^51^, a molecule that has, in other biological contexts, been shown to regulate TGFβ signaling by acting as Hyaluronan receptor ^52^. In contrast to the situation observed for AcvrIIAa & Ba, *myo1g* mutants and their wild-type siblings present similar numbers of CD44a-positive endosomes (Fig. 5k, l, Supplementary Fig. 7c), supporting the specificity of the observed Nodal receptor trafficking defects.

The observations that *myo1g* mutants present a reduced number of AcvrII endosomes (Fig. 5f-j) and that Myo1G is found on SARA-positive compartments (Fig. 5e, Supplementary Fig. 7a, b) raise the question whether Myo1G may be required for the formation of AcvrII/SARA-positive endosomes. Accordingly, *myo1g* mutants present both a reduction in the absolute number of SARA/AcvrIIAa-positive compartments (Fig. 5m-o) and a higher fraction of AcvrIIAa compartments that lack the signaling endosome marker SARA (Fig. 5m, n, p). In contrast, *myo1g* loss of function has no effect on the absolute number of SARA-positive endosomes (Supplementary Fig. 7d). Taken together, these findings suggest that Myo1G may ensure full strength Nodal signaling by promoting the formation of SARA/Nodal receptor-positive endosomes.

### Myosin1G promotes Southpaw-independent Nodal signaling

As *myo1g* mutants present a normal expression of LRO specification and differentiation markers (Supplementary Fig. 5), but reduced levels of the TGFβ superfamily ligand *spaw* (Fig. 4), we investigated the effect of *myo1g* loss of function on the LRO expression of other TGFβ signaling components.

To induce a biological response, Spaw heterodimerizes with the TGFβ superfamily member GDF3^53^. Our analysis shows that not only *spaw* itself, but also the expression of its partner *gdf3* is reduced in *myo1g* mutants (Fig. 6a). The Cerberbus/Dan family protein Dand5 antagonizes Spaw signaling at the LRO^24^. *myo1g* mutants not only present lower levels of *spaw*/*gdf3,* but also diminished *dand5* expression (Fig. 6b). Of particular interest, *myo1g* mutants display a reduced *dand5* expression already at the tail bud stage, 2 hours before the onset of *spaw* expression (Fig. 6b), suggesting that the function of Myo1G is not limited to Spaw signaling.

**Figure 6:**
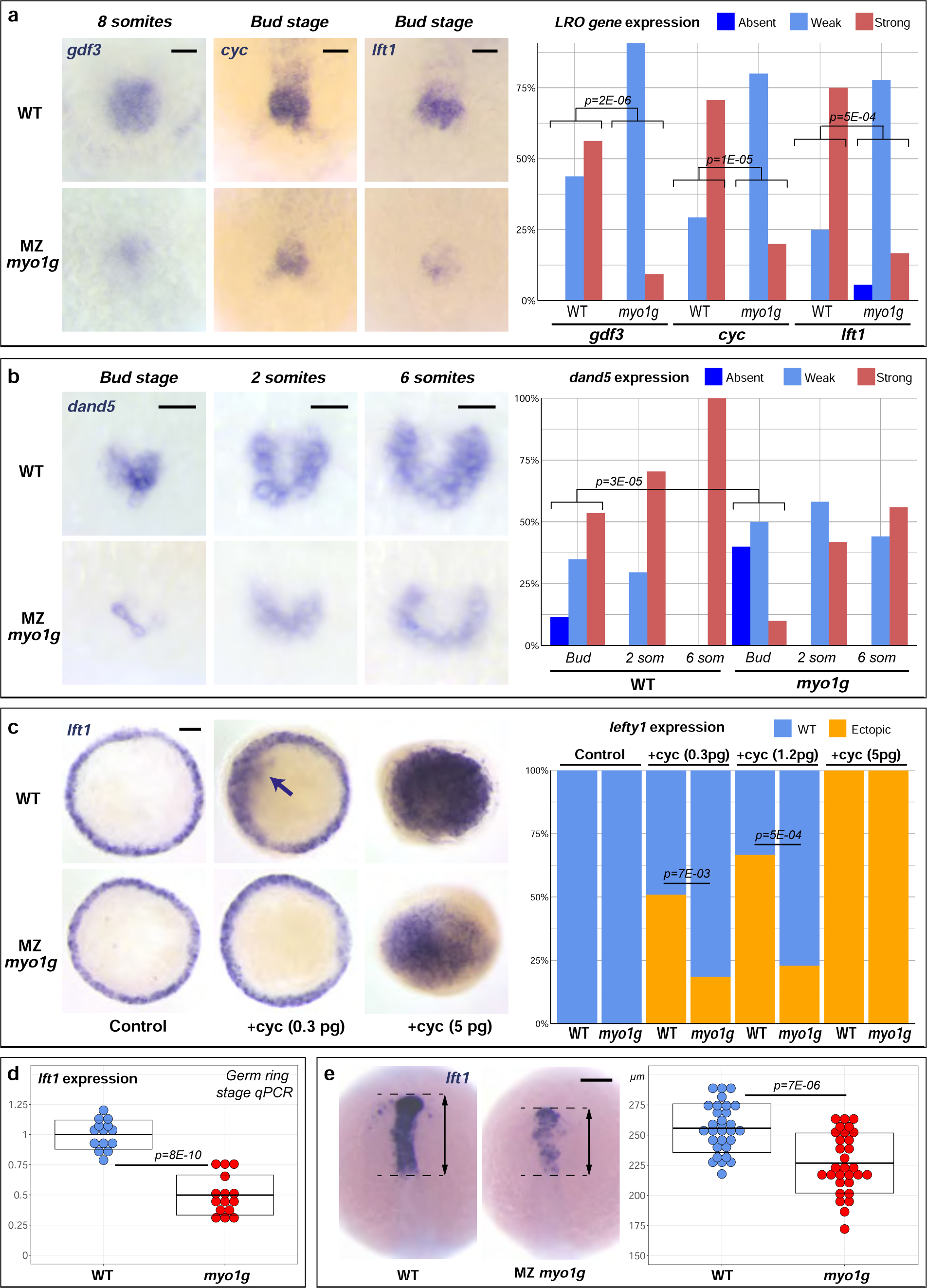
Myo1G promotes signaling by different Nodal ligands. **a** *myo1g* mutants present a reduced expression of the TGFβ ligands *gdf3* and *cyc* and the Nodal feed-back antagonist *lft1* in the LRO/tail bud region. **b** *myo1g* mutants display a reduced expression of the Nodal signaling antagonist *dand5*. **c** *myo1g* mutants display reduced *lft1* induction in response to ectopic Cyc expression (arrow indicates ectopic expression). **d** qPCR indicates that *myo1g* mutants present a significant decrease in the endogenous expression levels of the Nodal target gene *lft1*. **e** 8 somites stage *myo1g* mutants present a reduction in the antero-posterior extension of *lft1* expression in the anterior brain. **a,b** vegetal views of the LRO/tail bud region, anterior up. **c** animal pole views of germ ring stage embryos. **e** dorsal view of the anterior brain, anterior up. Scale bars: **a** 50 µm, **b,c,e** 100 µm.

Prior to *spaw* expression, the *nodal* ligand *cyc* and its target and feed-back inhibitor *lft1* are expressed in the LRO region (Fig. 6a)^33,36^. The observation that *myo1g* mutants present a reduction in the early LRO expression of *cyc* and *lft1* (Fig. 6a) suggests that, in addition to Spaw, Myo1G may potentiate signaling by other Nodal ligands. Accordingly, *myo1g* mutants display a reduced *lft1* induction in response to ectopic Cyc (Fig. 6c) or Squint/Nodal-related-3 (Sqt/Ndr3, Supplementary Fig. 8), the third zebrafish nodal homologue.

While the morphological LR asymmetry defects of *myo1g* mutants could be solely due to defective Spaw signaling, quantitative analysis of the Nodal downstream gene *lft1* provided evidence that Myo1G also promotes endogenous signaling by other Nodal ligands in different contexts. First, qRT-PCR revealed that the germ ring stage *lft1* expression which is induced by Cyc and Sqt is significantly reduced in *myo1g* mutants (Fig. 6d). Second, 8 somites stage *myo1g* mutants present a reduced expression of *lft1* in the anterior neurectoderm, which lacks *spaw* but expresses *cyc*^36,37^.

Taken together, our findings identify Myo1G as a novel positive regulator of the Nodal signaling pathway that is essential for LR asymmetry and potentiates responses elicited by different Nodal ligands in different biological contexts.

## DISCUSSION

A striking feature of LR asymmetry is that different species use seemingly different mechanisms for the determination of this third body axis^3^. Only recently has the unconventional type I Myosin Myo1D, which was initially identified as a regulator of *Drosophila* laterality^34,35^, been identified as an evolutionarily conserved regulator of animal LR asymmetry^4-7^. While studies in fish and frogs uncovered an essential role of Myo1D in LRO morphogenesis^4-6^, several observations suggested that additional functions of Myosin1 proteins in LR asymmetry remain still to be uncovered. First, previous studies had identified a function of Myo1D in the central LRO of fish and frogs, a biological structure that has no equivalent in *Drosophila*, where Myo1D ensures a local, organ-specific control of chiral morphogenesis. Second, in contrast to the unique *myo1d* gene present in flies, vertebrate genomes harbor not only *myo1d*, but also its close homologue *myo1g*. We present an in-depth analysis of the function of this second *myosin1* homologue in zebrafish and uncover a novel essential function of this gene in chiral morphogenesis that is different from the reported function of *myo1d* ^4-6^.

Myo1D controls the symmetry-breaking ciliary fluid flow in the central LRO^4-6^. Accordingly, *myo1d* loss of function causes defects in all lateralized organs^4-6^. In contrast, our findings show that *myo1g* is required for the chiral morphogenesis of the heart and brain, but dispensable for visceral laterality (Fig. 1, Supplementary Fig. 1). To specifically determine whether Myo1G exerts an LRO flow-independent function, we inactivated *myo1g* in the context of animals that lack the ciliary motility gene *dnaaf1* and therefore have no LRO flow. Lack of a LRO flow in *dnaaf1* mutants causes a distinctive randomization of LR asymmetry, in which the heart, brain and viscera develop either normally (*situs solitus*) or as their mirror image (*situs inversus*). In contrast, a different phenotype is observed in *dnaaf1 myo1g* double (or *dnaaf1 myo1d myo1g* triple mutants) in which cardiac and brain laterality are altogether lost (Fig. 1). These findings establish an essential role of Myo1G in the flow-independent, tissue-specific control of LR asymmetry.

How does Myo1G exert this tissue-specific control? Myo1G could be involved in the local, organ-specific execution of chiral morphogenesis, like *Drosophila myo1d*^18,35^. Alternatively, Myo1G could be involved in the transmission of laterality information from the central LRO to different target tissues. While our experiments do not allow to rule out the first possibility, *myo1g* mutants present a reduced propagation of the Nodal ligand *spaw* and a reduction in the expression of different Nodal target genes (Fig. 2, Supplementary Fig.2). *myo1g* mutant laterality defects can moreover be rescued through unilateral Spaw expression (Fig. 2e).

The observation that *myo1g* mutants can be rescued through Nodal overexpression shows that Nodal signaling is reduced but not abolished in these animals. Quantitative analysis of Nodal target indeed reveals a reduced responsiveness to Nodal ligands in mutant embryos (Fig. 4). In accordance with a disruption of Nodal signal transduction, Myo1G is found on endosomes that are positive for the TGFβ signaling adapter SARA and *myo1g* mutants present a reduced number of Nodal-receptor positive SARA endosomes (Fig. 5).

The observation that Myo1G is found on SARA endosomes and regulates Nodal receptor trafficking raises the question whether, beyond LR asymmetry, Myo1G may promote Nodal signaling in other biological contexts. In accordance with this hypothesis, our experiments show that *myo1g* mutants present a reduced responsiveness to the Nodal ligands Cyc and Sqt (Fig. 6c, Supplementary Fig. 8) and a reduction of endogenous Nodal target gene expression levels in domains that are independent from the *spaw*, the *nodal* gene entirely dedicated to LR asymmetry (Germ ring stage blastoderm margin, 8 somites stage forebrain, Fig. 6d, e).

Taken together, our findings identify Myo1G as a general positive regulator of Nodal signaling whose function is specifically required for LR asymmetry. Our work establishes for the first time a link between unconventional type 1 Myosins that are emerging as major regulators of animal laterality, and Nodal signalling which has long been known to be the key pathway regulating vertebrate LR asymmetry.

## METHODS

### Zebrafish strains and embryo maintenance

Embryos were raised in 0.3X Danieau medium (17.4mM NaCl, 0.21mM KCl, 0.12mM MgSO4, 0.18mM Ca (NO3)2, 1.5mM Hepes, pH 7.6) at 28.5 °C., and staged according to standard criteria^54^. If necessary, 1-phenyl-2-thiourea (Sigma) was added at 30 mg/l to prevent embryonic pigmentation.

*myo1d*/*g* inactivations were performed using the previously reported *myo1d^tj16b^*, *myo1d^tj16c^* and *myo1g^tj18b^* alleles^5^. All presented data were obtained using Maternal Zygotic (MZ) single or double mutants. Allele specific PCR was used to identify the WT *myo1d* allele (forward primer 5′-AGAGTGGAGCTGGAAAAACAGA-3′, reverse primer 5′-CCCATCCCTCGTGTGAAACTAAATCAC-3′, 339 bp amplicon) as well as the mutant alleles tj16b (forward primer 5′-TGGAGCTGGAAAAAGGCTCGT-3′, reverse primer 5′-CCATCACTGCAGCAGAAATGAGAG-3′, 133 bp amplicon) and tj16c (forward primer 5′-GTGGAGCTGGAAAAAGGCTATAC-3′, reverse primer 5′-CCATCACTGCAGCAGAAATGAGAG-3′, 145 bp amplicon). The allele-specific reverse primers 5′-TCTCATACAGTTCTCTTCCCCTAG-3′ (tj18b, 115 bp amplicon) and 5′-CTCATACAGTTCTCTTCCCCTGTAG-3′ (WT, 120 bp amplicon) were used with the generic forward primer 5′-GAGAAGAGTCGTATCTACACCTTC-3′ to genotype *myo1g* mutant fish.

*myo1Cb* inactivation was performed using the *myo1Cb^sa16637^* allele from the Zebrafish Mutation Project (http://www.sanger.ac.uk/resources/zebrafish/zmp/) obtained from the Zebrafish International Resource Center. The *sa16637* allele introduces a premature stop codon at the 228^th^ amino acid position. A generic forward primer 5’-GTCACATCCTGAACTACCTGCTAG-3’ was used along with a mutant specific reverse primer 5’-TATTACCAGTATCTGGTCAAG-3’ (164 bp amplicon) to identify the mutant allele. The WT allele was identified using the generic forward primer with the WT specific reverse primer 5’-CAGTACCAGTATCTGGTCAAG-3’ (164 bp amplicon).

*dnaaf1* was inactivated using the *dnaaf1^tm317b^*mutant allele^38^. The forward primer 5’-GCAAGCTTTGCACGCTTAATGTCTC-3’ and reverse primer 5’ - AACACTGGAGAATGTTTGTGAC - 3’ were used to amplify the tm317b mutant allele (199 bp amplicon). The *dnaaf1* WT allele was identified using the forward primer 5’-GCAAGCTTTGCACGCTTAATGTCTC-3’ and reverse primer 5’- CACACTGGAGAATGTTTGTGAC-3’ (199 bp amplicon). Beyond 24 hrs of development, *dnaaf1* mutants can be identified through the oval phenotype that is diagnostic for ciliary mutations.

### Plasmid generation

The *myo1g* ORF was amplified from mixed stage pool of cDNAs using primers 5’-**GATCCCATCGATTCGA**TGGCGGAGCTGGAGGGCTTG-3’ and 5’-**AGGCTCGAGAGGCCTT**ACTGGGGCAGGAGTAAGG-3’ and cloned into the pCS2+ vector using Gibson assembly mix (NEB). Bold letters in the primer sequences indicate Gibson overhangs that are also present in the pCS2+ sequence. For generating the *myo1g*-GFP construct, the *myo1g* ORF was amplified from the *myo1g*-pCS2 construct using the primer pair 5’-**GCAGGATCCCATCGATTCGACAGTAAAC**ATGGCGGAGCTGGAGGGCTTG-3’ and 5’-**ACCATGGACCCTCCGCTGGTGC**CCTGGGGCAGGAGTAAGGTAAATC-3’, and was ligated onto pCS2-GFP.

*CD44a* was amplified using the primers 5’-**ATCCCATCGATTCGACAGTAAAC**ATGTGGACTTTGTTATTTGTAGTGTT-3’ and **5’-ACCATGGACCCTCCGCTGGTGCCC**ATTAAATATTCTTTTTCGTGTTCA-3’ and ligated into pCS2-eGFP. For *acvr2aa*, the following primer pairs were used 5’-**GGATCCCATCGATTCGACAGTAAAC**ATGGGACCTGCAACAAAGCTGGC-3’ and 5’-**ACCATGGACCCTCCGCTGGTGCC**TAGACTAGACTCCTTTGGGGGATA-3’, and for *acvr2ba*, the forward primer 5’-**GATCCCATCGATTCGACAGTAAAC**ATGTTCGCTTCTCTGCTCACTTTGG-3’ was used with the reverse primer 5’-**ACCATGGACCCTCCGCTGGTGCC**GATGCTGGACTCTTTGGGCGG-3’ to amplify the ORFs, which were ligated into pCS2-eGFP. The *her4.1*-pBSK construct used to generate *in situ* probe was cloned using the primer pair 5’-**GTCGACGGTATCGATAAGC**CACACAGCAATGACTCCTAC-3’ and 5’-**CTAGAACTAGTGGATCCCCC**TTAAGTCTACCAGGGTCTCC-3’. *her15.1* was amplified using the forward primer 5’-**GTCGACGGTATCGATAAGC**GCTCAGAGAAACAGCATCTCTCC-3’ and reverse primer 5’-**CTAGAACTAGTGGATCCCCC**CTCCACAGGAGTTCAACATTGAC-3’ and cloned into pBSK. A Squint in situ probe was amplified using the primers 5’-**CGAGGTCGACGGTATCGATAAGC**ACATGTTTTCCTGCGGGC-3’ and 5’-**CTCTAGAACTAGTGGATCCCCC**GTTTGAAGAATCAGTGGCAGC-3’ and cloned into pBSK.

### RNA and Morpholino injections

mRNAs were synthetized using the SP6 mMessage mMachine kit (Ambion). RNAs were diluted in 0.1M KCl 0.2% Phenol Red. The following constructs and quantities were used: Acvr2Aa-GFP-pCS2+ (25 pg, this study), Acvr2Ba-GFP-pCS2+ (25 pg, this study), CA-SMAD2-pCS2+ (20pg^55^). CD44a-GFP-pCS2+ (50 pg, this study). GFP-Spaw-pCS2+ (20pg^46^). Histone2B-mRFP-pCS2+ (12.5 pg^56^). mRFP-SARA-pCS2+ (25 pg^50^). Myo1G-pCS2+ (50pg, this study). Myo1G-GFP-pCS2+ (50pg, this study). For Spaw^57^, Cyclops^58^ and Squint^36^ different concentrations used in individual experiments are indicated in the figures.

The previously reported *dnaaf1* Morpholino 5’-ATGCACTGTAATTTACCAAGTCAGG-3’^40^ was injected at a concentration of 500 µM diluted in 1x Danieau 0.2% Phenol Red.

For rescuing the cardiac jogging defects of *myo1g* mutant by Spaw mRNA injection, a mix of Spaw and GFP RNA was co-injected into one blastomere of two cell stage embryos. At bud stage embryos with a unilateral segregation of GFP expressing cells were selected using a fluorescent dissection scope (Leica M205FA), and grown further to score for cardiac jogging and looping phenotypes.

### RNA *in situ* hybridization

Whole mount RNA in situ hybridizations were performed as previously described (Thisse and Thisse, 2008). For the following genes probes were transcribed from previously reported plasmids: spaw-pGEMT^57^, lefty1-pBSK^31^, lefty2-pBSK^33^, pitx2c-pBSK^59^, foxa1-pBSK^60^, cyclops-pBSK^36^, squint-pBSK (this study), dand5-pBSK^24^, sox32-pBSK^61^, sox17-pBSK^62^, odad1-pME18S-FL3^63^, dnah9-pCRII^64^, foxj1a-pBSK^65^, cmlc2-pCS2^66^, gdf3-pBSK^63^, Her4.1-pBSK (this study), Her15.1-pBSK (this study). The elovl6 probe was transcribed from a PCR product containing a T7 promotor sequence at the 3’ end. elovl6 was amplified from genomic DNA using the forward primer 5’–CCCGTCCCATGTGCAGAACATTG–3’ and the reverse primer 5’–GGTGTCCATTGTGCTCGTGTGTCTCCCTATAGTGAGTCGTATTACGC– 3’.

### qPCR analysis

qPCR was performed using PowerUP SYBR Green Master Mix (Applied Biosystems) in an Applied Biosystems Step-One PCR system. Individual reactions were performed in triplicates to account for pipetting errors. For sample preparation, whole cell mRNA was isolated from 50 embryos using TRI-Reagent (Sigma). Reverse transcription was performed on 2.5 µg of RNA using Superscript III (Invitrogen) to generate cDNA. Fold changes in gene expression were normalized to the internal control gene *36b4*. The primers used for the amplification reactions are as follows: lefty1: forward 5’-AGAGGAGTTTGGGTCTAGTGG-3’, reverse 5’-TACGGAGAGAGGAAATGCG-3’. Spaw: forward 5’-TGACTTCGTCCTGAGCTTGA-3’, reverse 5’-TCAAGCTCAGGACGAAGTCA - 3’. 36b4: forward 5-ACGTGGAAGTCCAACTACT-3’, reverse: 5’-GTCAGATCCTCCTTGGTGA-3’. For estimating relative gene expression, the Ct values at 40 cycles of qPCR amplification were used according to the ΔΔCT method^67^. Individual data points in figures documenting qPCR experiments correspond to technical replicates. Complete statistical information including numbers of biological and technical replicates is provided in the Supplementary statistical information file.

### Immunocytochemistry

Dechorionated embryos were fixed for 1.5 hours at Room temperature in PEM (80 mM Sodium-Pipes, 5 mM EGTA, 1 mM MgCl2) - 4% PFA - 0.04% TritonX100 and then washed 2 × 5 min in PEMT (PEM - 0.2% TritonX100), 10 min in PEM 50 mM NH4Cl, 2 × 5 min in PEMT.

### Microscopy and image analysis

Embryos were mounted in 0.75% low melting agarose (Sigma) in glass bottom dishes (Mattek) for confocal imaging. Imaging was performed using Laser scanning confocal microscopes (Zeiss LSM710, 780 and 880) using 40 x Water immersion (NA1.1) objectives. Airyscan super-resolution imaging was performed on a Zeiss LSM 880 system. In situ hybridizations were documented on a Leica M205 microscope with a Lumenera Infinity camera. Image analysis was performed using ImageJ (http://rbs.info.nih.gov/ij/).

### Statistical Analyses

Appropriate statistical tests were selected for each experiment based on the nature of the comparison (bi-or multifactorial, ordinal or categorical data), data distribution and variance. Statistical analysis and representations were performed using R. Complete informations regarding the applied statistical tests, test statistics, sample sizes and displayed error bars for all experiments is provided in the supplementary statistical information file.

## ACKNOWLEDGEMENTS

This study was supported by an ARC project grant (PJA20181208167) and the ANR DroZeMyo (ANR-17-CE13-0024-02) (MF). AJK benefited from a 4^th^ year PhD fellowship from La Ligue Contre le Cancer. Confocal microscopy was performed with the help of the iBV PRISM imaging platform. We thank M.Gonzalez-Gaitan, C.P.Heisenberg, T.Lepage, S.Lopes, and C. & B.Thisse for the sharing of reagents. We are grateful to S.Polès, R.Rebillard and G.Dupuy for technical assistance and excellent fish care.

## AUTHOR CONTRIBUTIONS

The genetic analysis of *myosin1* function in zebrafish Nodal signaling was entirely performed by A.J.K. F.B. generated reagents used in the present study. M.F. designed the study, performed experiments, analyzed the data and wrote the manuscript.

## COMPETING INTERESTS

The authors declare no competing interests.

## Supplementary Information

**Supplementary Figure 1:**
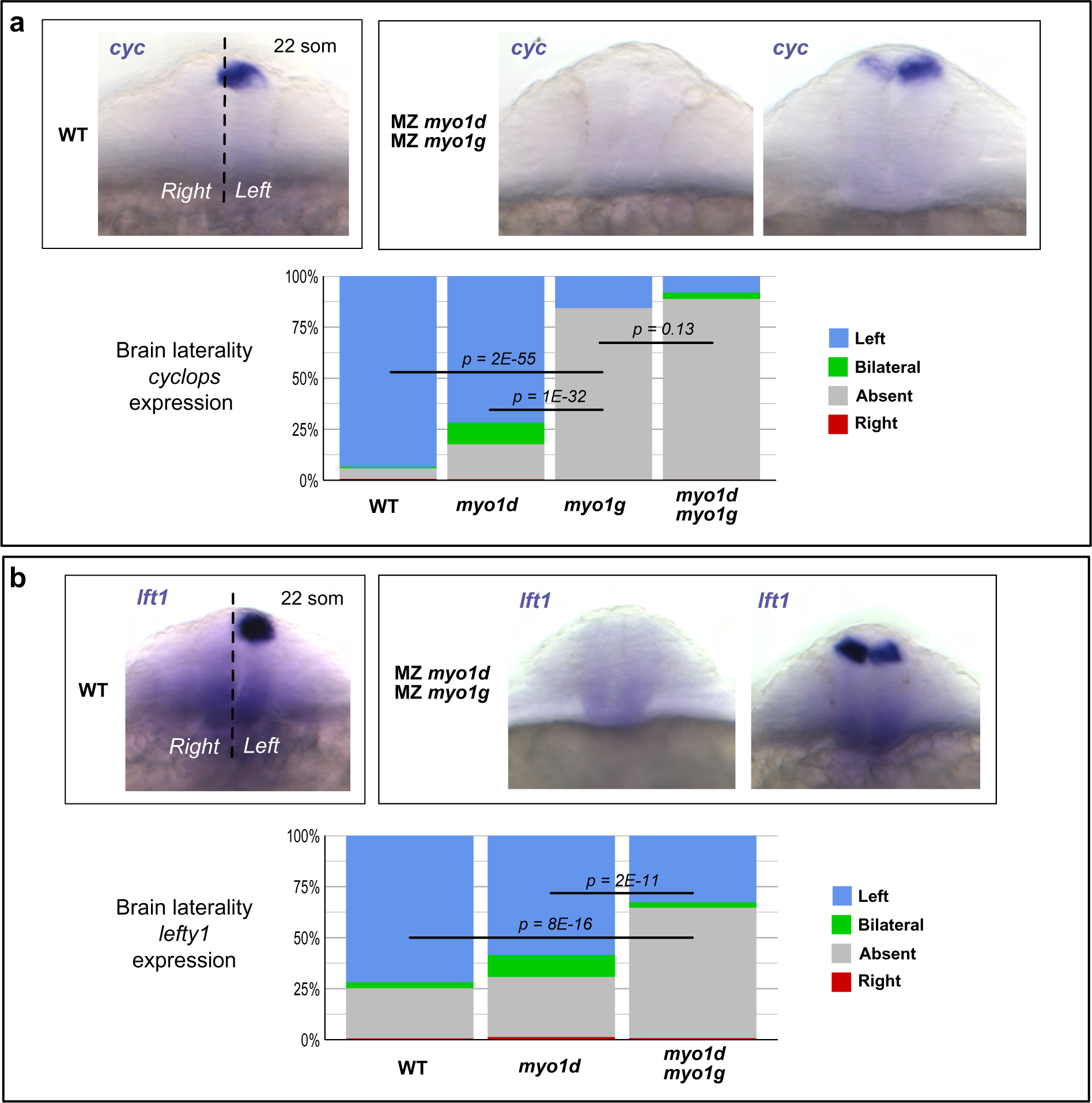
*myo1g* is required for brain laterality. **a,b** *myo1g* loss of function impairs the asymmetric expression of the Nodal ligand *cyc* (**a**) and the Nodal target gene *lft1* (**b**) in the dorsal epithalamus. **a,b** Frontal views of the brain, dorsal up.

**Supplementary Figure 2:**
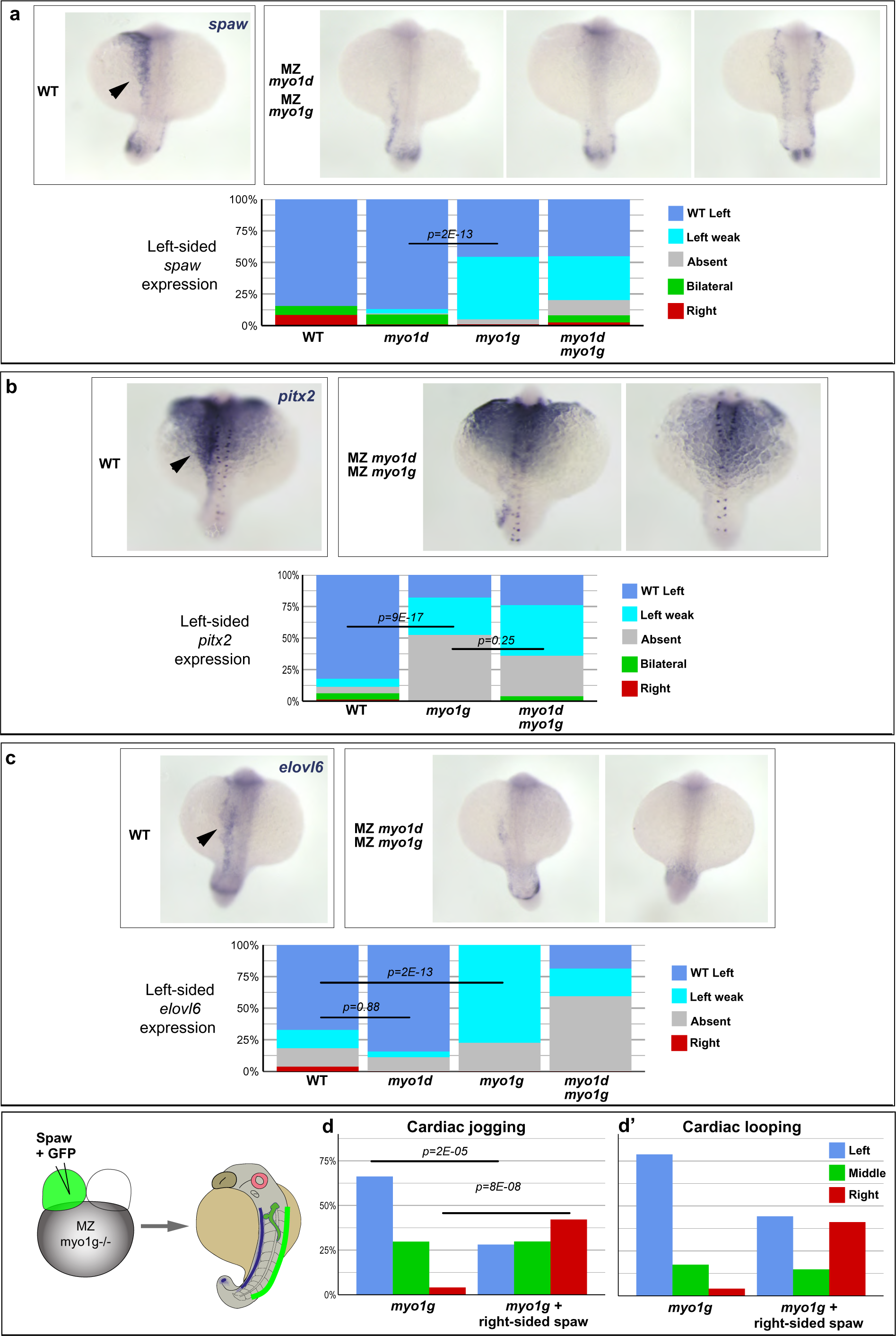
Nodal pathway gene expression is impaired in *myo1g* mutants. **a-c** *myo1g* mutants display a reduced expression of the Nodal ligand *spaw* (**a**), the Nodal effector *pitx2* (**b**) and the Nodal target gene *elovl6* (**c**) in the Left Lateral Plate Mesoderm (LLPM, black arrows). **d,d’** Right-sided expression of Spaw rescues neither the cardiac jogging (**d**) nor the cardiac looping (**d’**) defects of *myo1g* mutants. The embryos displayed in **d,d’** are derived from the same series of experiments also used in Fig. 2e,e’. **a-c** Dorsal views of 22 somites stage embryos, anterior up.

**Supplementary Figure 3:**
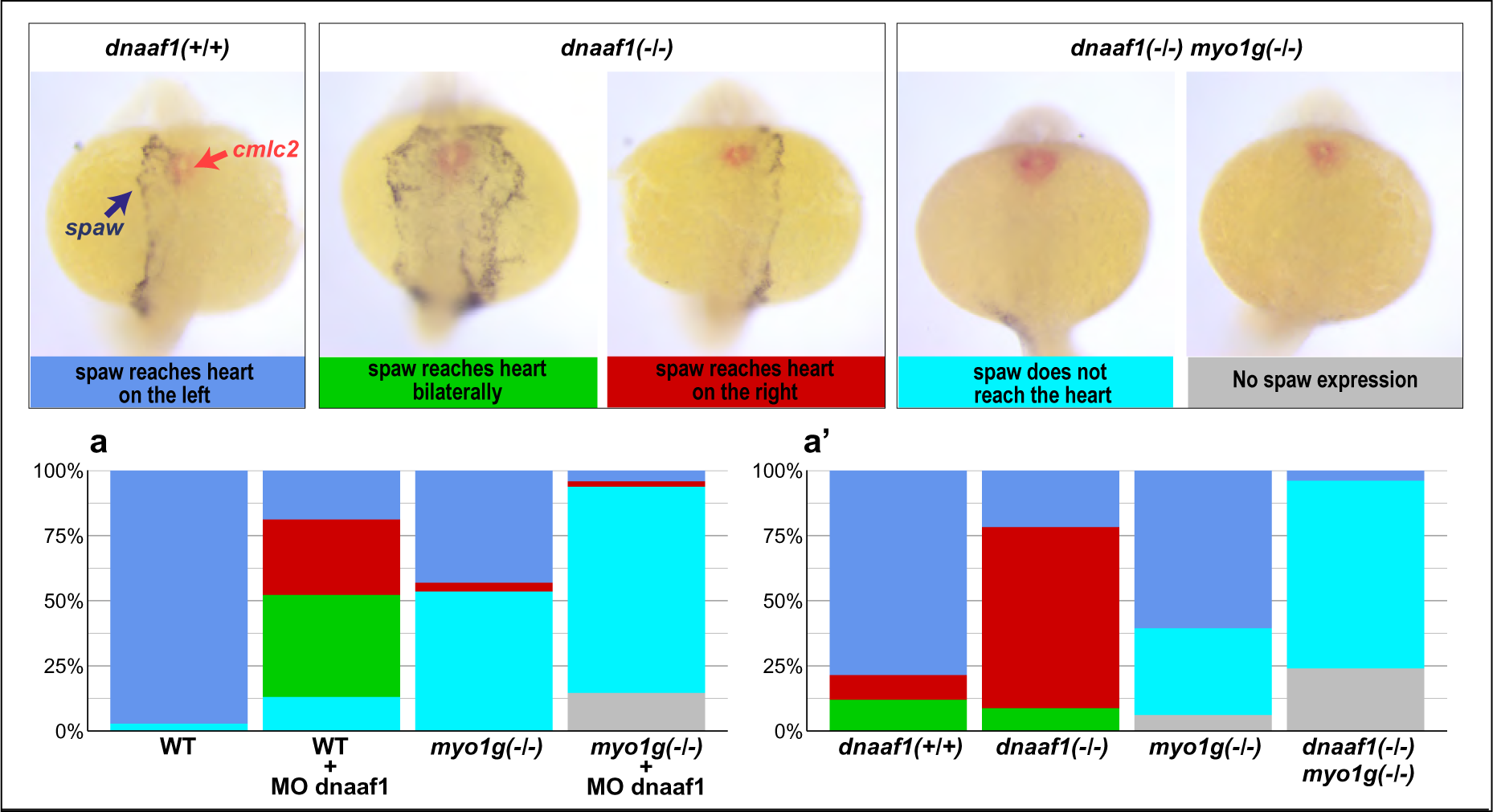
*spaw* ligand expression fails to reach the cardiac primordium in *myo1g* mutants. **a,a’** Double *in situ* hybridization for *spaw* (purple) and the cardiac marker *cmlc2* (red) shows that spaw expression reaches the cardiac primordium on the left, right or both sides of the embryo in most WT control and LRO flow-deficient dnaaf1 morphant (**a**) or *dnaaf1* mutant (**a’**) embryos. In contrast, *spaw* expression fails to reach the cardiac primordium in a large fraction of *myo1g* single mutants (**a,a’**), *myo1g* mutant dnaaf1 morphants (**a**) or *myo1g dnaaf1* double mutants (**a’**). The dataset used in this figure is the same that is also displayed in Fig. 3c, c’. Dorsal views of 22 somites stage embryos, anterior up.

**Supplementary Figure 4:**
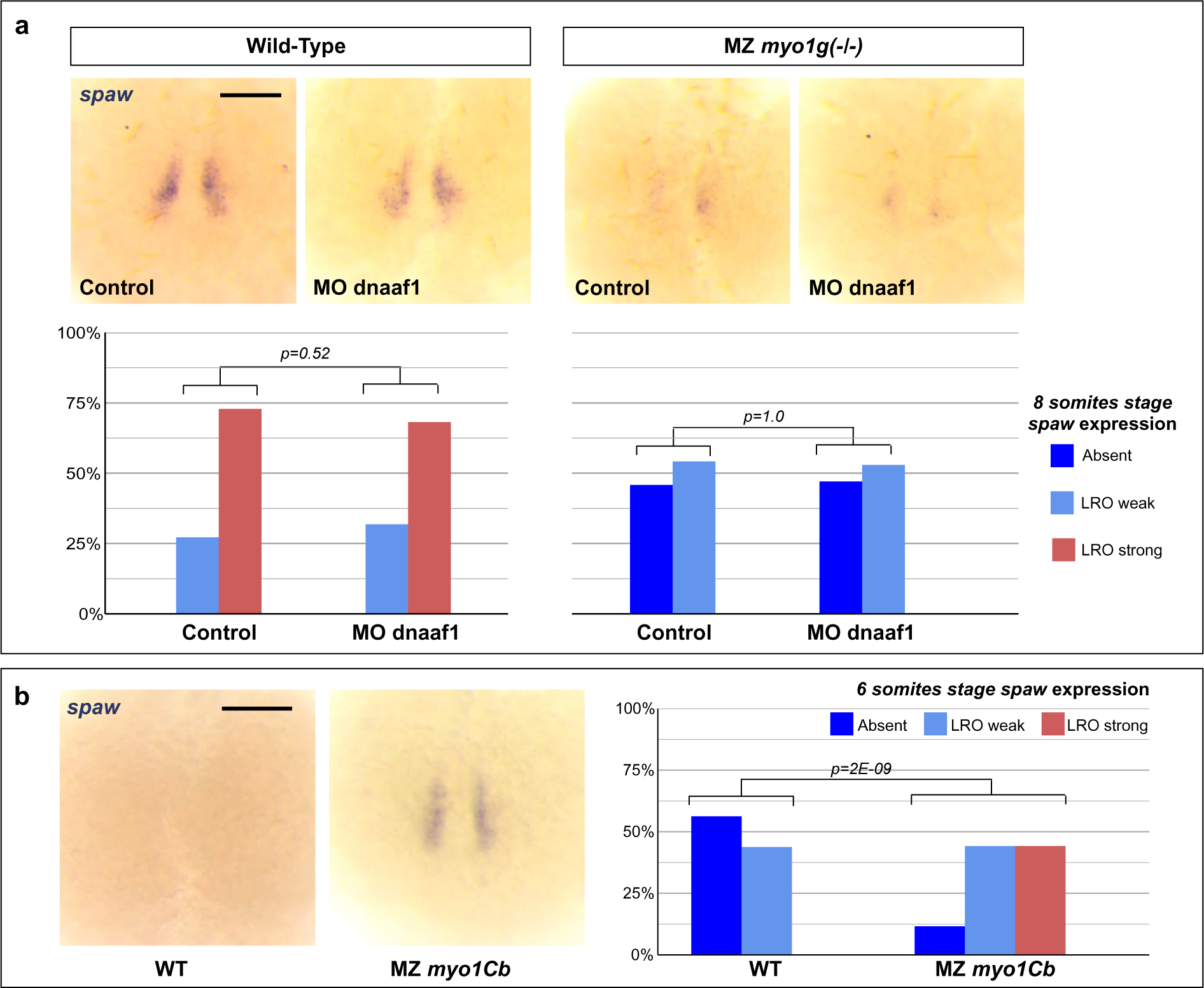
Myosin1 proteins control the initiation of *spaw* expression at the LRO independently of the LRO flow. **a** The 8 somites stage expression of *spaw* in the LRO region is unaffected by the loss of the LRO flow (dnaaf1 morphants), but reduced in *myo1g* mutants. **b** Conversely, 6 somites stage mutants for the *myo1d/g* antagonist *myo1Cb* present a premature expression of *spaw* in the LRO region. Vegetal views of the LRO region, anterior up. Scale bars: 100 µm.

**Supplementary Figure 5:**
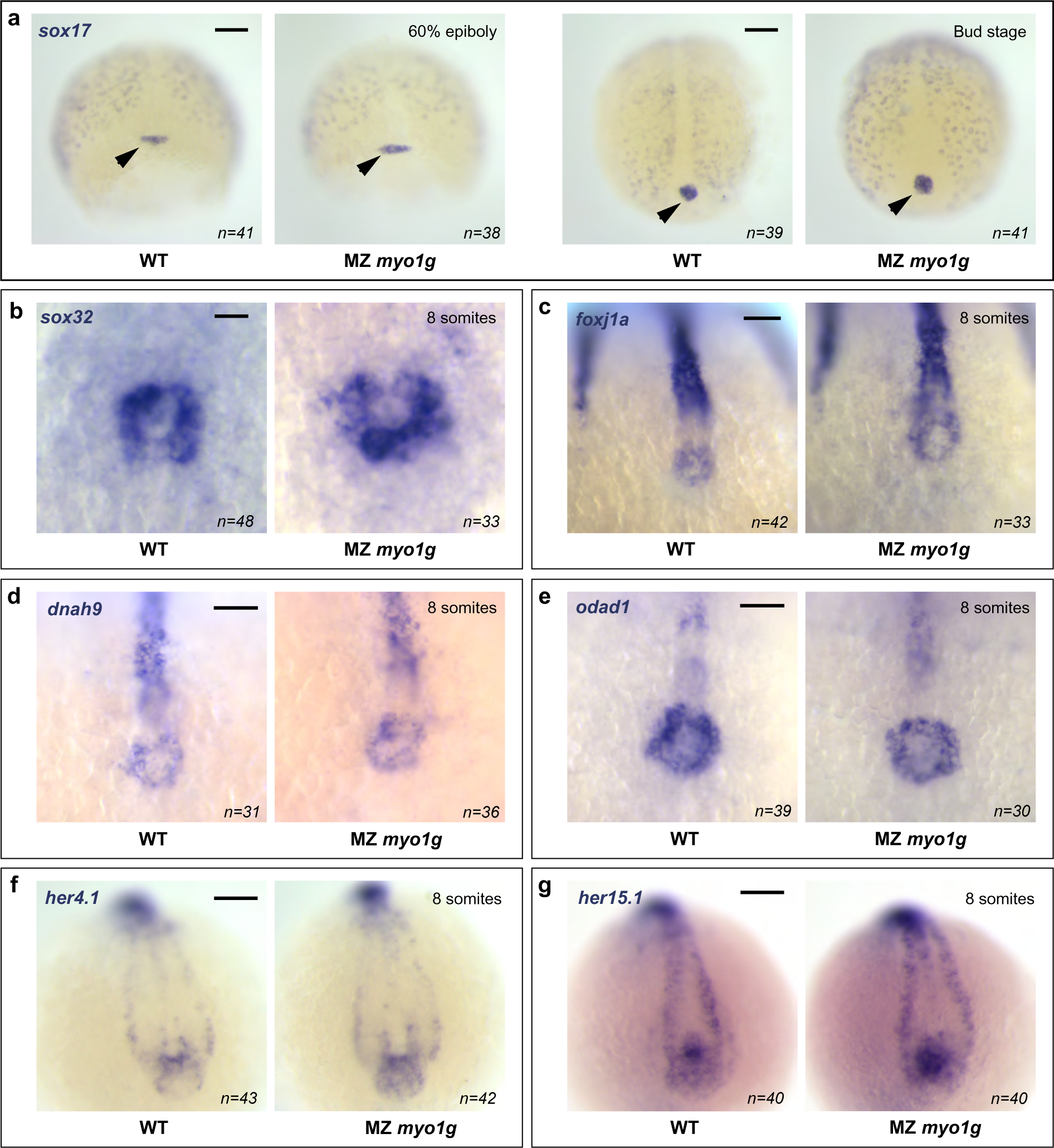
Myosin1G is dispensable for LRO specification. **a** *In situ* hybridization for the endodermal marker *sox17* shows that *myo1g* loss of function has no effect on the specification and behavior of the dorsal forerunner cells (black arrows) that are the precursors of the zebrafish LRO. **b** Accordingly, *myo1g* mutants present a normal expression of the endodermal marker *sox32* in the LRO/Kupffer’s Vesicle of 8 somites stage embryos. **c-e** The ciliary motility genes *foxj1a* (**c**), *dnah9* (**d**) and *odad1* (**e**) are expressed normally in *myo1g* mutant LROs. **f,g** Analysis of the Notch signaling targets *her4.1* (**f**) and *her15.1* (**g**) indicates that **myo1g** loss of function has no effect on Notch signaling at the LRO. **a** Dorsal views, anterior up. **b-g** Vegetal views of the LRO region, anterior up. Scale bars: **a,f,g** 100 µm. **b** 20 µm. **c-e** 50 µm.

**Supplementary Figure 6:**
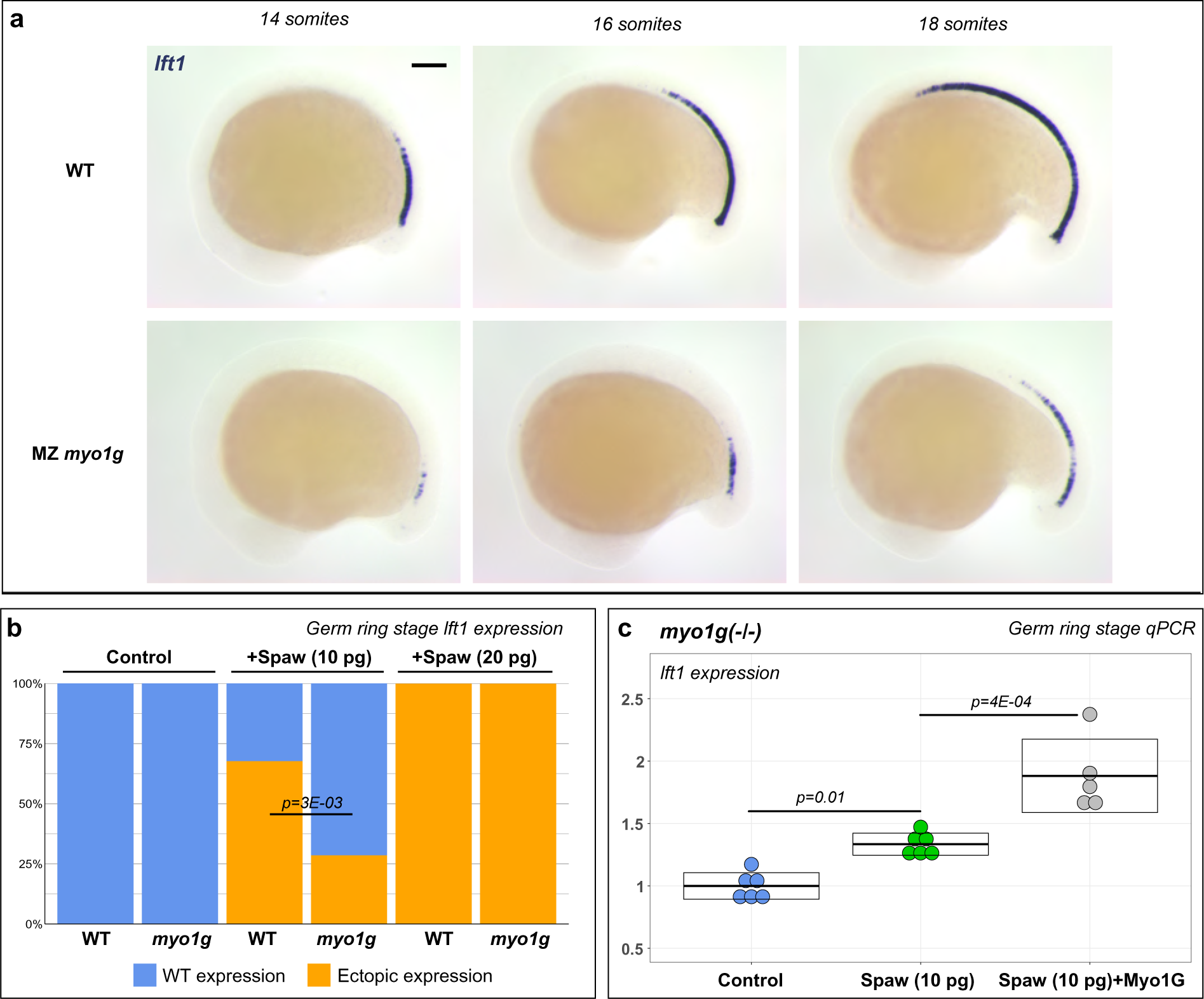
Myosin1G promotes Spaw signaling. **a** *In situ* hybridization shows that *myo1g* mutants present a reduced anterior propagation of the Spaw target gene *lft1*. Lateral views, anterior to the left. Scale bar: 100 μm. The displayed embryos are part of the data set used to estimate the rate of *lft1* propagation in Fig. 4c. **b** *myo1g* mutants display a weaker induction of the nodal target gene *lft1* in response to Spaw overexpression. Quantification for the experimental dataset displayed in Fig. 4d. **c** Wild-type Myo1G RNA injection significantly enhances the capacity of Spaw to induce *lft1* expression in *myo1g* mutants.

**Supplementary Figure 7:**
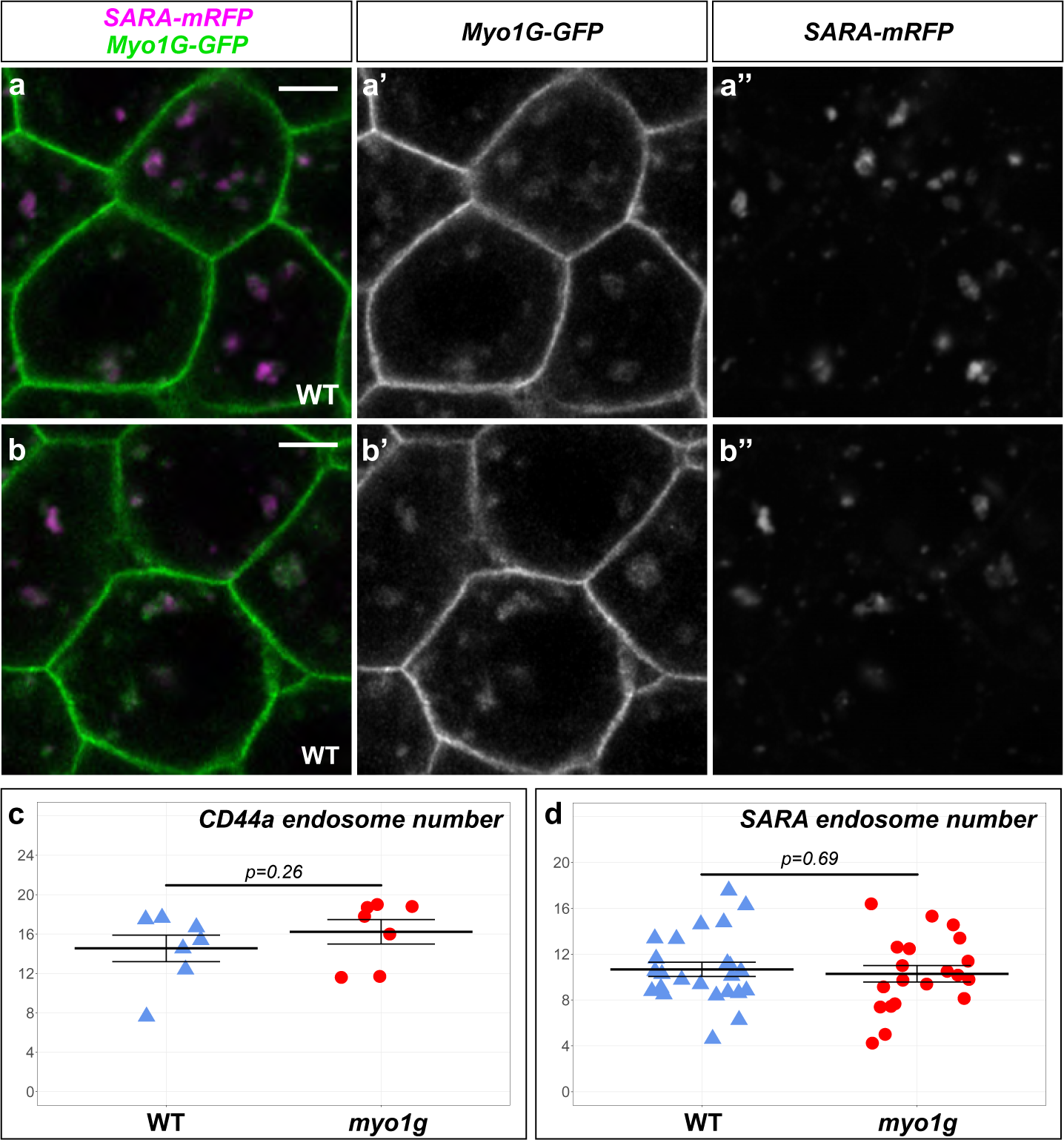
Myosin1G is associated with TGFβ signaling endosomes. **a,b** Airy scan super-resolution microscopy indicates that the signaling endosome marker SARA localizes to sub-domains of large Myo1G-positive compartments. Animal pole views of germ ring stage WT embryos. Scale bars: 5 µm. **c** *myo1g* mutants and their WT siblings display a similar number of CD44a endosomes. Quantification of the dataset displayed in Fig. 5k,l. **d** *myo1g* mutants and their WT sibling display a similar number of SARA endosomes. Quantification of the dataset displayed in Fig. 5m-p.

**Supplementary Figure 8:**
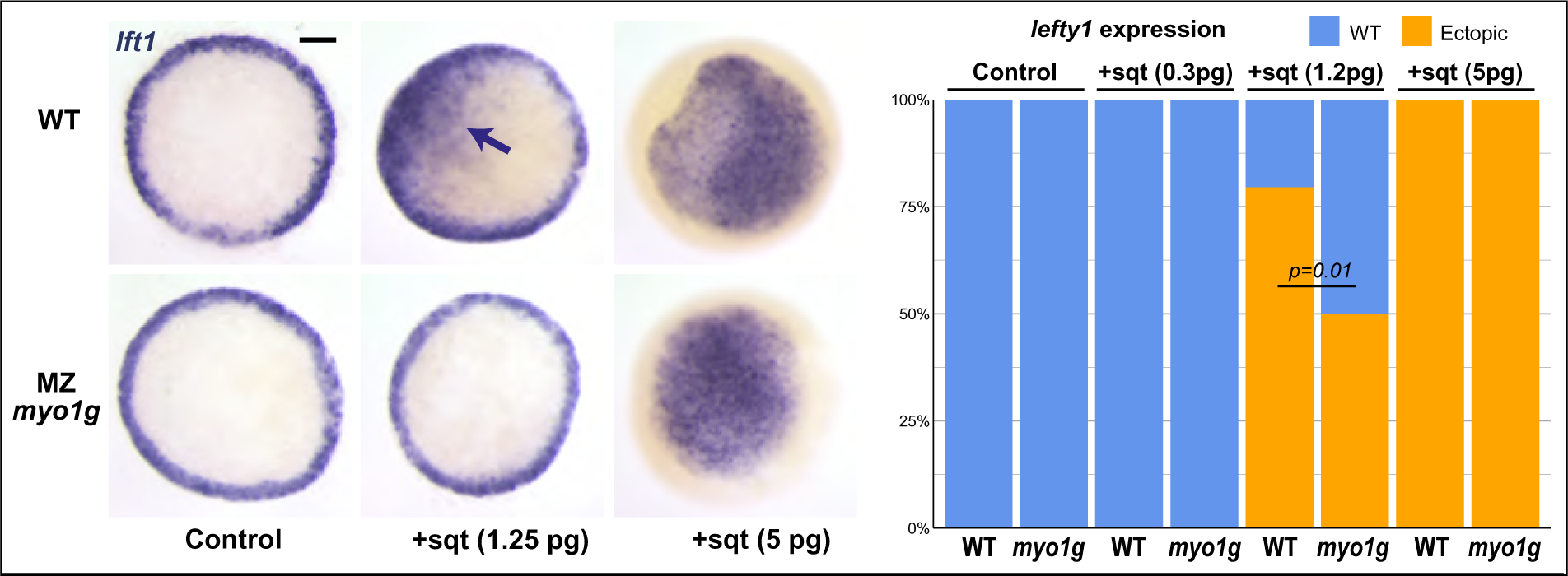
myo1g mutants present a reduced response to Squint overexpression. *myo1g* mutants display a weaker induction of the nodal target gene *lft1* in response to Sqt overexpression. While high amounts (5 pg) of Sqt induce similar ectopic *lft1* expression, a reduced *lft1* induction is observed in response to moderate amounts (1.2 pg) of Sqt RNA.

## SUPPLEMENTARY STATISTICAL INFORMATION

### Complete statistical information for the experiments reported in different display items

**Figure 1a:**
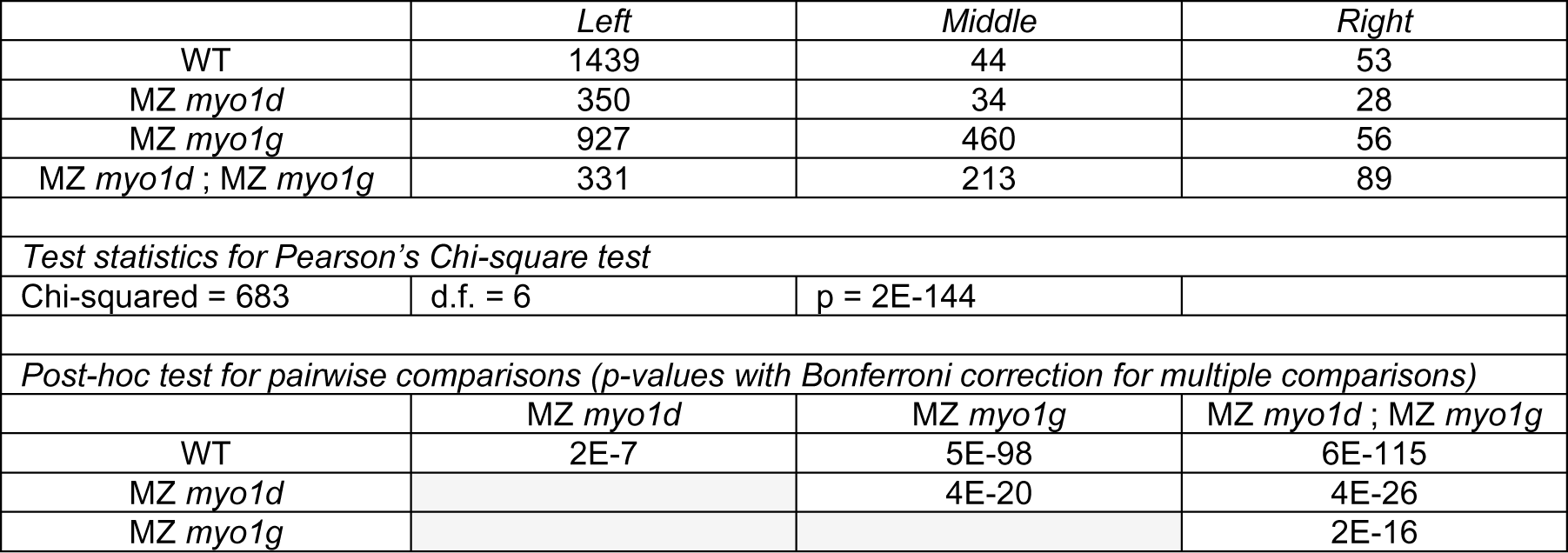
Cardiac jogging in *myosin1* single and double mutants.

**Figure 1a’:**
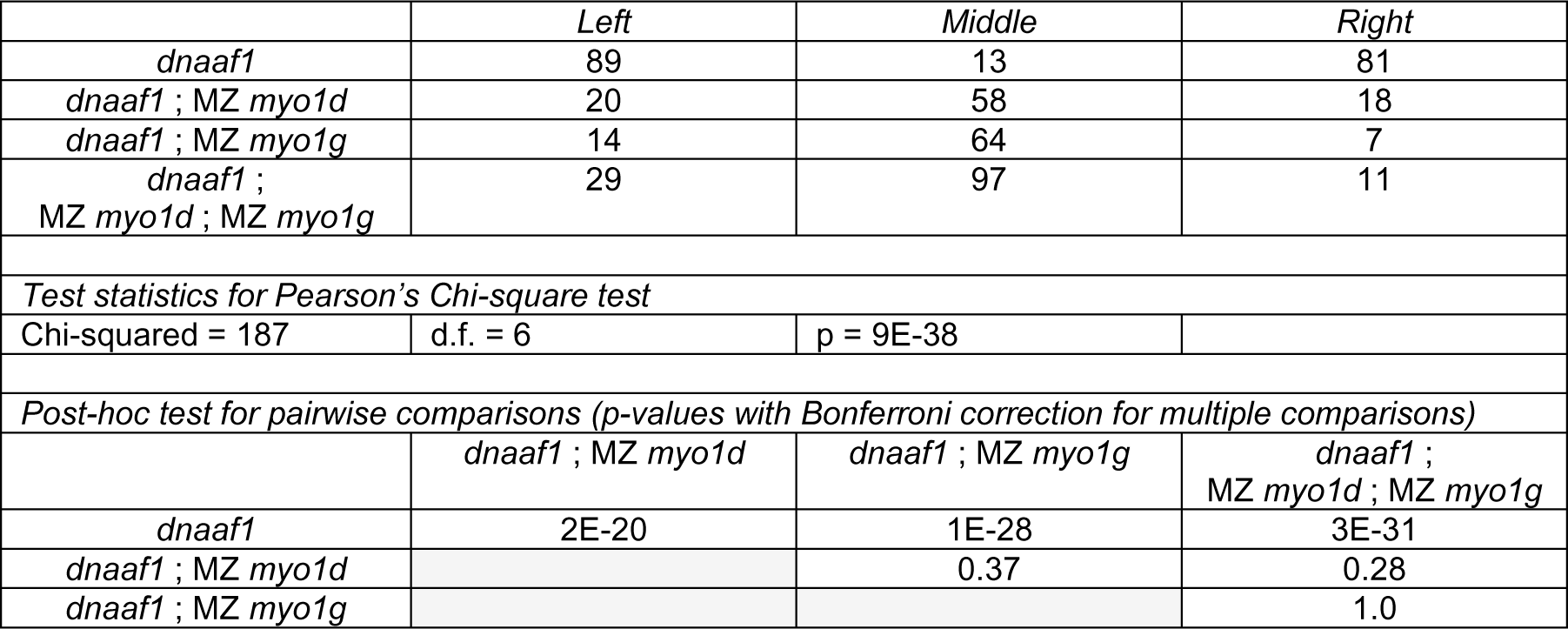
Cardiac jogging in *dnaaf1* single and *dnaaf1*; *myosin1* double and triple mutants.

**Figure 1b:**
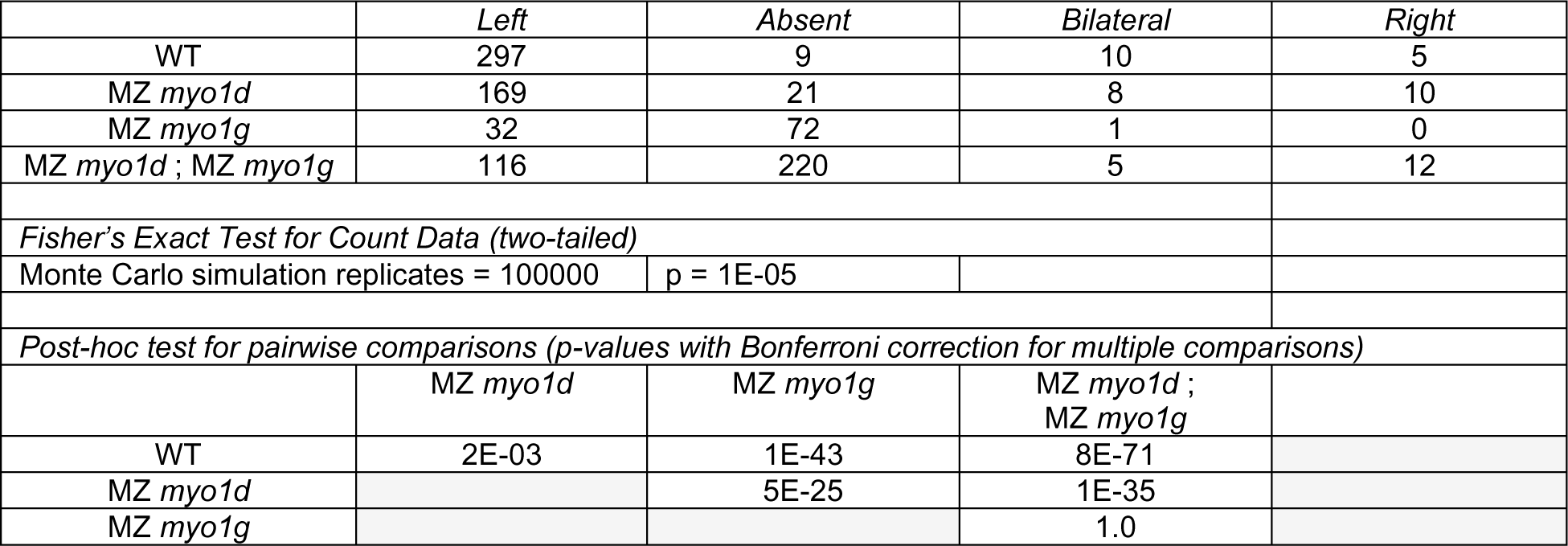
Brain *pitx2* expression in *myosin1* single and double mutants.

**Figure 1c:**
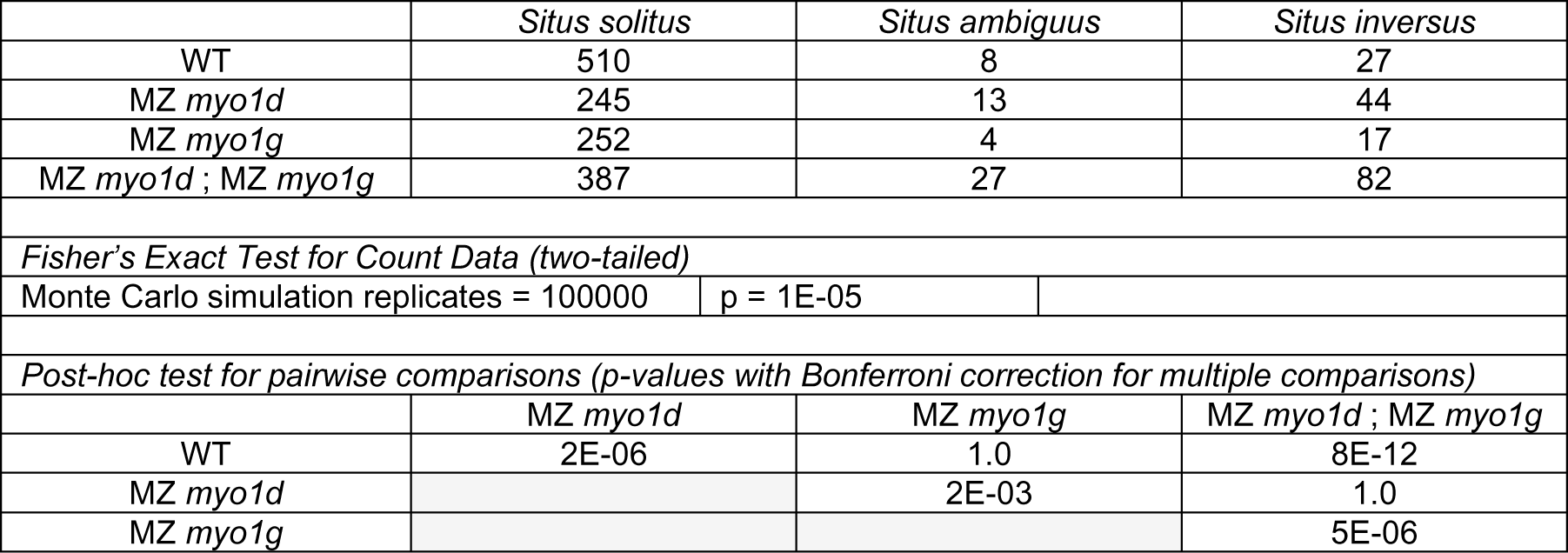
Visceral laterality in *myosin1* single and double mutants.

**Figure 1d:**
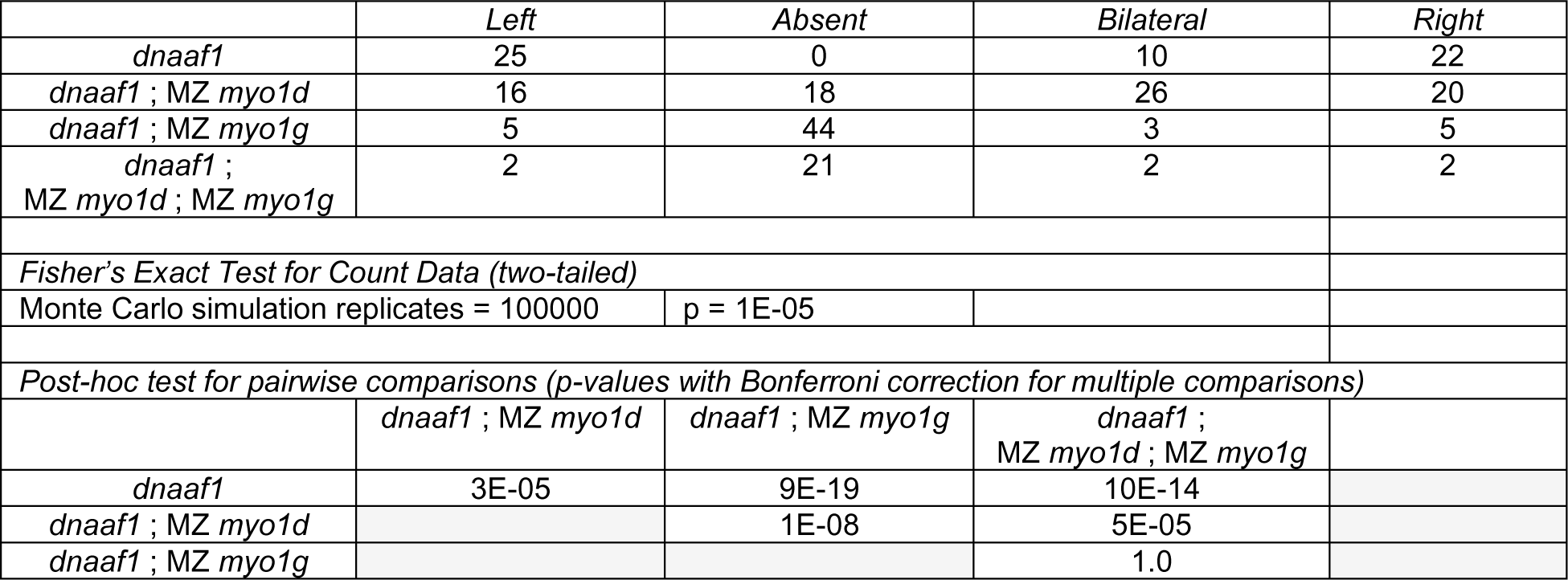
Brain *pitx2* expression in *dnaaf1* single and *dnaaf1*; *myosin1* double and triple mutants.

**Figure 1e:**
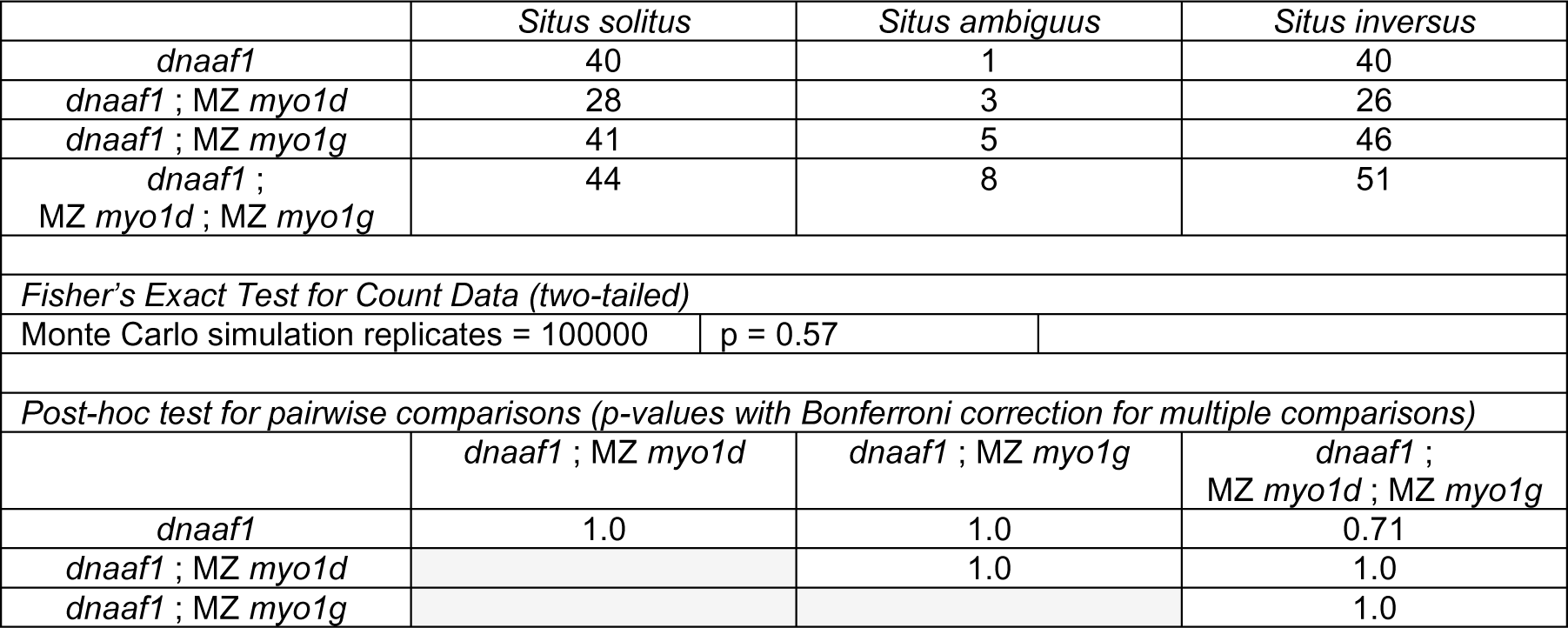
Visceral laterality in *dnaaf1* single and *dnaaf1*; *myosin1* double and triple mutants.

**Figure 2a:**
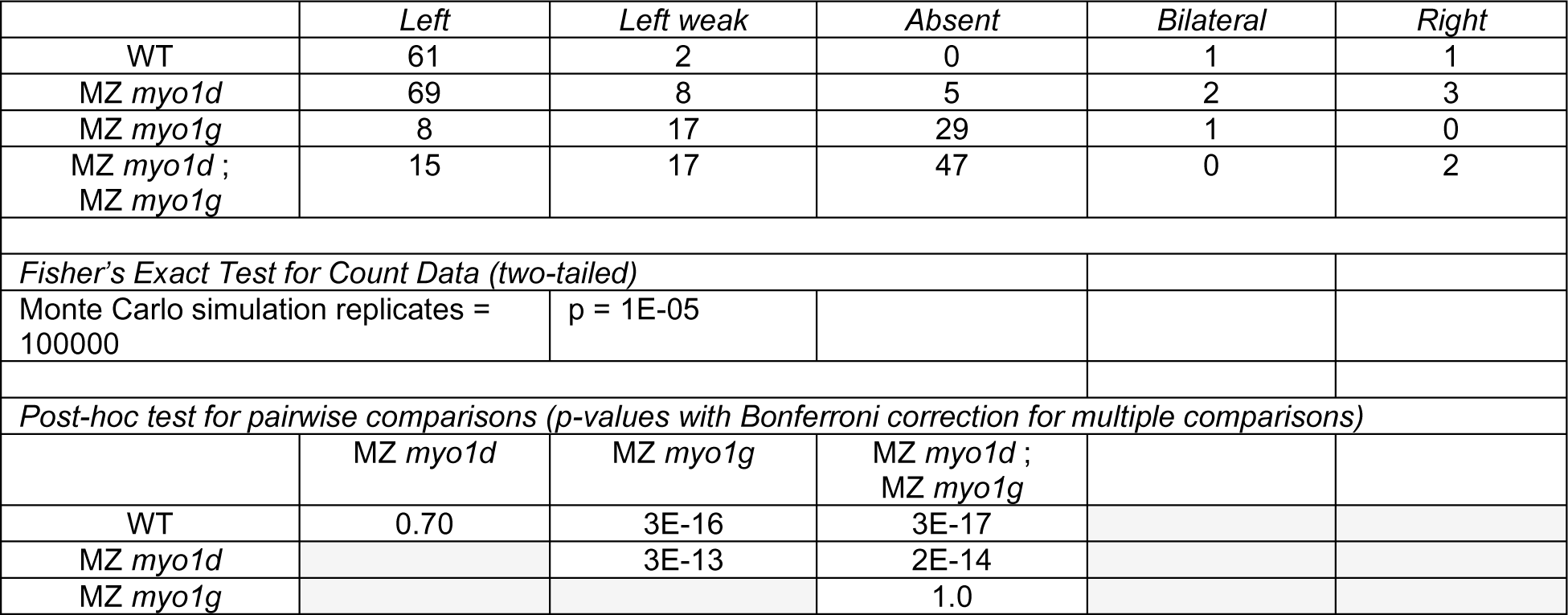
Cardiac *lefty2* expression in *myosin1* single and double mutants.

**Figure 2b:**
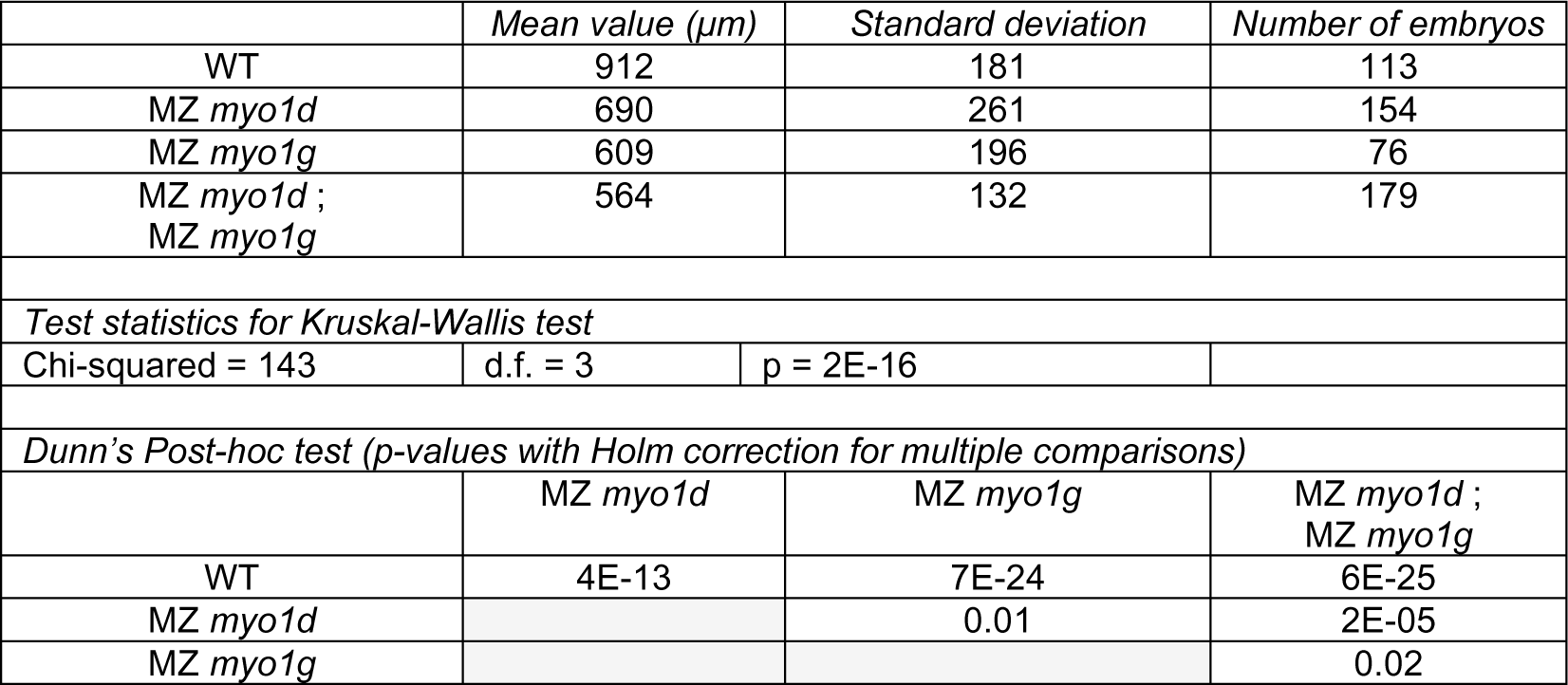
*southpaw* extension in the left lateral plate of *myosin1* single and double mutants.

**Figure 2c:**
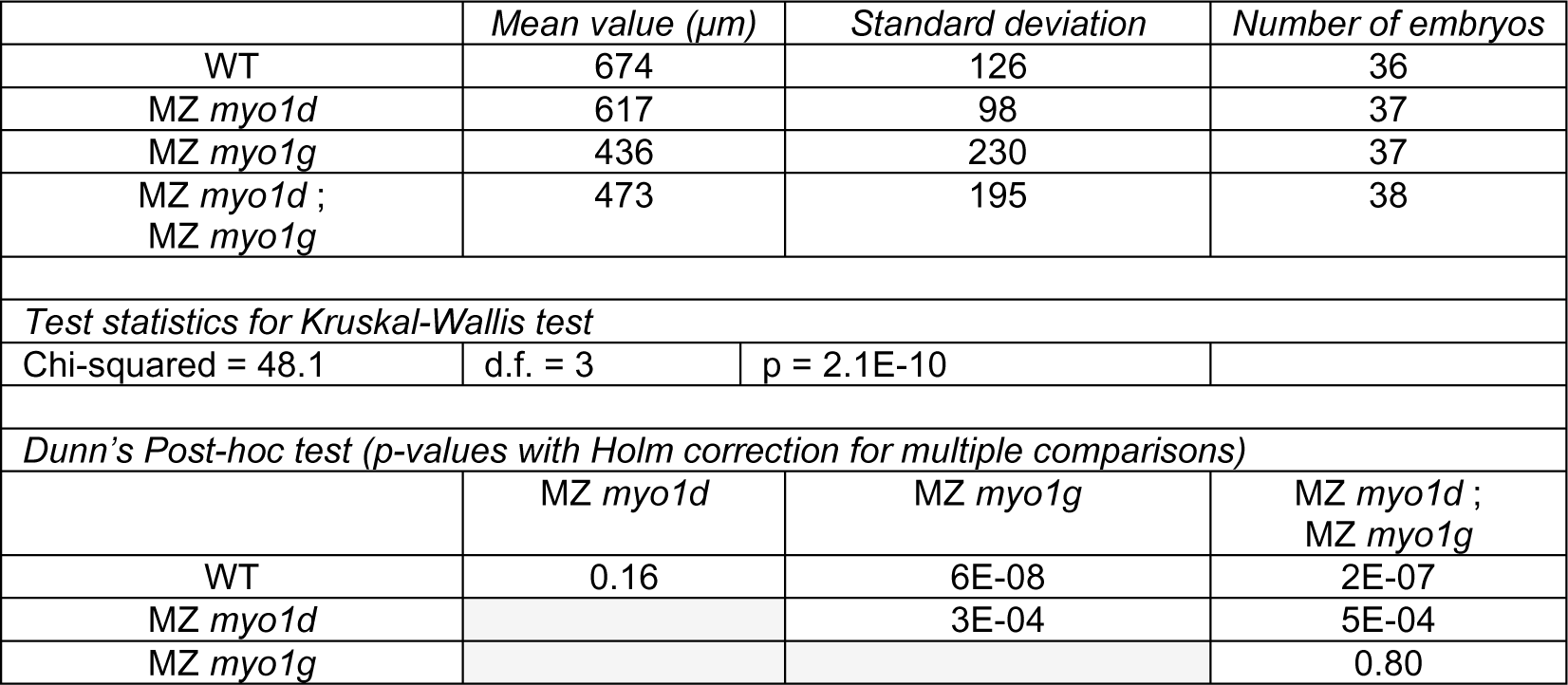
*pitx2* extension in the left lateral plate of *myosin1* single and double mutants.

**Figure 2d:**
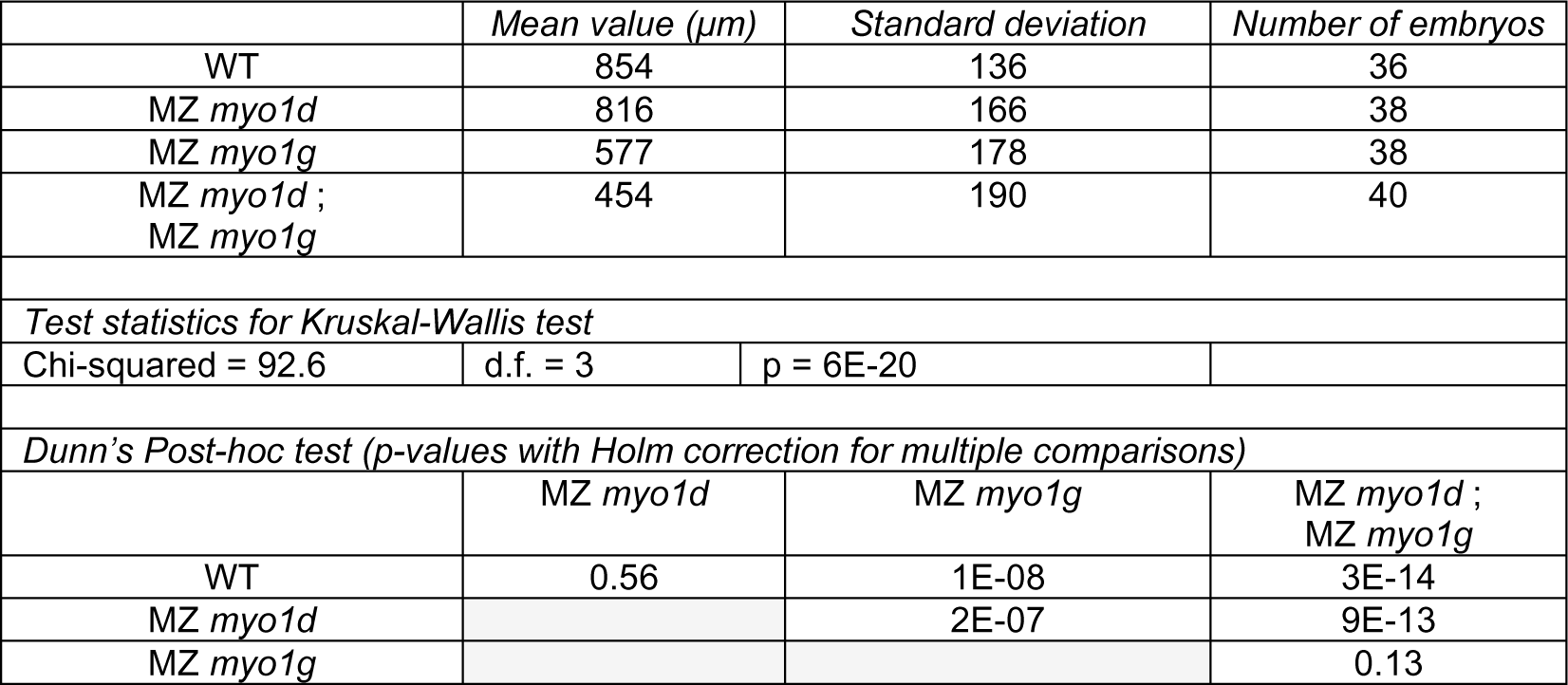
*lefty1* extension in the notochord of *myosin1* single and double mutants.

**Figure 2e and Supplementary Figure 2d:**
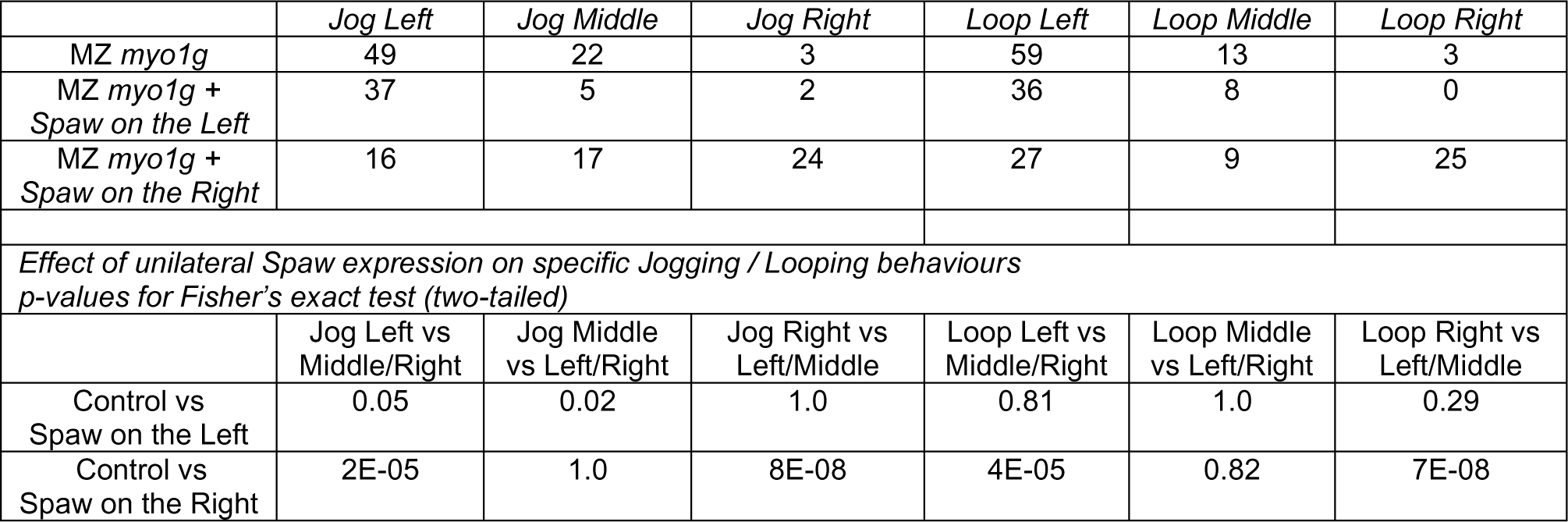
Cardiac jogging and looping in Spaw-injected myo1g mutants.

**Figure 3a:**
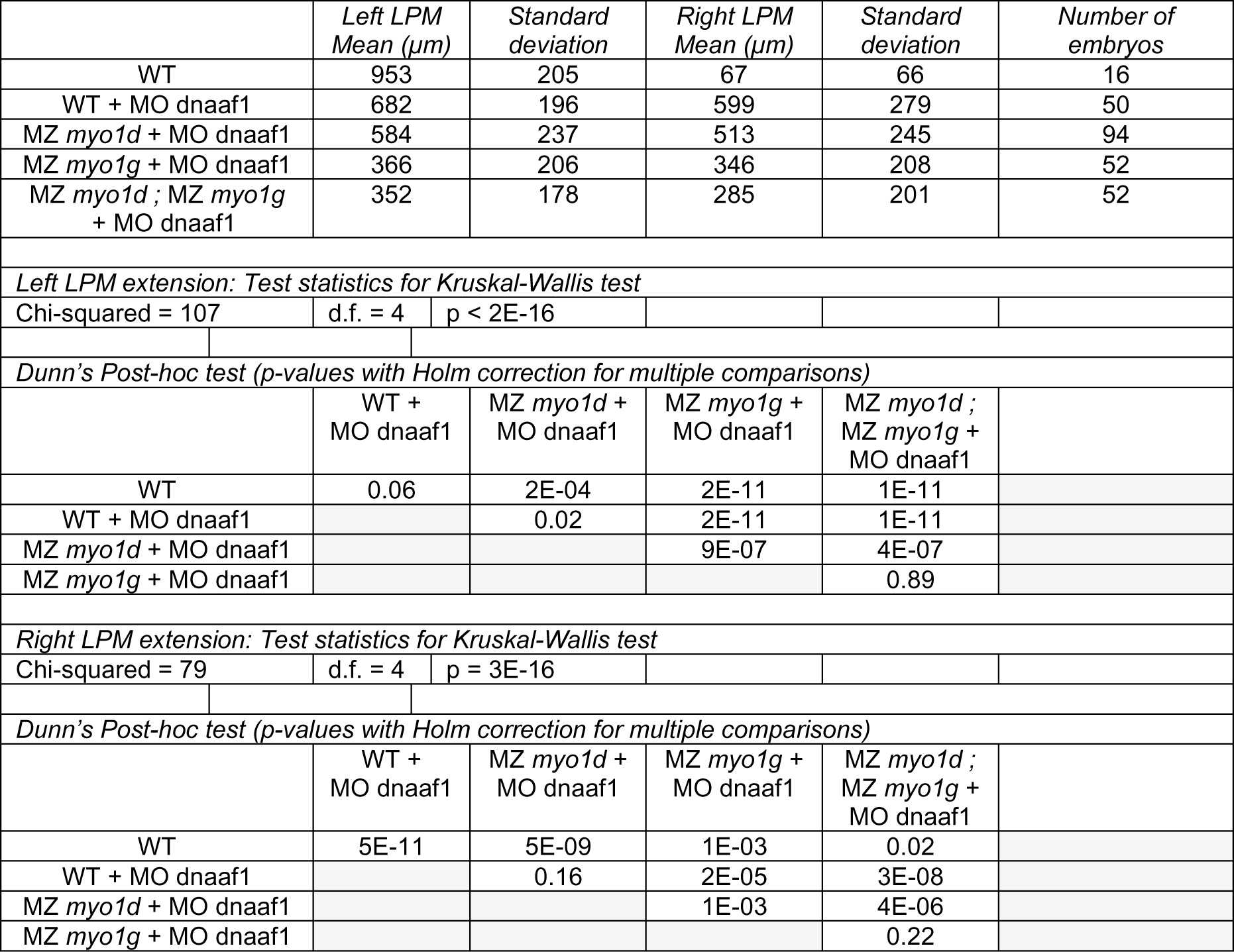
*southpaw* extension in the lateral plate of dnaaf1 Morpholino-injected *myosin1* mutants.

**Figure 3b:**
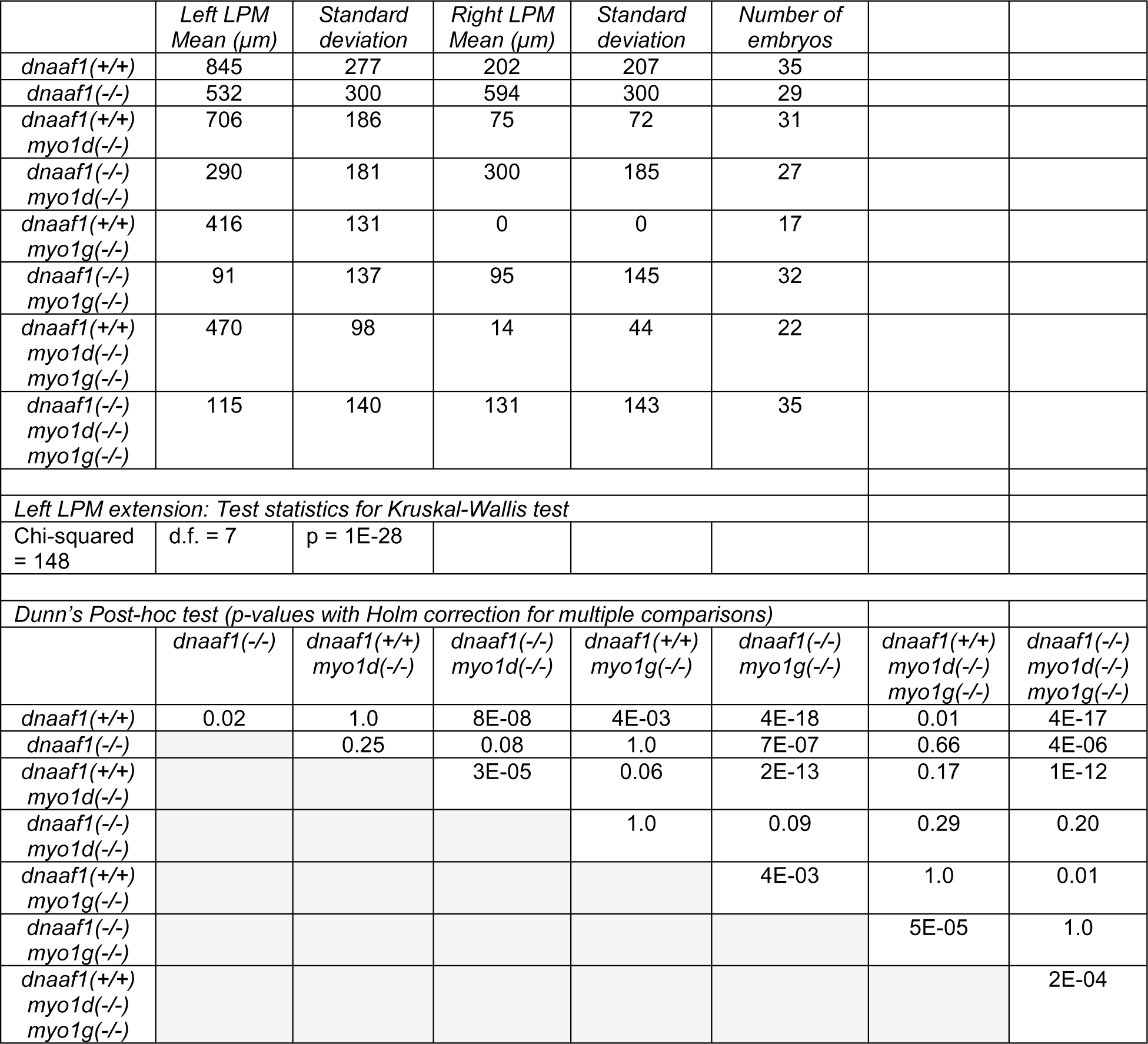
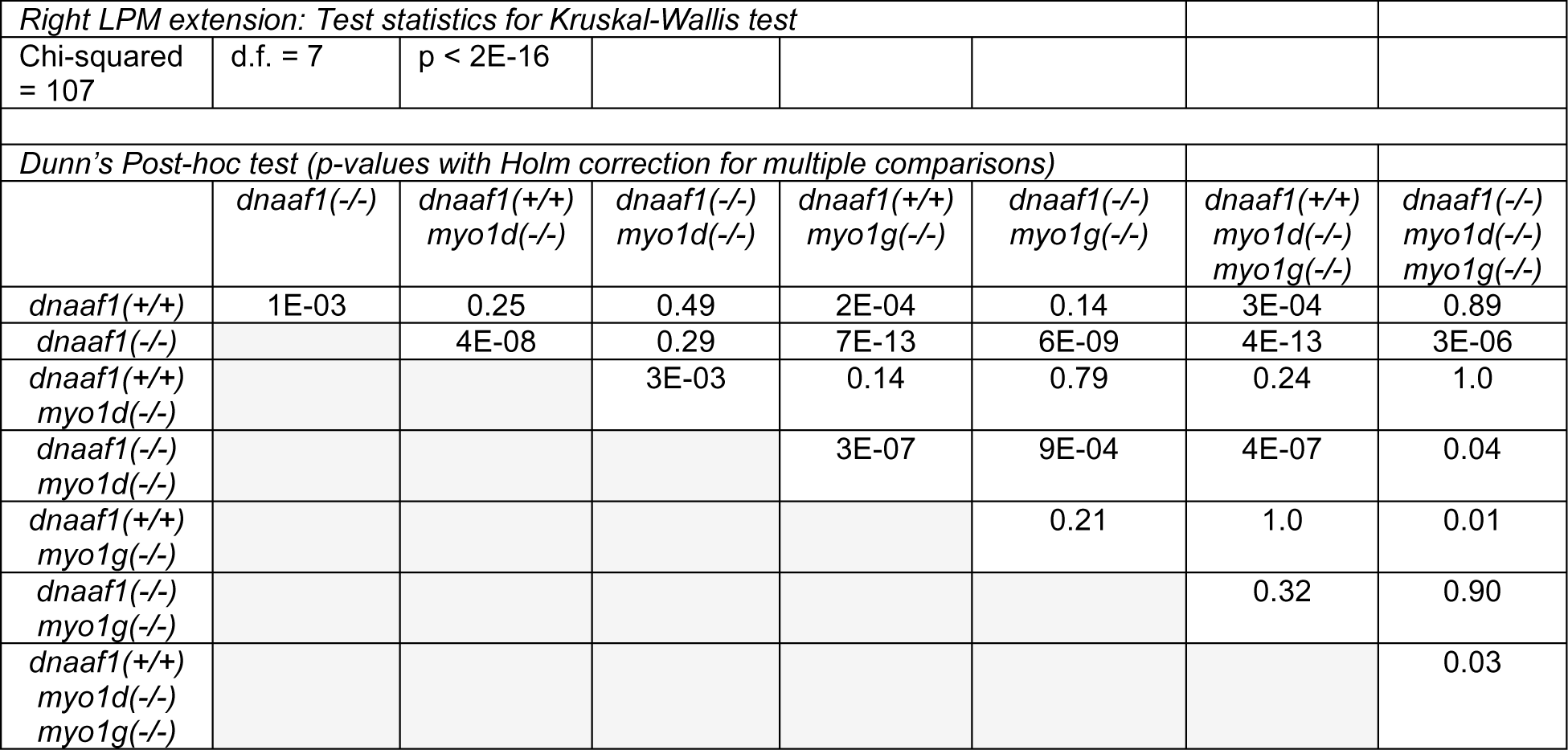
*southpaw* extension in the lateral plate of *dnaaf1 myosin1* double and triple mutants.

**Figure 3c and Supplementary Figure 3a:**
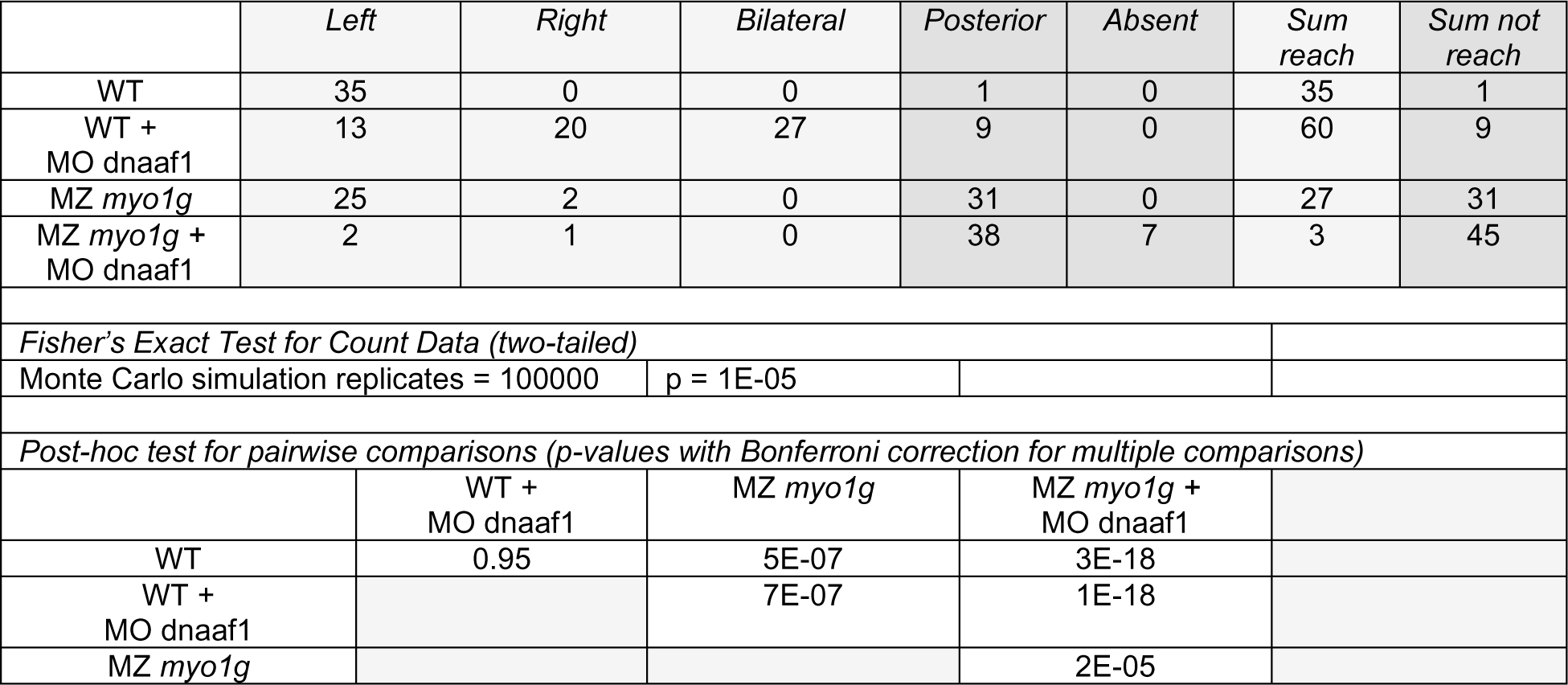
*southpaw* propagation to the cardiac primordium in dnaaf1 Morpholino-injected *myo1g* mutants. The table indicates the number of embryos in which *spaw* reaches the cardiac primordium in the Left LPM (Left), Right LPM (Right), Left and Right LPM (Bilateral), remains posterior to the primordium (Posterior) or is altogether absent (Absent), as displayed in Supplementary Fig. 3a. For statistical analysis, the former three and latter two categories are separately pooled into two categories for which *spaw* either reaches (Sum reach) or does not reach (Sum not reach) the primordium, as displayed in Fig. 3c.

**Figure 3c’ and Supplementary Figure 3a’:**
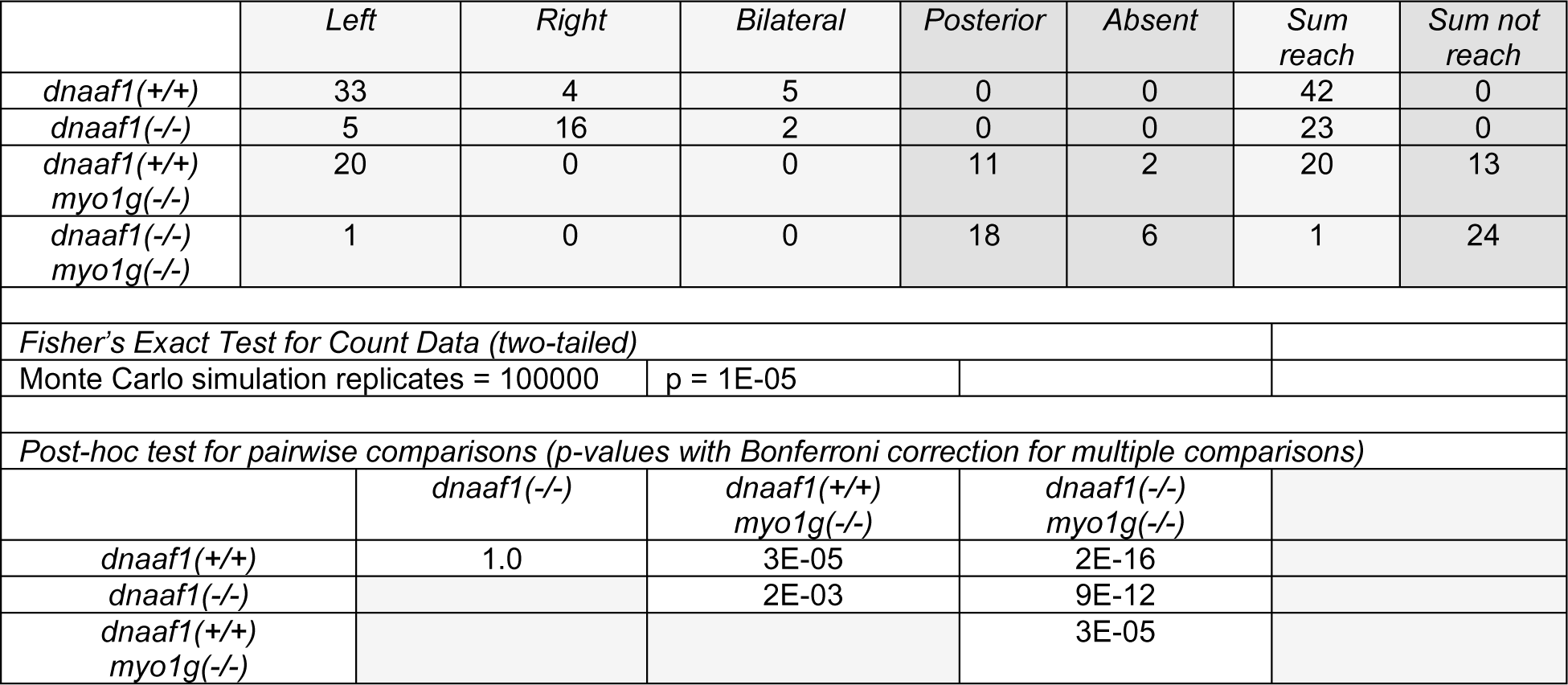
*southpaw* propagation to the cardiac primordium in *dnaaf1 myo1g* single and double mutants. The table indicates the number of embryos in which *spaw* reaches the cardiac primordium in the Left LPM (Left), Right LPM (Right), Left and Right LPM (Bilateral), remains posterior to the primordium (Posterior) or is altogether absent (Absent), as displayed in Supplementary Fig. 3a’. For statistical analysis, the former three and latter two categories are separately pooled into two categories for which *spaw* either reaches (Sum reach) or does not reach (Sum not reach) the primordium, as displayed in Fig. 3c’.

**Figure 4a:**
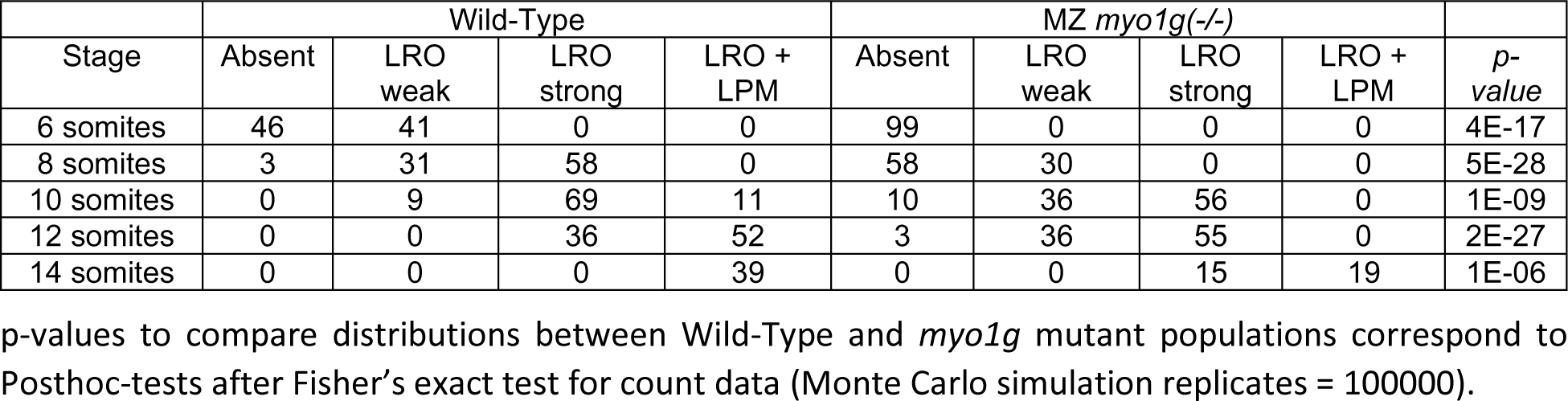
Timecourse of *myo1g* mutant *southpaw* expression.

**Figure 4b:**
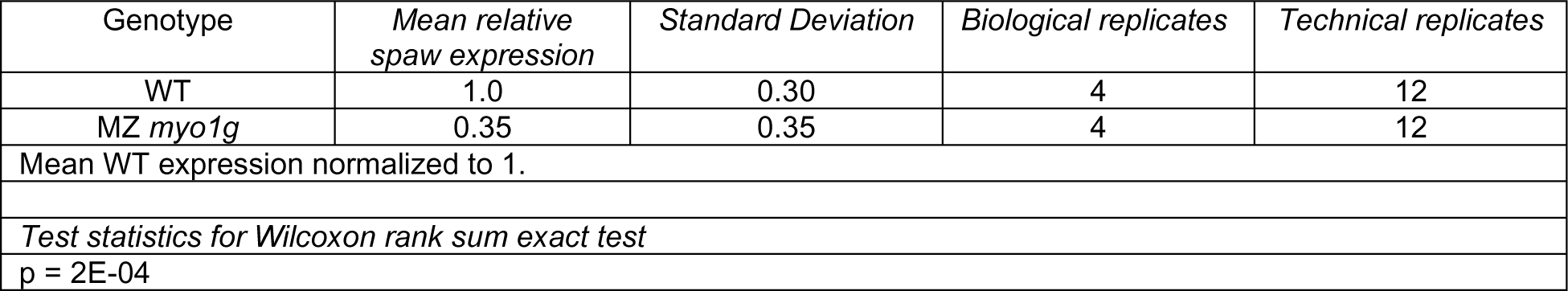
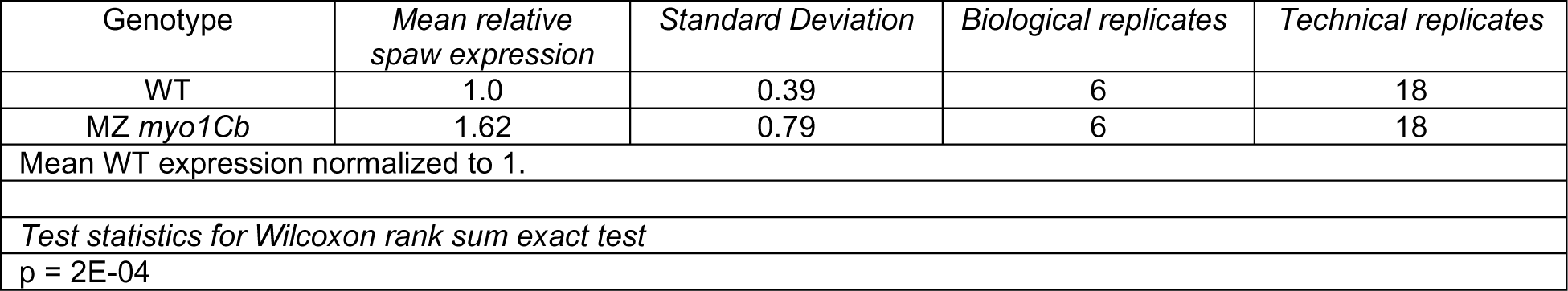
qPCR analysis of 8 somite stage *southpaw* expression in *myo1g and myo1Cb* mutants.

**Figure 4c:**
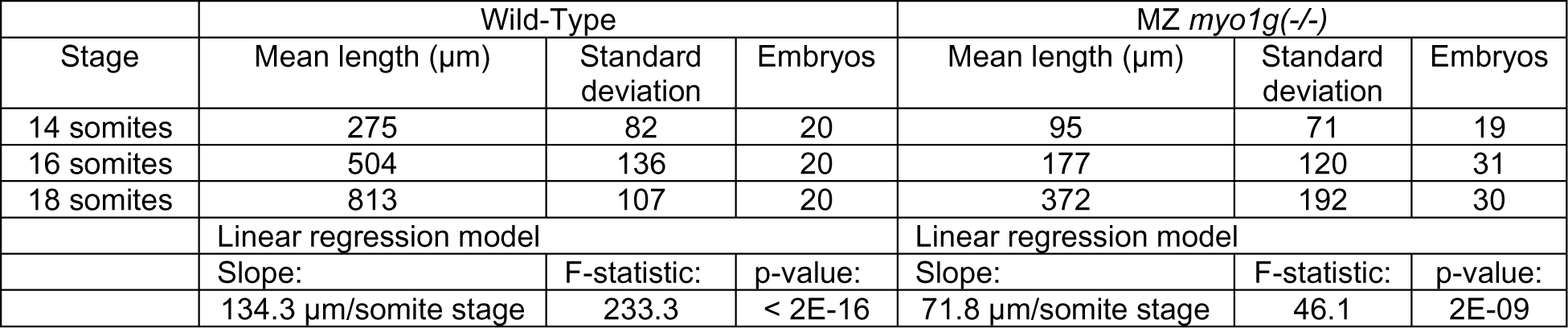
*lefty1* expression propagation in the notochord of *myo1g* mutants.

**Figure 4e:**
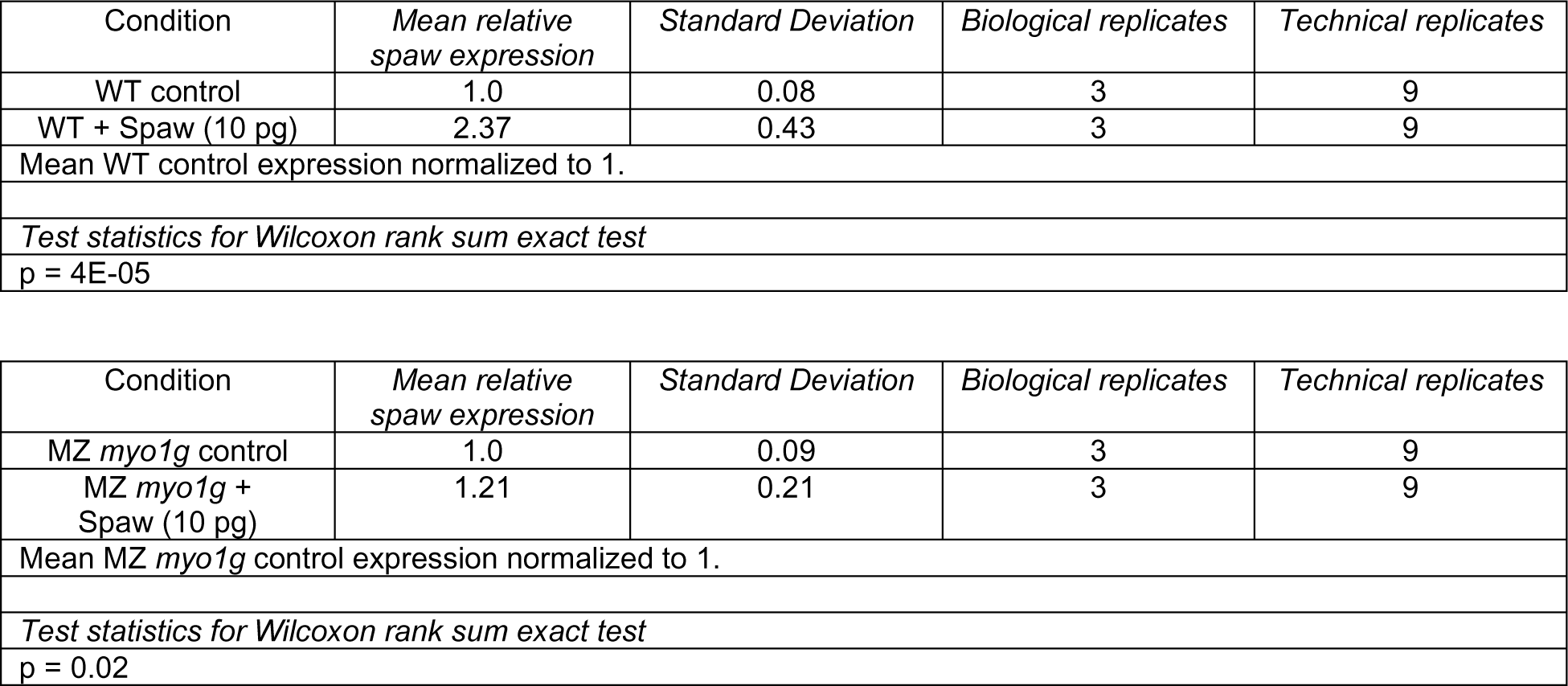
Germ ring stage *lefty1* expression in Spaw-injected *myo1g* mutants (qPCR)

**Figure 4f:**
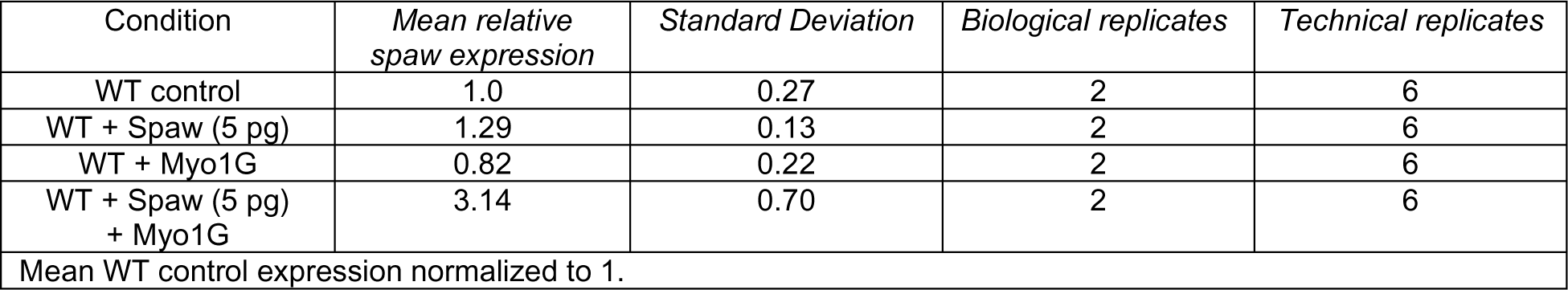
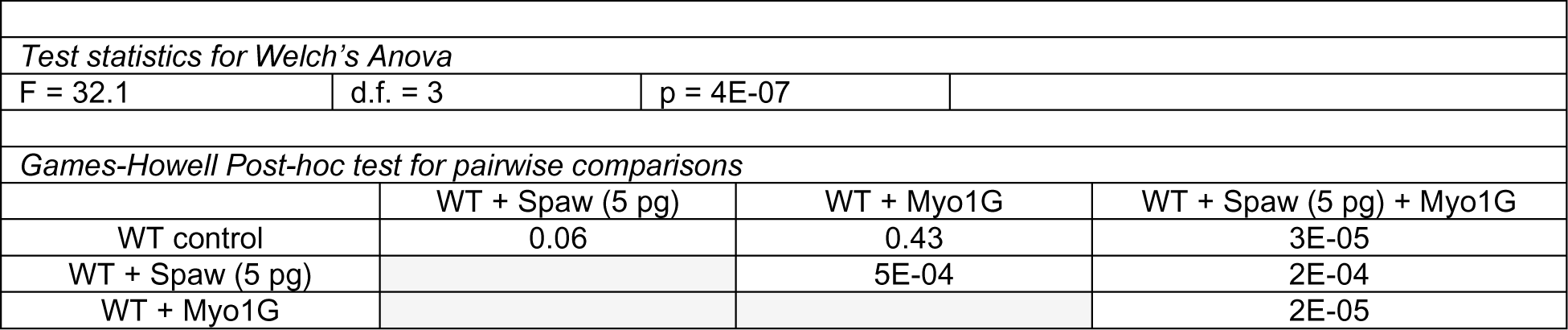
Germ ring stage *lefty1* expression in Spaw + Myo1G injected WT embryos.

**Figure 5b:**
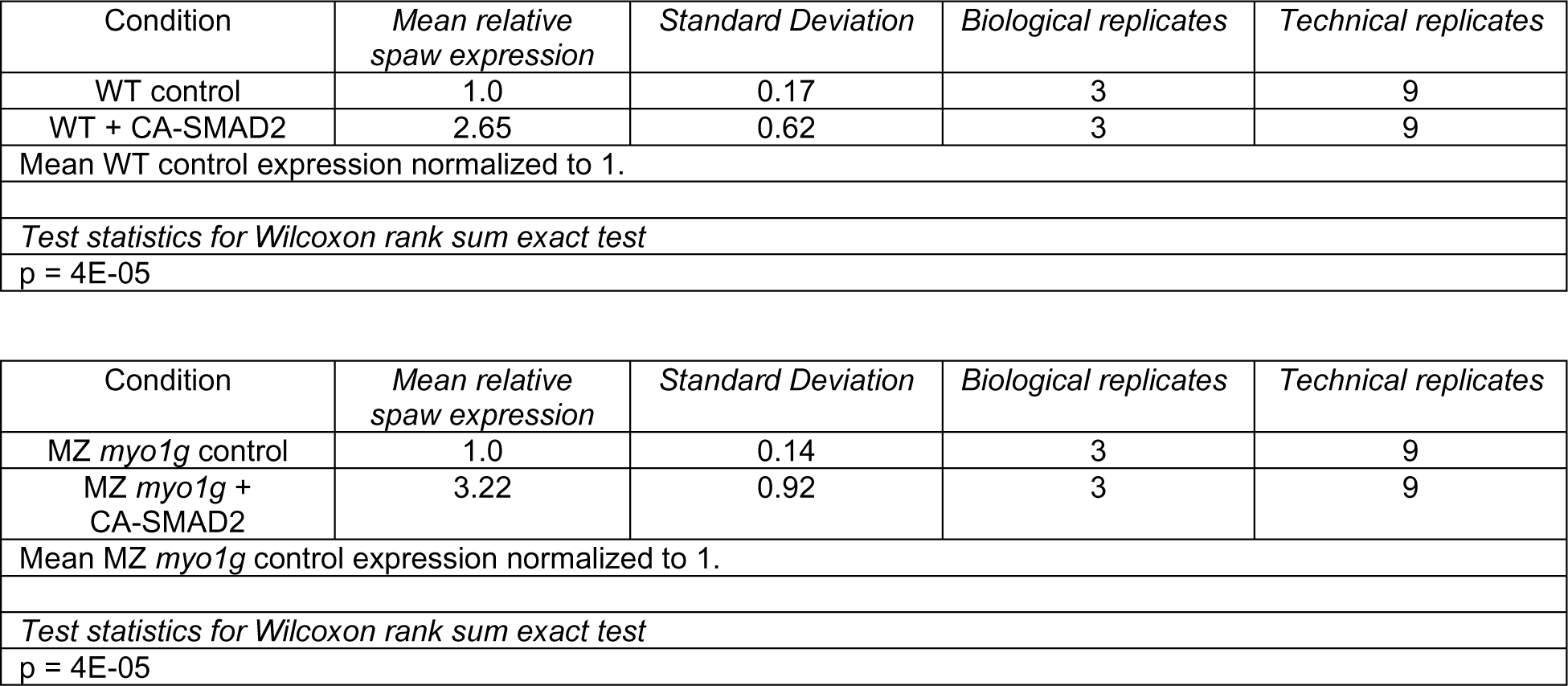
Germ ring stage *lefty1* expression in CA-SMAD2-injected *myo1g* mutants (qPCR)

**Figure 5j:**
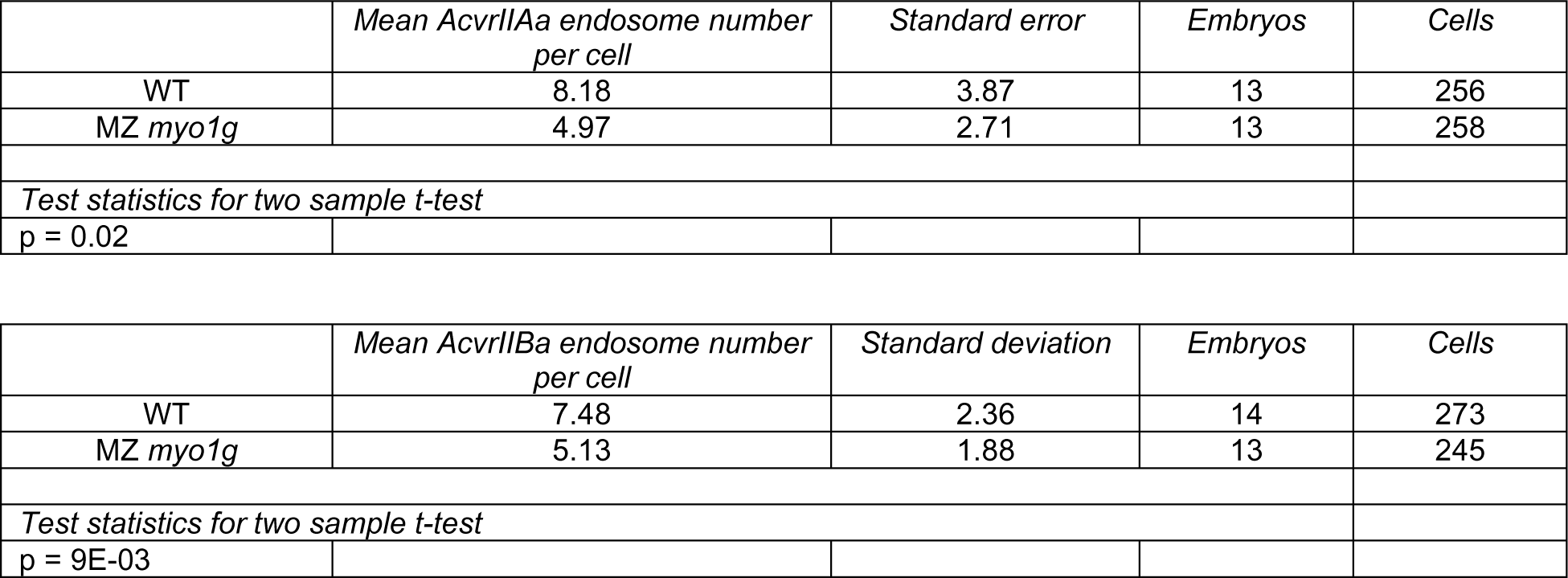
Activin receptor endosome number in *myo1g* mutants.

**Figure 5n:**
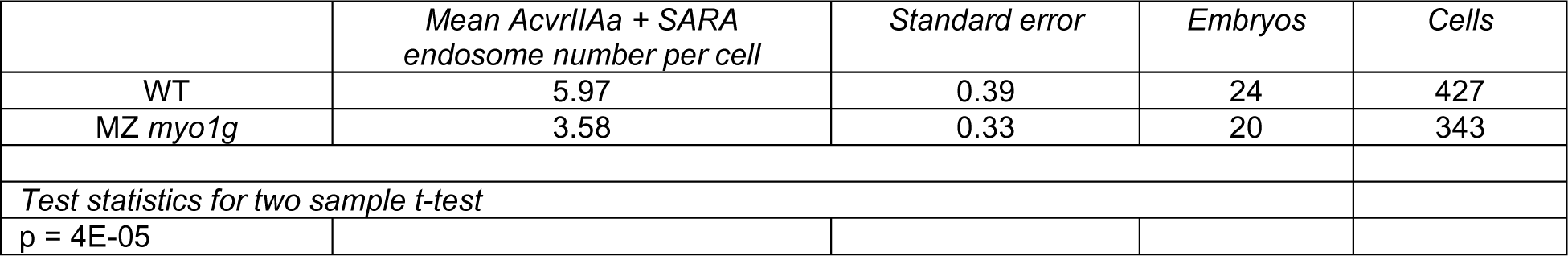
Number of AcvrIIAa + SARA positive endosomes in *myo1g* mutants.

**Figure 5p:**
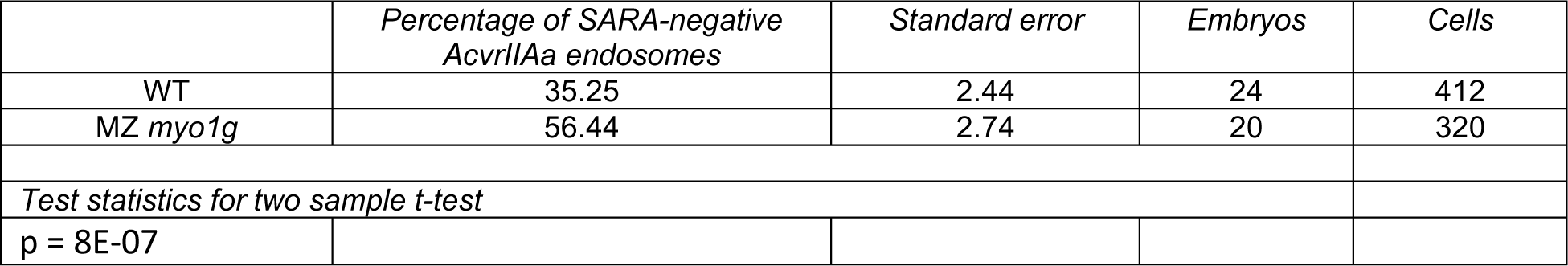
Percentage of SARA-negative AcvrIIAa endosomes in *myo1g* mutants.

**Figure 6a:**
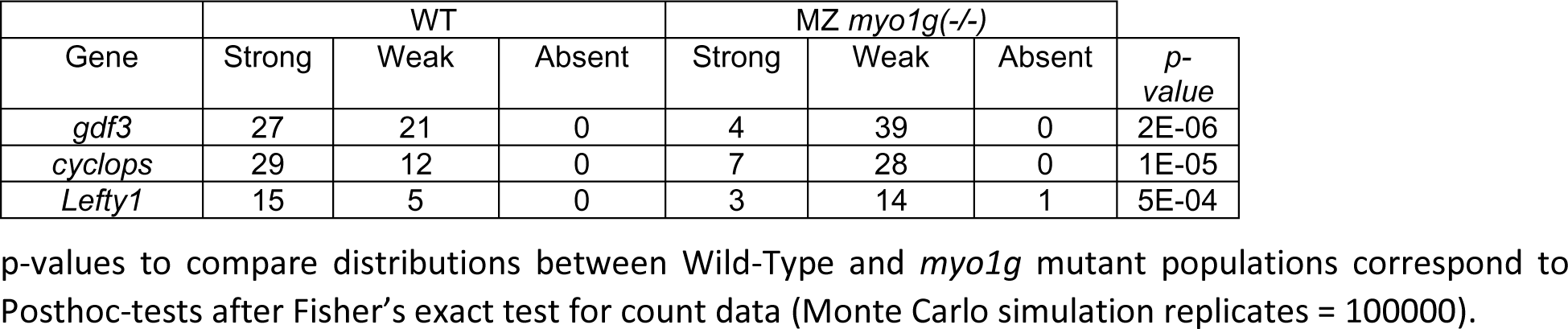
LRO gene expressions in of *myo1g* mutants.

**Figure 6b:**
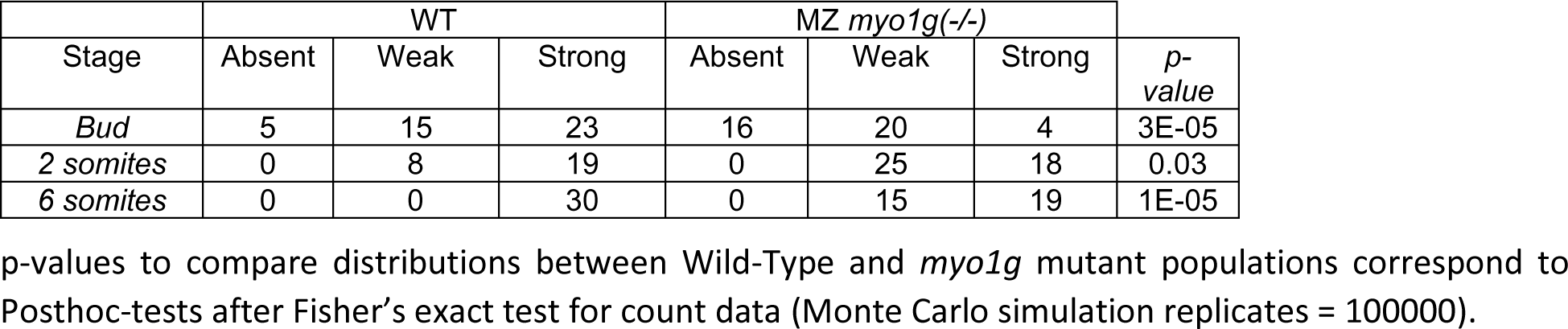
dand5 gene expressions in of *myo1g* mutants.

**Figure 6c:**
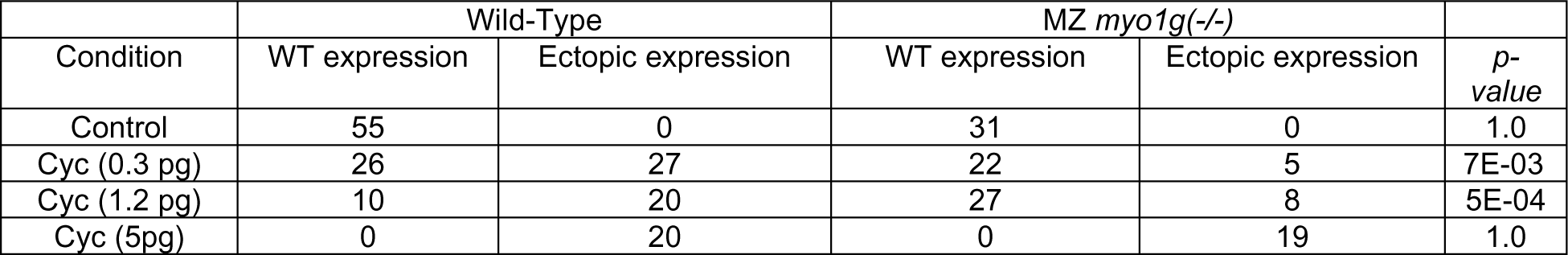
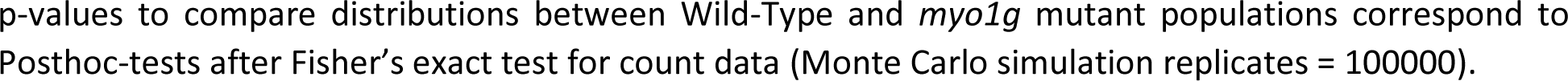
*lefty1* expression in Cyclops-injected *myo1g* mutants.

**Figure 6d:**
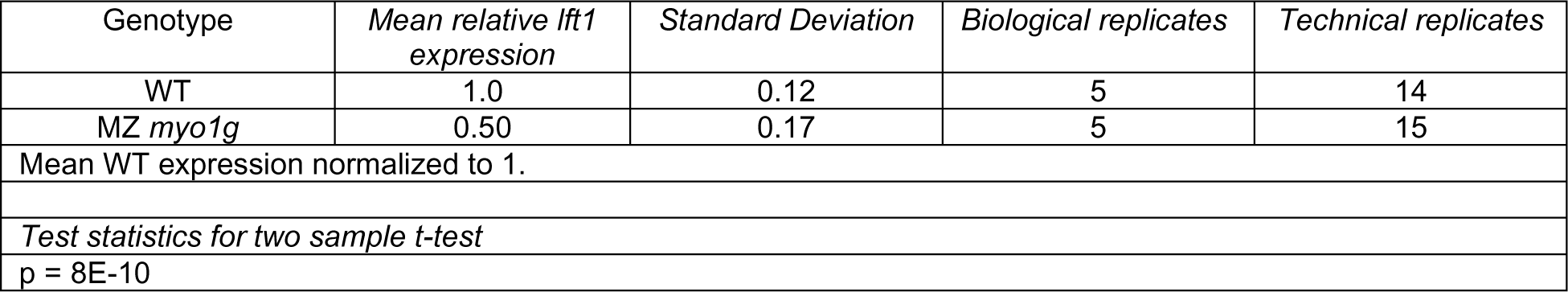
qPCR analysis of germ ring stage *lefty1* expression in *myo1g* mutants.

**Figure 6e:**
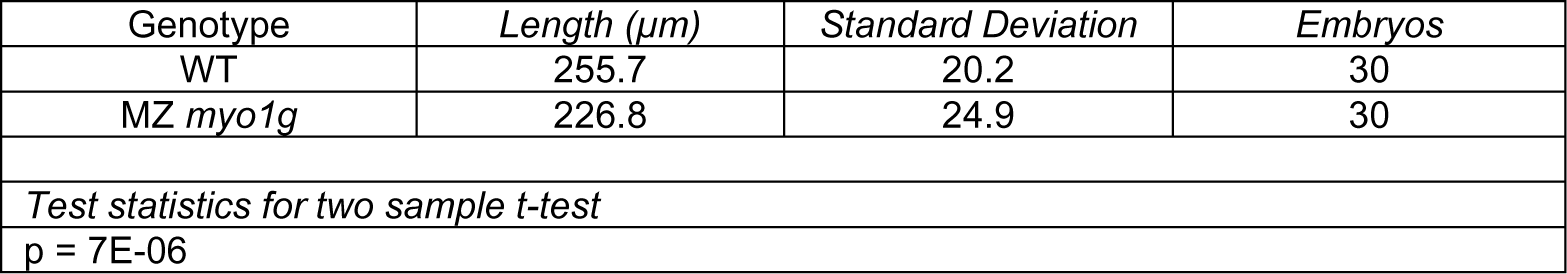
Antero-posterior extension of the 8 somites stage forebrain *lefty1* expression.

**Supplementary Figure 1a:**
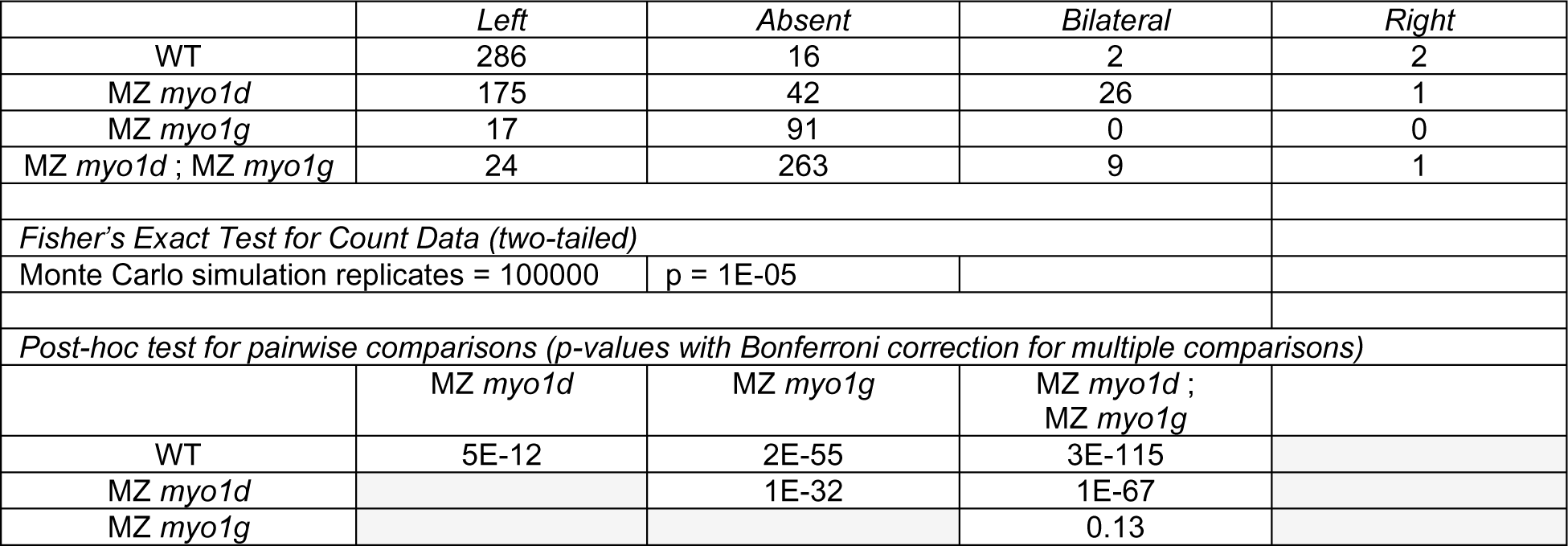
Brain *cyclops* expression in *myosin1* single and double mutants.

**Supplementary Figure 1b:**
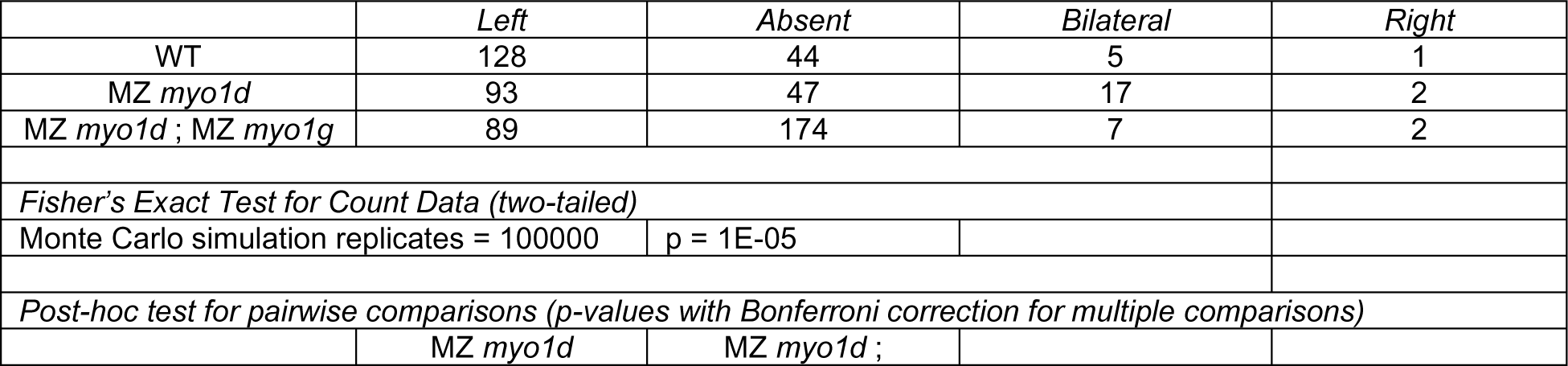
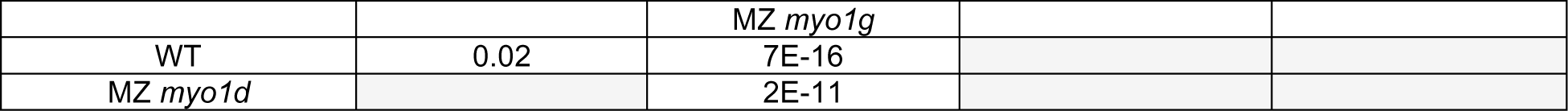
Brain *lefty1* expression in *myosin1* single and double mutants.

**Supplementary Figure 2a:**
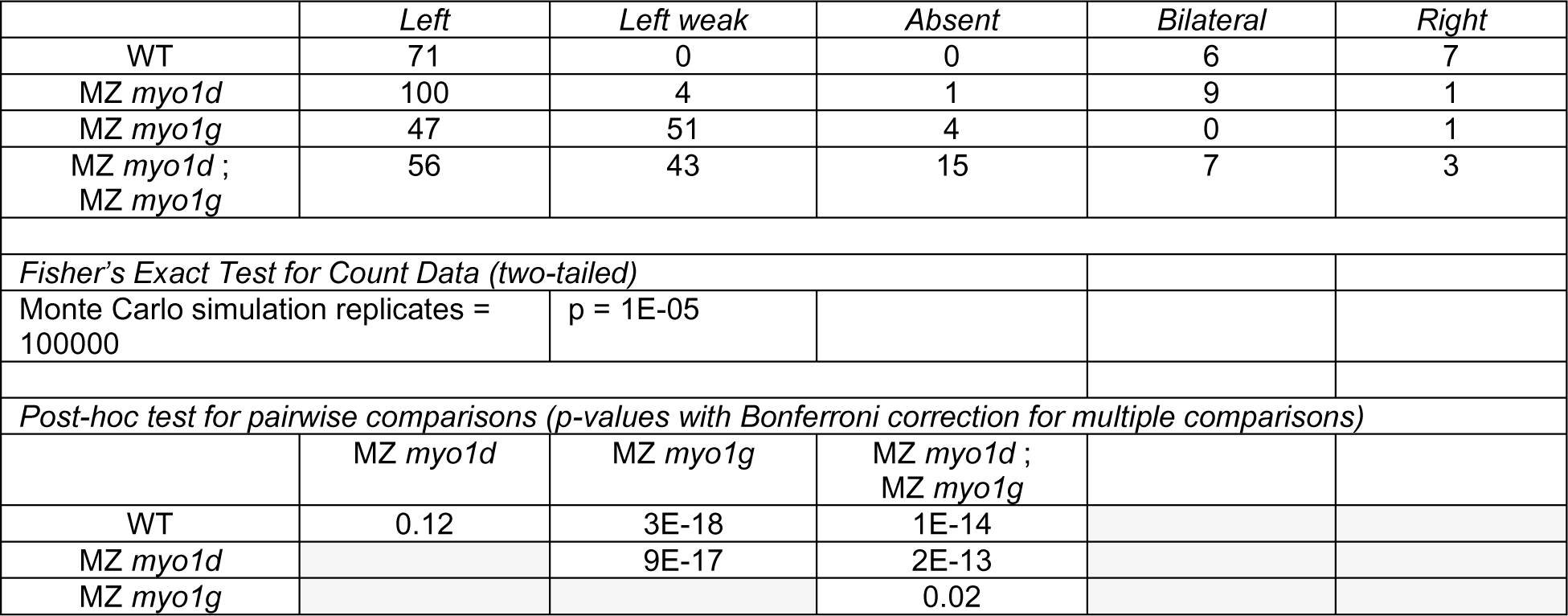
*southpaw* expression in the left lateral plate mesoderm of *myosin1* mutants.

**Supplementary Figure 2b:**
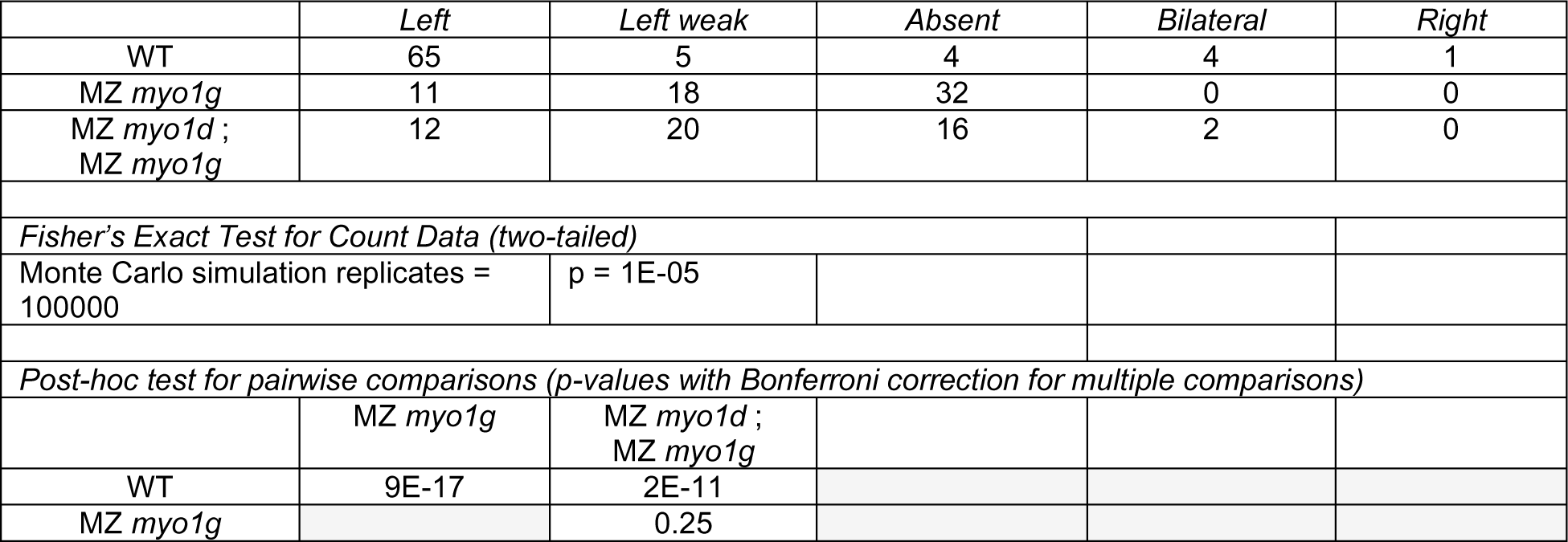
*pitx2* expression in the left lateral plate mesoderm of *myosin1* mutants.

**Supplementary Figure 2c:**
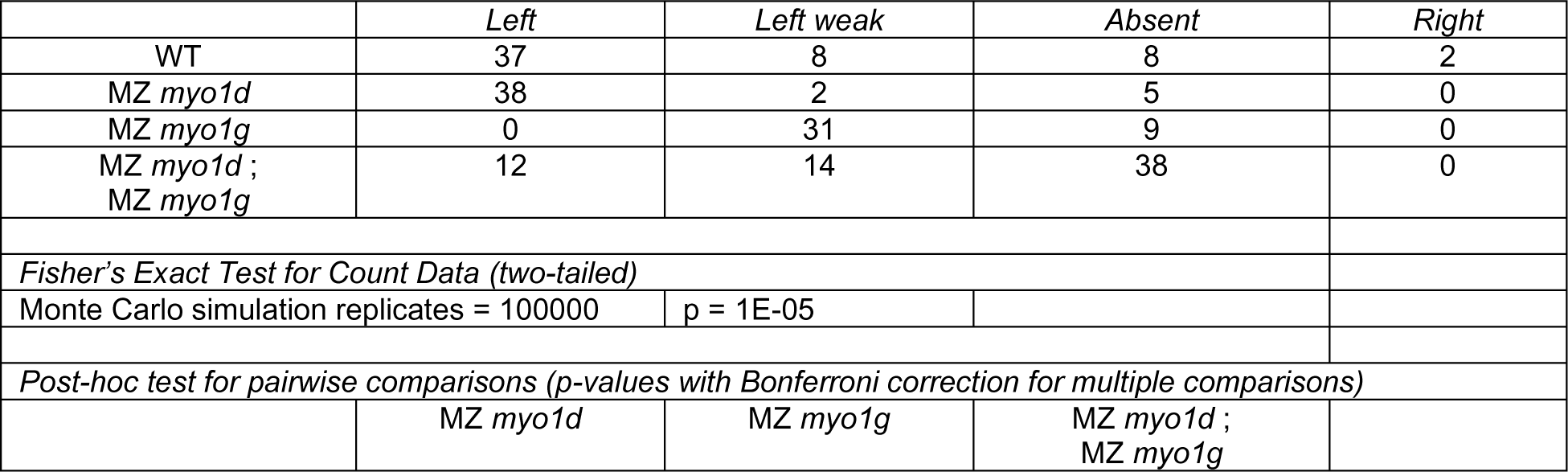
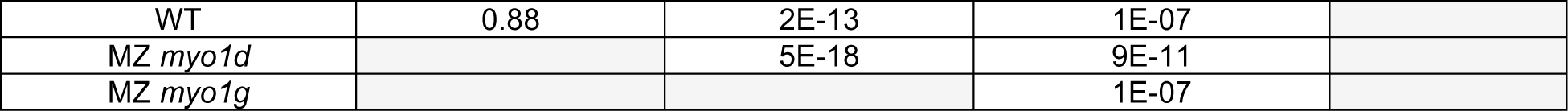
*elovl6* expression in the left lateral plate mesoderm of *myosin1* mutants.

**Supplementary Figure 2d: *Cardiac jogging and looping in Spaw-injected myo1g mutants***

Embryos expressing Spaw on the Left (Fig. 2e) and Right (Supplementary Fig. 2d) were generated in the same series of experiments. Embryo numbers and statistics for this Supplementary Fig. 2d are therefore displayed in the table provided for Fig. 2e.

**Supplementary Figure 3a: *southpaw* propagation to the cardiac primordium in dnaaf1 Morpholino-injected *myo1g* mutants**

Displayed embryos belong to the same data set also displayed in Fig. 3c. Embryo numbers are therefore displayed in the table provided for Fig. 3c.

**Supplementary Figure 3a’: *southpaw* propagation to the cardiac primordium in *dnaaf1 myo1g* single and double mutants**

Displayed embryos belong to the same data set also displayed in Fig. 3c’. Embryo numbers are therefore displayed in the table provided for Fig. 3c’.

**Supplementary Figure 4a:**
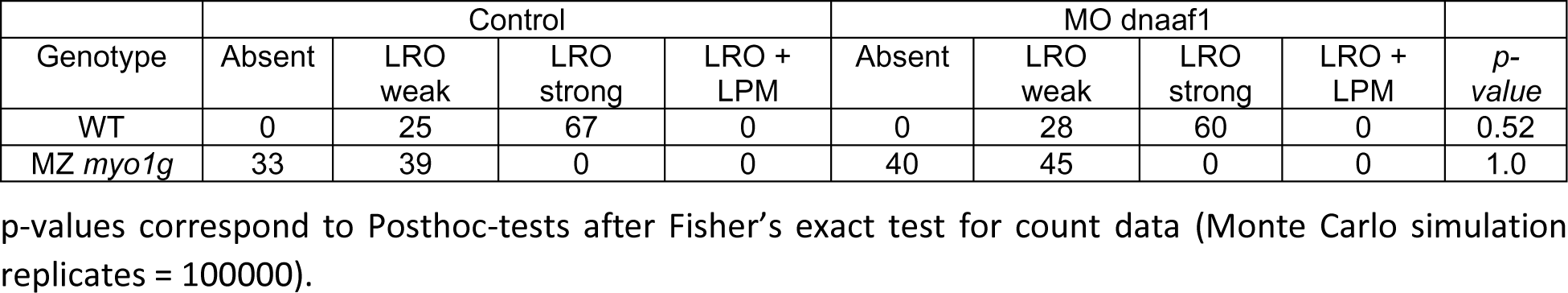
8 somites stage *southpaw* expression in *dnaaf1*-depleted embryos.

**Supplementary Figure 4b:**
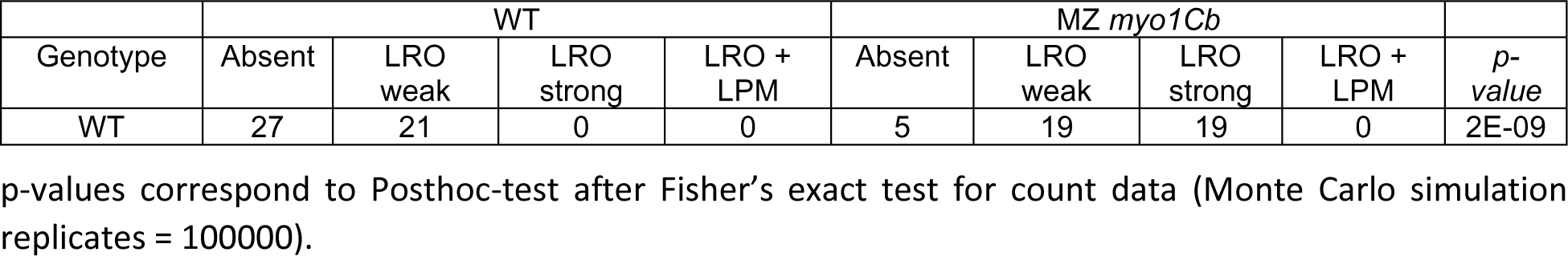
6 somites stage *southpaw* expression in *myo1Cb* mutants.

**Supplementary Figure 6b:**
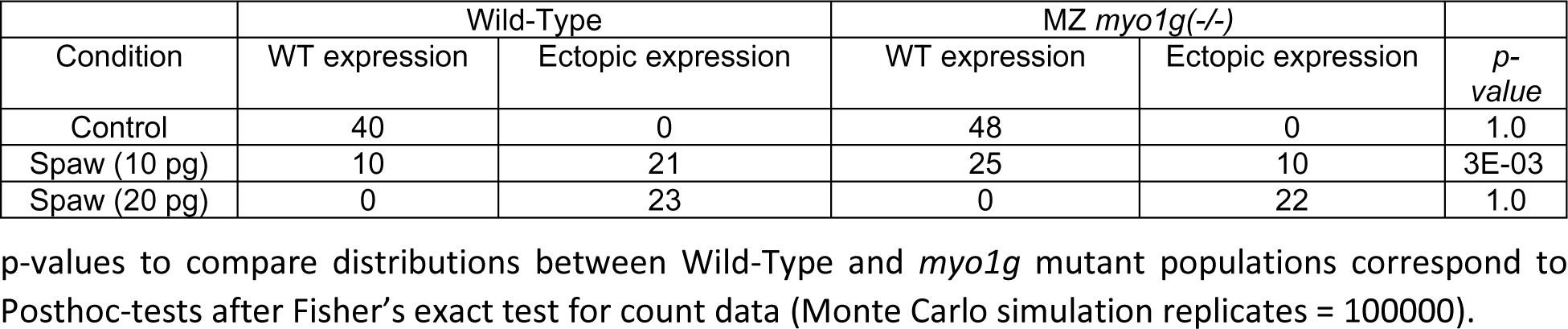
*lefty1* expression in Spaw-injected *myo1g* mutants.

**Supplementary Figure 6c:**
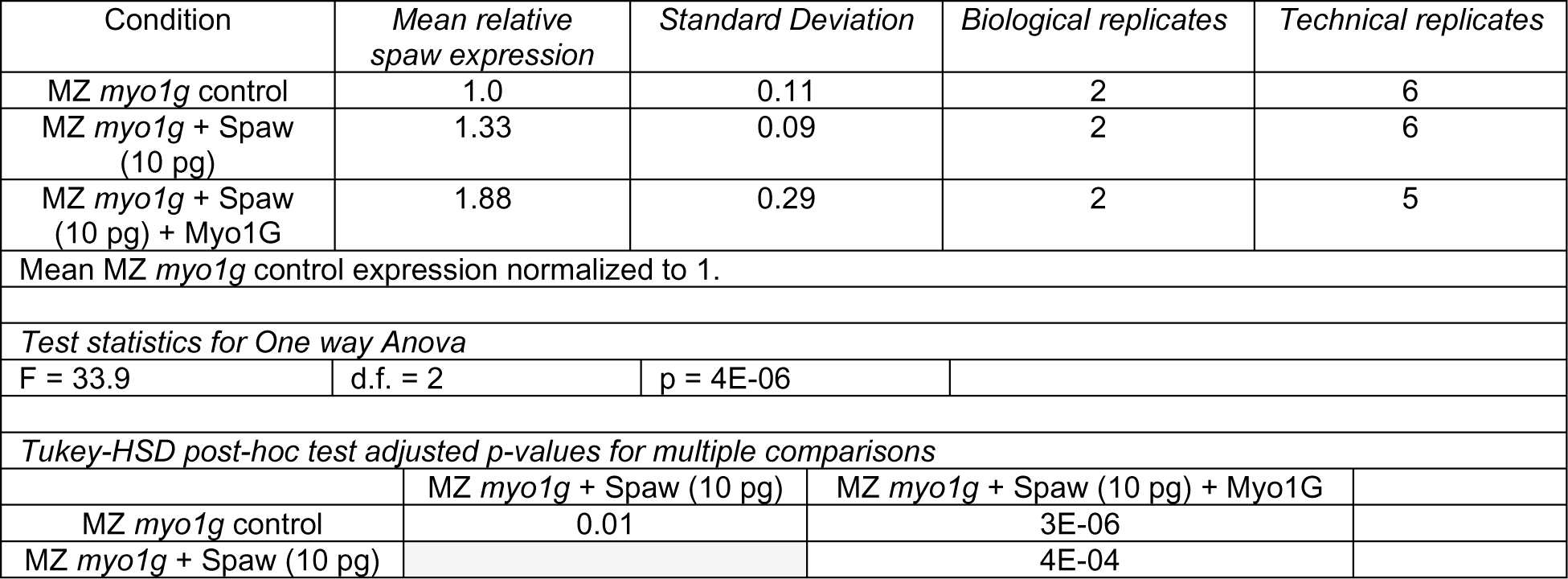
Germ ring stage *lefty1* expression in Spaw + Myo1G injected *myo1g* mutants.

**Supplementary Figure 7c:**
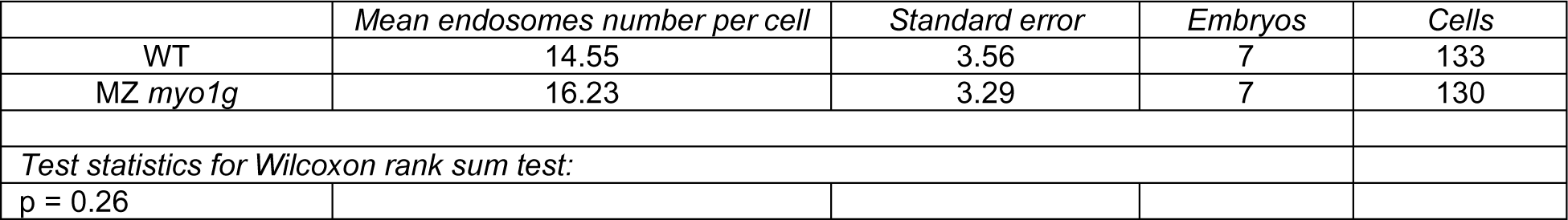
CD44a endosome number in *myo1g* mutants.

**Supplementary Figure 7d:**
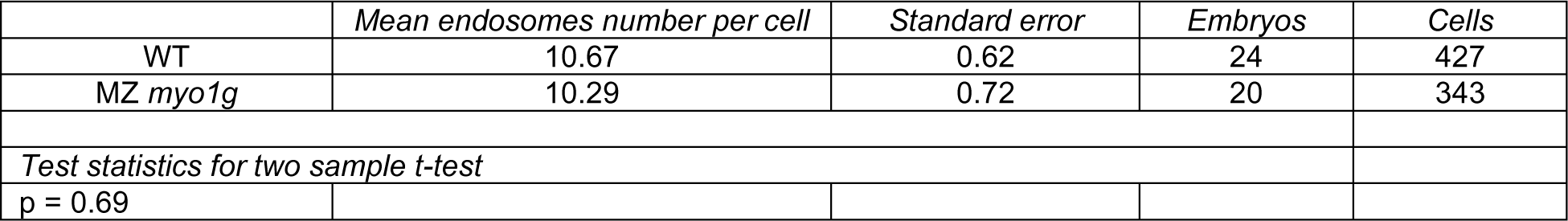
SARA endosome number in *myo1g* mutants.

**Supplementary Figure 8:**
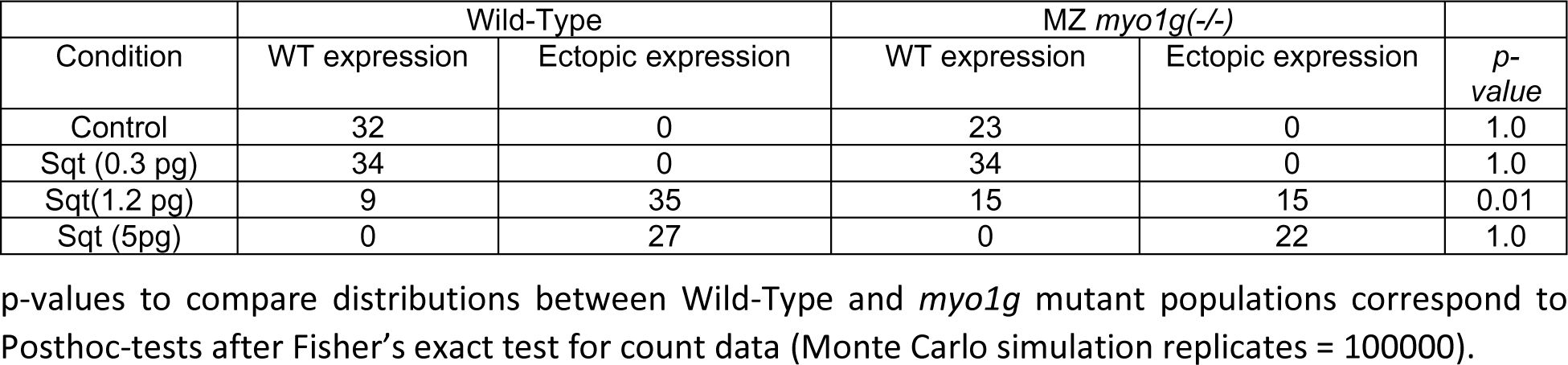
*lefty1* expression in Squint-injected *myo1g* mutants.

## Notes

### Competing Interest Statement

The authors have declared no competing interest.

